# Spartan: activation-aware framework for spatial domain and variable gene discovery

**DOI:** 10.64898/2026.02.18.706570

**Authors:** Mohammad Faiz Iqbal Faiz, Elliot Jokl, Rachel Jennings, Karen Piper Hanley, Andrew Sharrocks, Mudassar Iqbal, Syed Murtuza Baker

## Abstract

Spatial transcriptomics is rapidly advancing toward single-cell-level resolution, revealing complex tissue architectures organized across continuous anatomical gradients. However, accurate identification of spatial domains remains a central computational challenge, as many existing clustering approaches blur anatomical boundaries, merge transitional zones, or fail to resolve localized microstructures. Here we introduce Spartan, an activation-aware multiplex graph framework for high-resolution domain discovery. Spartan integrates spatial topology and Local Spatial Activation (LSA), a neighborhood deviation signal that captures localized transcriptional heterogeneity often attenuated by similarity-based clustering. By jointly modeling cohesion within domains and localized activation structure, Spartan recovers anatomically aligned partitions across spatially resolved transcriptomics technologies including Visium HD, MERFISH, Stereo-seq, and STARmap. We further demonstrate its utility in a high-resolution Visium HD section of developing human esophagus and stomach, where activation-aware graph integration enables precise delineation of complex transitional regions such as the gastroesophageal junction and supports stable multi-scale domain recovery without fragile hyperparameter tuning. Beyond domain identification, Spartan leverages activation-aware structure to detect spatially variable genes associated with localized tissue remodeling. Spartan scales near-linearly with dataset size, providing a robust and interpretable framework for spatial systems-level analysis.

## Introduction

Spatially resolved transcriptomics (SRT) enables gene expression to be measured directly in intact tissue, allowing molecular states to be mapped onto anatomical structures [1, 2, 3]. As SRT technologies increase in resolution and scale, datasets now contain tens of thousands of spatial locations exhibiting fine-grained transcriptional heterogeneity [4]. Accurately resolving spatial organization within such data remains a central computational challenge.

A core task in SRT analysis is spatial domain identification, in which spatial locations are grouped into coherent regions with shared transcriptional programs. Most existing approaches integrate spatial proximity with gene expression similarity to infer such domains, which effectively separates major tissue compartments. However, complex tissues frequently contain localized microenvironments embedded within broader regions, including transitional zones and niche-like structures, where subtle transcriptional shifts define functional specialization. Under these conditions, purely similarity-driven clustering can smooth localized heterogeneity, merge anatomically distinct regions, and overlook fine spatial structure that is biologically meaningful.

To better capture this fine-scale organization, we reasoned that spatial domain discovery should measure not only what each cell or spot expresses and where it resides, but also how its molecular state relates to its immediate neighborhood context. We introduce this additional signal as Local Spatial Activation (LSA), a quantitative measure of neighborhood-relative transcriptional deviation in a low-dimensional latent space. Rather than reinforcing similarity alone, LSA models joint deviation of spatially adjacent spots relative to their shared local transcriptional context. Conceptually, LSA captures localized shifts in molecular programs while preserving spatial coherence. Although related in spirit to local spatial statistics used in computational geography such as local Moran’s I [5, 6], LSA is not a statistical test of spatial autocorrelation. Instead, it is formulated as a non-negative, variance-normalized graph signal designed for integration within a multiplex community detection framework.

Motivated by this principle, we developed Spartan (Spatial Activation-Aware Transcriptomic Analysis Network), a framework for spatial domain recovery that integrates three complementary layers within an aggregated graph: spatial topology, LSA, and gene expression connectivity. Unlike existing spatial clustering approaches that encode spatial proximity and expression similarity within a single homogeneous graph layer, Spartan introduces neighborhood-relative activation as an explicit structural component of the clustering objective. Spartan applies Leiden community detection [7] to this activation-aware graph, enabling efficient and reproducible domain identification while preserving interpretability through explicit layer weighting and transparent parameterization. By incorporating activation as a graph layer, Spartan enables activation-aware clustering while supporting robust scale adaptation across datasets. The aggregated formulation allows Spartan to transition between settings dominated by broad compartmental organization and those requiring sensitivity to fine local heterogeneity.

In high-resolution samples, such as Visium HD sections of the developing human esophagus and stomach, biologically meaningful organization is expressed as thin layers, curved anatomical boundaries, and junctional transition zones, including the gastroesophageal junction (GEJ), where epithelial and stromal programs change gradually rather than forming sharply separated compartments. Resolving such continuous spatial organization illustrates the type of challenge that activation-aware graph integration is designed to address.

We benchmarked Spartan across multiple SRT technologies, including MERFISH, Stereo-seq, STARmap, and osmFISH, spanning diverse resolutions and tissue architectures. Spartan achieved strong and consistent performance across datasets, frequently matching or exceeding recently proposed methods including BANKSY [8], NichePCA [9], and SpatialLeiden [10]. Ablation analyses confirmed the complementary contribution of each graph component, with the multiplex configuration providing the most robust domain recovery across technologies and experimental settings. Beyond spatial domain identification, Spartan extends naturally to spatially variable gene detection by leveraging LSA to identify genes associated with localized spatial structure, providing a scalable and extensible framework for spatial transcriptomics and future spatial multi-omics integration.

## Results

### Overview of Spartan

Spartan is a computational framework for the accurate identification of spatial domains and detection of spatially variable genes (SVGs) across diverse SRT platforms. It integrates spatial topology and gene expression structure through three biologically grounded graphs that jointly encode spatial proximity, LSA (conceptual illustration in Fig. 1A), and expression-derived connectivity.

**Figure 1.**
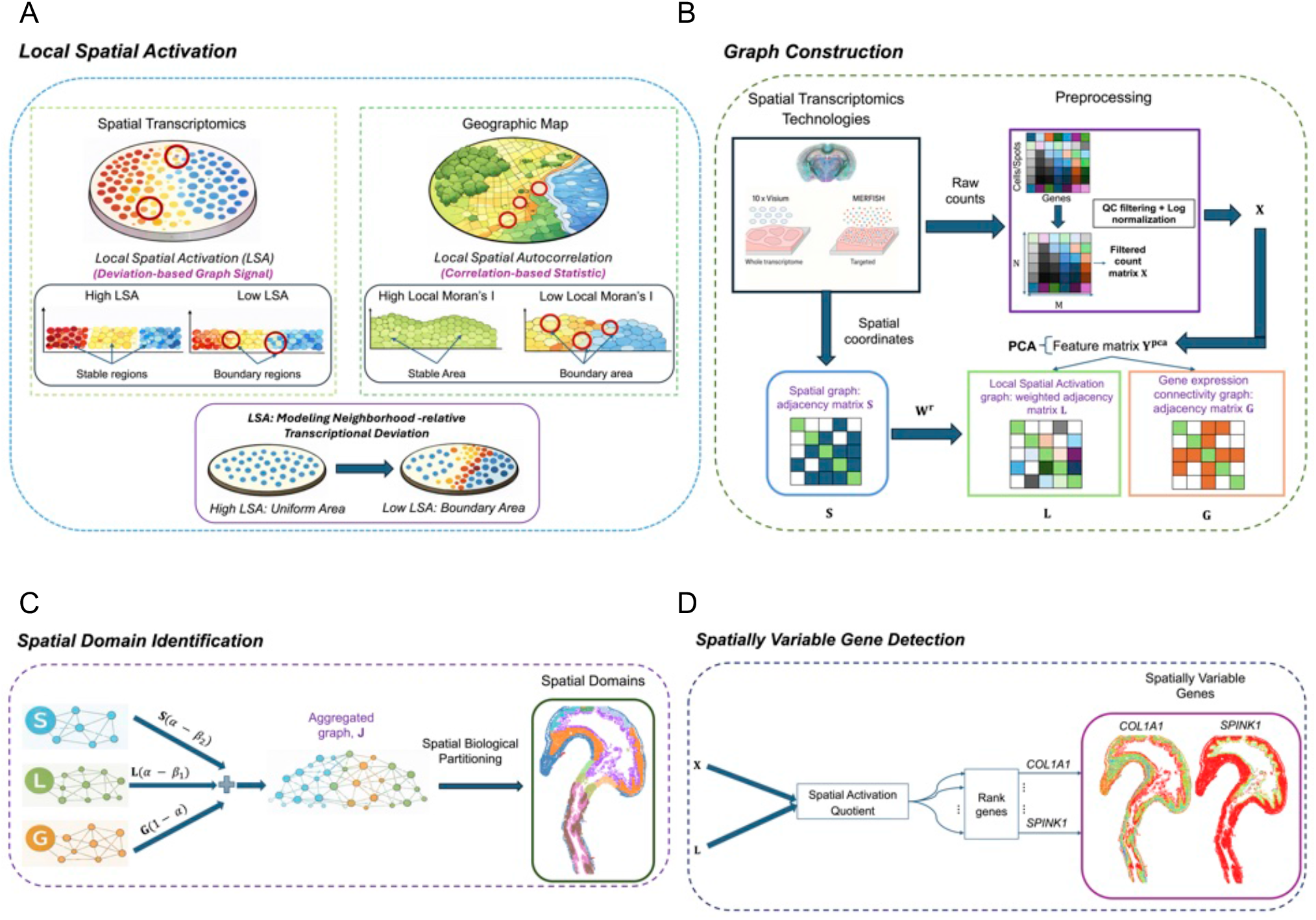
Overview of the Spartan framework for spatial domain identification and spatially variable gene detection. (A) Conceptual comparison between Local Spatial Activation (LSA) in spatial transcriptomics and classical local spatial autocorrelation (Local Moran’s I) in geostatistics. Left: In spatial transcriptomics, LSA models neighborhood-relative transcriptional deviation, capturing localized shifts in molecular programs within tissue architecture. High LSA values occur in spatially coherent regions, with lower values at boundaries exhibiting structured transcriptional variation. Right: In geostatistics, Local Moran’s I (LISA) measures local spatial autocorrelation, identifying regions where neighboring locations exhibit similar values. Boundary regions exhibit reduced autocorrelation in heterogeneous areas and elevated autocorrelation in spatially coherent regions. Unlike classical statistical autocorrelation measures, LSA is formulated as a constructive graph signal designed for activation-aware community detection in spatial transcriptomics. (B) Graph construction within Spartan. Spartan identifies spatial domains and spatially variable genes (SVGs) through integration of spatial and transcriptomic information. From spatial transcriptomics data, Spartan performs PCA to obtain **Y**^**pca**^ and constructs three complementary graphs: a spatial graph encoding spot proximity, a Local Spatial Activation (LSA) graph capturing fine-scale spatial variation, and a gene expression connectivity graph encoding local transcriptomic neighborhood structure. (C) These graphs are combined into an aggregated graph (represented by weighted adjacency matrix **J**), where contributions from the spatial, gene expression connectivity, and LSA graphs are explicitly aggregated. Leiden clustering on **J** yields biologically coherent spatial domains. (D) The Spatial Activation Quotient (SAQ), derived from the LSA graph, ranks genes by the strength of their spatially structured activation to identify spatially variable genes (SVGs) (green indicates higher enrichment, and red indicates lower enrichment).

From each SRT sample, Spartan extracts spatial coordinates and a preprocessed count matrix **X**, performs principal-component analysis to obtain **Y**^*pca*^, and constructs three complementary graphs (Fig. 1B). The spatial graph, represented by adjacency matrix **S** (Figs. 1B, 2A), encodes physical neighborhood relationships between spots using either k-nearest-neighbor (KNN) or Delaunay connectivity. A central component of Spartan is the LSA graph, represented by **L** (Figs. 1B, 2B). For each pair of neighboring spots *i* and *j*, the weight *l*_*ij*_ combines their attribute deviations *ϕ*_*i*_ and *ϕ*_*j*_ with a spatial proximity term 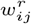. This formulation quantifies LSA, providing an additional spatial signal that captures neighborhood-relative transcriptional deviation beyond expression similarity alone. The gene expression connectivity graph, represented by adjacency matrix **G** (Figs. 1B, 2C), encodes local transcriptomic neighborhood structure in PCA-reduced space.

**Figure 2.**
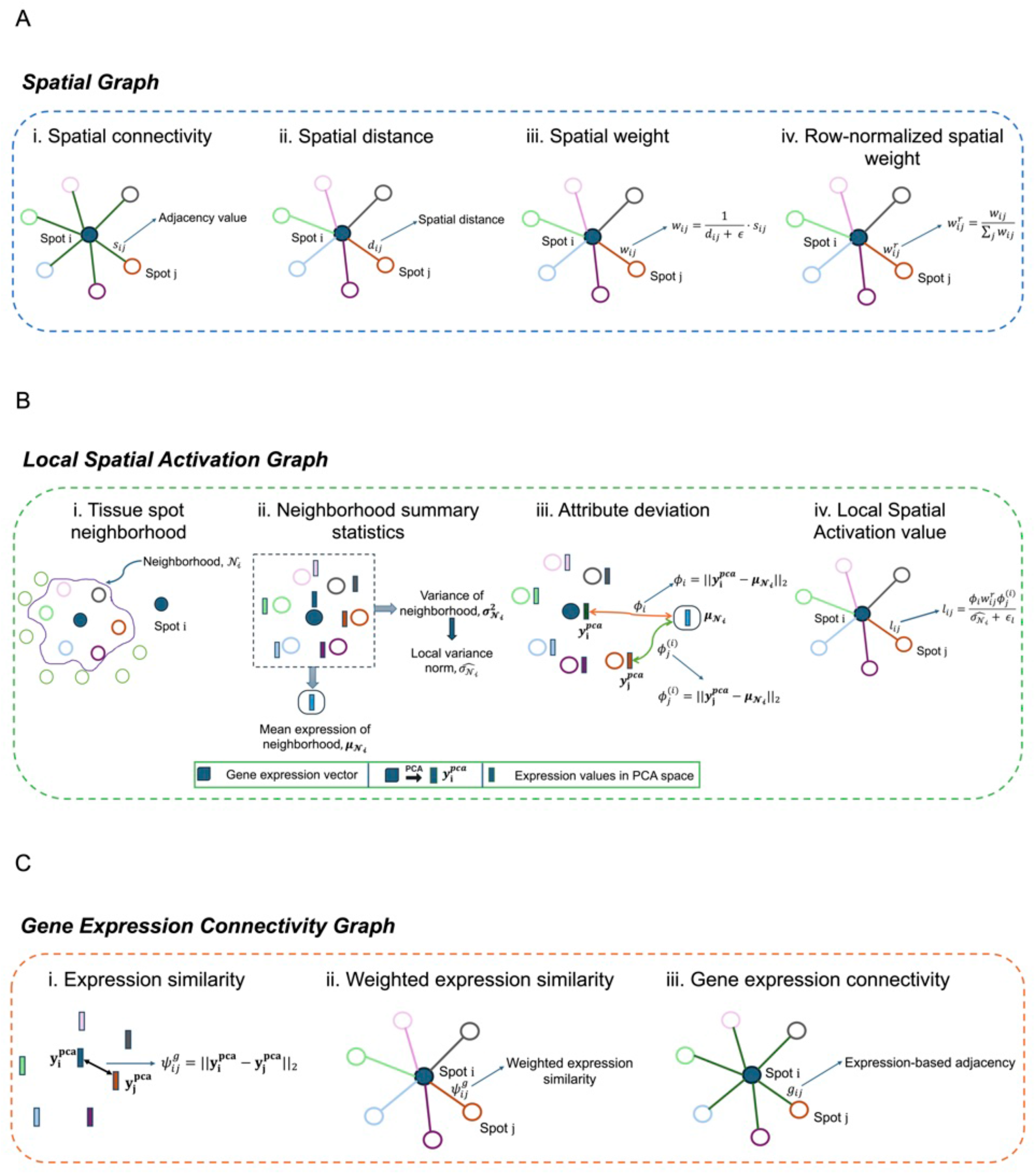
Overview of Spartan’s graph construction. (A) Spartan constructs a spatial graph represented by the adjacency matrix **S**. The spatial weight *w*_*ij*_ for each edge is computed from the Euclidean distance *d*_*ij*_ between spot *i* and spot *j*, together with the adjacency value *s*_*ij*_. The row-normalized weight 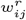 is then obtained from *w*_*ij*_. (B) Spartan constructs the Local Spatial Activation (LSA) graph, represented by the weighted adjacency matrix **L**. Each edge is assigned a weight *l*_*ij*_, which captures the variance-normalized, spatially weighted joint transcriptional deviation of spots *i* and *j* relative to the local neighborhood of spot *i*. (C) Spartan constructs a gene expression connectivity graph represented by the adjacency matrix **G**, where each edge is assigned an adjacency value *g*_*ij*_ based on KNN relationships in the PCA-reduced expression space.

Spartan integrates these three graphs into an aggregated graph, represented by the weighted adjacency matrix **J** (Fig. 1C) that balances spatial proximity, LSA, and expression-derived connectivity. Leiden clustering on **J** partitions the tissue into coherent spatial domains that align with known anatomical regions. To further exploit the LSA signal, Spartan computes a Spatial Activation Quotient (SAQ) (Fig. 1D) for each gene by combining **L** and **X**, ranking genes by their spatial activation strength to identify SVGs.

By modeling spatial relationships at multiple levels, including global proximity (**S**), LSA (**L**), and transcriptomic similarity (**G**), Spartan captures biologically coherent tissue organization and spatially patterned gene programs. Its independence from specific SRT technologies and its ability to detect both structured domains and fine-grained local patterns make Spartan a general and extensible framework for spatial omics analysis.

### Ablation study of multiplex graph components

Next, we conducted an ablation study to evaluate the contribution of each component in the aggregated graph *gr*^mul^. The aggregated graph comprises three elements (Equation 15): (i) the LSA graph *gr*^lsa^, represented by the weighted adjacency matrix **L**, (ii) the gene expression connectivity graph *gr*^g^, represented by **G**, and (iii) the spatial graph *gr*^s^, represented by **S**. We considered three combinations of these graphs that have both structural and expression-based terms: (i) **L** + **S** + **G**, (ii) **L** + **G**, and (iii) **S** + **G**. For dual-graph formulations (**L** + **G** or **S** + **G**), the combined adjacency matrix was defined as *α* · (**L** or **S**) + (1 − *α*) · **G**. The *β*_1_ and *β*_2_ parameters were only used in **L** + **S** + **G** configurations. Each combination was evaluated using two spatial connectivity schemes, *k*-nearest neighbors (KNN) or Delaunay triangulation, resulting in six total configurations. These configurations were evaluated across four datasets representing different SRT technologies. For multi-sample datasets, the multiplex weighting parameter *α* was selected using a cross-sample hierarchical strategy to avoid sample-specific overfitting (*α*-selection strategy) (conceptual illustration in Supplementary Fig. 1).

We evaluated all configurations across multiple datasets using the R score and ARI score (Fig. 3A-B). Overall, the ablation study supports the aggregated graph formulation implemented in Spartan. Across eight SRT samples from different technologies, including MERFISH and STARmap, the multiplex configuration (**L**+**S**+**G**) performed strongly across all samples. Specifically, the aggregated graph using KNN-based spatial connectivity ranked best in two samples and second best in five, demonstrating strong generalization across datasets. The Delaunay-based configuration also performed competitively, ranking best in one sample, although KNN-based connectivity offers greater stability across all datasets. In contrast, the dual-graph configurations (**L** + **G** or **S** + **G**) exhibited greater variability across samples and technologies. Notably, these dual configurations displayed an inverse relationship: when **L** + **G** achieved high performance, **S** + **G** tended to perform poorly, and vice versa. Together, these findings indicate that the three graph components contribute complementary spatial and transcriptomic information, contributing to consistent clustering performance across diverse SRT datasets.

**Figure 3.**
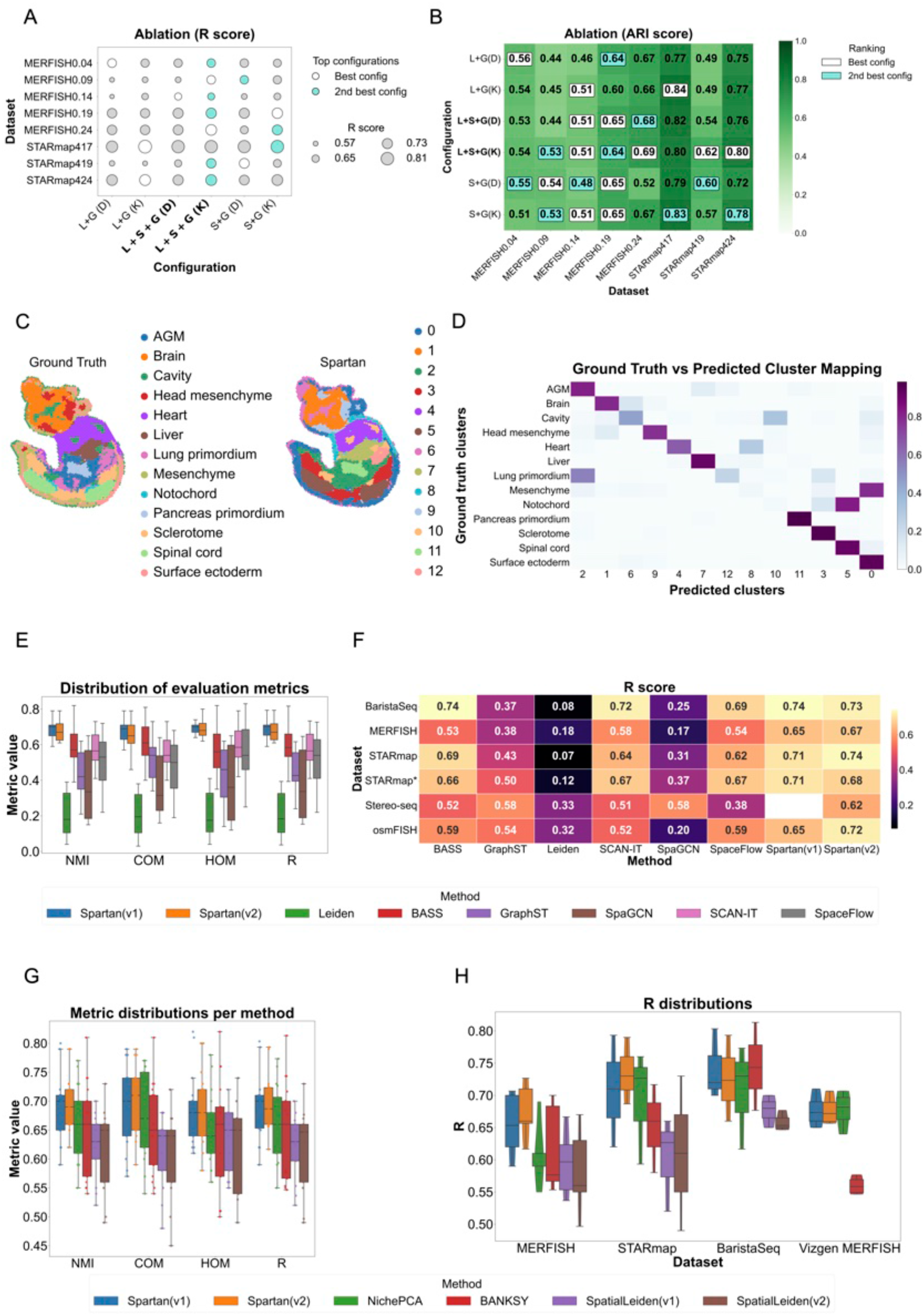
(A–B) Graph ablation. (A) R scores for combinations of LSA (**L**), spatial (**S**), and gene expression connectivity (**G**) graphs using KNN (K) or Delaunay (D) neighborhoods; circle size denotes R, colors indicate best and second-best configurations. (B) Dataset-wise ablation confirms the contribution of each graph component. (C–D) Spatial reconstruction. (C) Example Stereo-seq E9.5E2S2 sample showing ground-truth anatomy (left) and Spartan-derived domains (right). (D) Confusion matrix between predicted clusters and anatomical labels. (E–F) Benchmarking against established methods (SDMBench). (E) Distributions of NMI, COM, HOM, and R using SDMBench-reported results. (F) Dataset-wise R score heatmap. (G–H) Benchmarking against recent methods. (G) Metric distributions for recently published methods evaluated in this study. (H) R score distributions across datasets.

### Validation of Local Spatial Activation (LSA) graph structure

Having established the importance of the multiplex formulation through ablation analysis, we next evaluated whether the Local Spatial Activation (LSA) graph itself captures biologically meaningful spatial organization. We examined the relationship between LSA edge weights and ground truth spatial domains (Supplementary Methods). Because LSA quantifies neighborhood-relative transcriptional activation rather than simple spatial proximity, this analysis tests whether the LSA graph aligns with biologically annotated spatial domains.

First, we compared LSA weights for spatial neighbor edges connecting spots within the same spatial domain and those connecting spots across different domains. Intra-domain edges exhibited systematically higher LSA weights than inter-domain edges, indicating that LSA preferentially strengthens connections within spatial domains (median intra-domain edge weights: 0.444; median inter-domain edge weights: 0.357). The difference between these distributions was highly significant according to a Mann–Whitney U test (*p* = 3.16 × 10^−94^), with a moderate effect size as measured by the rank-biserial correlation (RBC = 0.246) (Supplementary Fig. 2A−C).

To assess whether LSA weights provide discriminative information about spatial domain structure, receiver operating characteristic (ROC) analysis was performed. In the MERFISH 0.24 sample, LSA edge weights demonstrated measurable ability to distinguish intra-domain from inter-domain relationships (AUC = 0.623) (Supplementary Fig. 2D). Next, permutation analysis confirmed that the observed enrichment of LSA weights within spatial domains exceeded that expected under random domain assignments. The observed difference between median intra-domain and inter-domain LSA weights (≈ 0.0869) was substantially larger than values obtained under random permutations (*p* ≈ 0.000999), supporting the conclusion that LSA captures non-random spatial organization in the tissue (Supplementary Fig. 2E).

Similar analysis was done for the Stereo-seq sample E2S3 (Supplementary Fig. 3A-C). Specifically, local LSA magnitudes showed similar discriminative ability for distinguishing boundary from interior regions (AUC = 0.618), consistent with the higher spatial resolution and complex boundary structure characteristic of this technology (Supplementary Fig. 3D).

### Spartan reveals biologically coherent domains in the mouse embryo

The Spartan clustering results on the Stereo-seq E9.5E2S2 sample demonstrate its strong effectiveness in identifying biologically meaningful domains (Fig. 3C). Visually, Spartan generates spatially coherent domains with minimal fragmentation, which aligns well with the tissue morphology. This behavior is consistent with the activation-aware graph construction strategy, which leverages LSA in addition to expression similarity and is well suited to early embryo tissues where spatial gradients and sharp regional boundaries coexist.

The contingency table (ground truth vs. predicted clusters) (Fig. 3D) provides a quantitative view of Spartan’s mapping accuracy. Most tissues, such as the spinal cord, surface ectoderm, heart, and liver, map cleanly to unique predicted clusters, visible as strong diagonal blocks. The predicted clusters broadly recapitulate the major anatomical compartments annotated [11]: forebrain, heart, liver, spinal cord, mesenchyme, and others (Supplementary Fig. 4A). Some tissues, such as mesenchyme and head mesenchyme, exhibit partial overlap across multiple predicted clusters. This does not necessarily indicate misclassification: mesenchyme is known to be heterogeneous and spread across numerous anatomical regions in E9.5 embryos [11], so splitting into multiple clusters may reflect biological heterogeneity or annotation granularity rather than misclassifications. Thus, Spartan not only recovers the ground truth domains but also reveals finer granularity potentially providing new biological insights.

Overall, these results highlight Spartan’s effectiveness not just in matching ground truth but in extending beyond it by resolving complex tissues into finer subdomains.

### Spartan demonstrates competitive and statistically supported performance on public SRT datasets

The benchmarking study was divided into two parts. First, Spartan was benchmarked against representative and widely used methods for spatial domain identification using datasets from the SDMBench suite. Second, Spartan was compared against recently proposed methods using multiple datasets, including those from SDMBench as well as the Vizgen MERFISH dataset. Two versions of Spartan were evaluated: Spartan (v1), which employs a Delaunay graph to represent neighborhood connectivity in spatial coordinate space, and Spartan (v2), which employs a KNN graph to represent neighborhood relations.

#### Benchmarking on SDMBench

Spartan was benchmarked on six datasets from SDMBench: STARmap, STARmap*, MERFISH, BaristaSeq, osmFISH, and Stereo-seq. For competing methods covered by SDMBench, we report the published benchmark results generated under the centralized SDMBench framework using matched cluster counts and standardized evaluation metrics while respecting method-specific preprocessing and architectural settings. Performance was assessed in comparison with five representative and widely used methods: BASS [12], GraphST [13], SpaGCN [14], SCANIT [15] and SpaceFlow [16], as well as the standard Leiden clustering algorithm.

The distribution of evaluation metrics (NMI, COM, HOM, and R) across datasets and methods is illustrated (Fig. 3E; Supplementary Figs. 5A, 6A, 7A). Both versions of Spartan attain the highest or near-highest R scores in five of six SDMBench datasets and rank within the top-performing methods across all datasets. In MERFISH, Spartan improves median NMI by approximately +0.05 and HOM by +0.08 relative to the next-best method under the SDMBench protocol. In Stereo-seq, median improvements reach approximately +0.03 in NMI and +0.05 in HOM. In STARmap, STARmap*, and osmFISH, Spartan achieves the highest or near-highest R scores under the centralized SDMBench protocol.

These gains are observed across both sequencing-based and imaging-based platforms under the SDMBench evaluation protocol. The relatively small interquartile ranges indicate consistent performance across datasets, consistent with the stability of the multiplex integration of LSA, neighborhood connectivity, and gene expression connectivity.

The R score heatmap across individual datasets (Fig. 3F; Supplementary Figs. 5B, 6B, 7B) further supports this observation. Spartan consistently achieves high or near-high performance, with particularly strong gains in MERFISH and Stereo-seq.

An aggregated radar plot (Supplementary Fig. 4B) highlights trade-offs across evaluation metrics. Both versions of Spartan display a balanced performance profile across COM and HOM, whereas competing methods often exhibit stronger trade-offs between completeness and homogeneity.

To evaluate whether the observed performance differences reflect systematic cross-sample improvements rather than sampling variability, we performed paired two-sided Wilcoxon signed-rank tests comparing both versions of Spartan (v1 and v2) against each representative method across samples within MERFISH (n=5) and Stereo-seq (n=9). All statistical tests were conducted on per-sample metric values rather than aggregated dataset summaries (not median summaries). P-values were adjusted using the Benjamini–Hochberg procedure within each dataset and metric. Median paired differences and adjusted P-values are summarized in Supplementary Fig. 8.

Across Stereo-seq (n=9), Spartan (v2) demonstrates statistically significant paired improvements relative to several competitors after FDR correction across NMI, HOM, and COM. In MERFISH (n=5), median paired differences remain consistently positive but do not uniformly reach statistical significance after correction, consistent with the smaller number of paired samples. These findings indicate that Spartan’s performance gains are reproducible across independent samples and are not attributable to isolated dataset-specific instances.

#### Benchmarking against recently proposed methods

Three recently published methods for identifying spatial domains were implemented. These included SpatialLeiden [10], NichePCA [9], and BANKSY [8]. Two versions of SpatialLeiden were evaluated: SpatialLeiden (v1), which uses PCA for feature selection, and SpatialLeiden (v2), which employs MULTISPATI PCA [17]. The KNN graph was used to encode the neighborhood connectivity of cells in both versions of SpatialLeiden, consistent with their reported benchmarking configuration.

Five datasets from the SDMBench (MERFISH, STARmap, STARmap*, BaristaSeq, osmFISH) were used to benchmark Spartan against these methods. In addition, the Vizgen MERFISH dataset was employed to assess Spartan’s scalability and its ability to detect spatial domains in high-resolution samples.

To verify faithful baseline implementation, we quantitatively compared our reproduced results against the originally reported values. For SpatialLeiden (v1 and v2 configurations), reproduced median NMI values across MERFISH, STARmap, osmFISH, and BaristaSeq deviated from the originally reported medians by typically 0.01–0.04 (mean absolute deviation ≈0.03), with a single larger deviation of 0.086 in one MERFISH sample. For NichePCA and BANKSY, reproduced median NMI values closely matched published results across Vizgen MERFISH, MERFISH, STARmap, and BaristaSeq (absolute deviation ≤0.04 in all but one case, where BANKSY improved by +0.08 under our implementation). These deviations confirm faithful reproduction of baseline pipelines and indicate that implementation differences are unlikely to account for comparative performance differences.

In several datasets, methods such as NichePCA, BANKSY, and SpatialLeiden exhibit wider spreads and greater variability across datasets. Importantly, Spartan performs well on both COM and HOM simultaneously, whereas competing methods often trade one for the other (Fig. 3G; Supplementary Figs. 9A, 10A, 11A, 12A). This balanced performance is consistent with Spartan’s multiplex design, which integrates LSA and neighborhood structure to stabilize performance against dataset-specific challenges.

The R-score distributions across datasets are further examined and both versions of Spartan consistently rank among the top performers, with the strongest gains observed in MERFISH and STARmap (Fig. 3H; Supplementary Figs. 9B-C, 10B-C, 11B-C, 12B-C). These datasets are characterized by fine-grained spatial heterogeneity between neighboring cells and non-uniform cell density. Spartan’s incorporation of the LSA component plausibly contributes to improved sensitivity to fine-scale differences by considering attribute deviations of spots or cells relative to their neighborhood.

The ARI scores of Spartan (v1 and v2), NichePCA, SpatialLeiden (v1 and v2), and BANKSY across five datasets are shown in (Fig. 4A-B; Supplementary Fig. 13A-D). Regardless of version, Spartan attains the highest ARI scores in three of five datasets (MERFISH, STARmap, and STARmap*) and ranks second in the remaining two (osmFISH and BaristaSeq).

**Figure 4.**
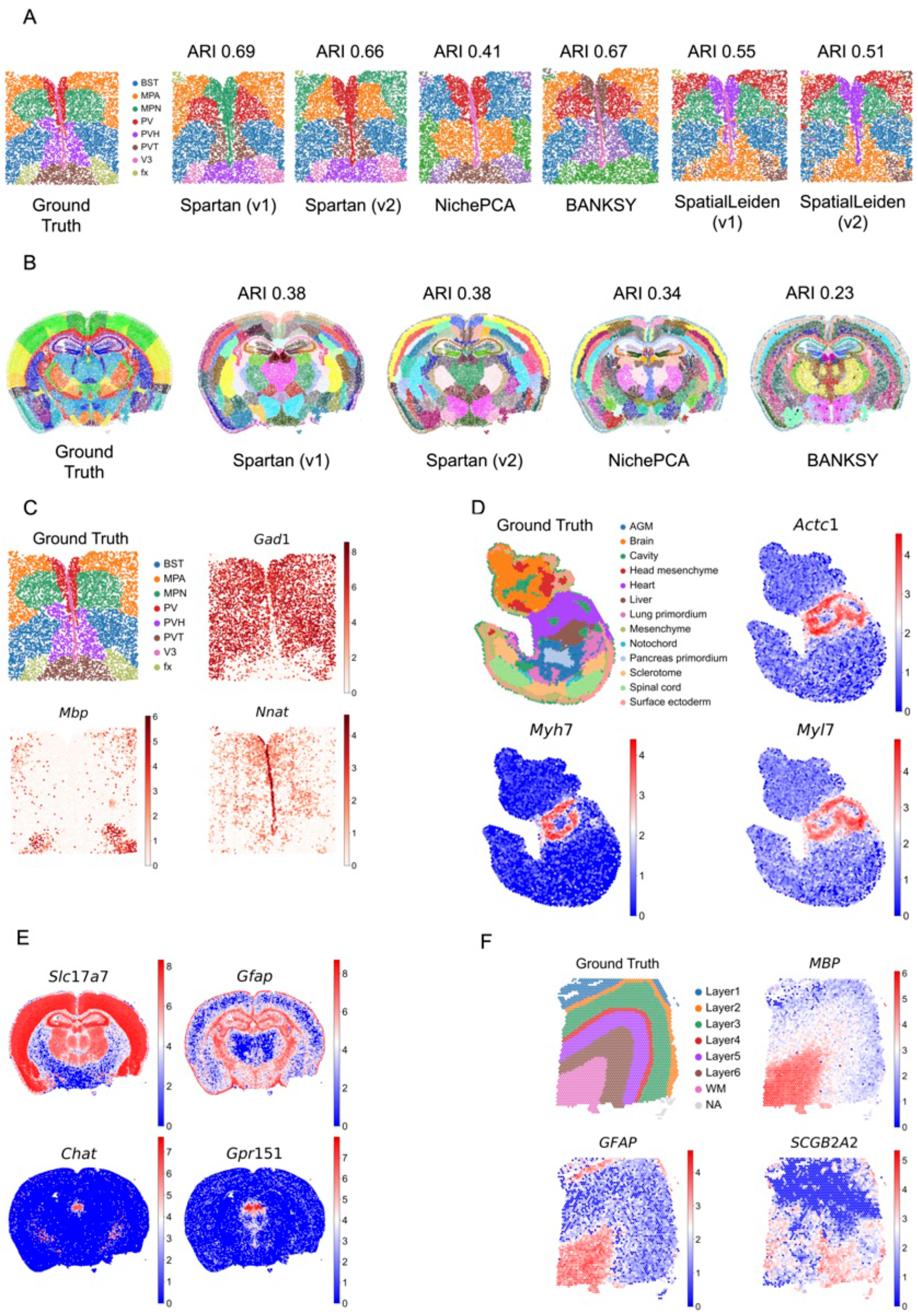
(A–B) Spatial domain identification for MERFISH 0.19 (SDMBench) and Vizgen MERFISH S2R3 samples. Ground-truth annotations (left) are compared with clustering results from Spartan (v1, v2), NichePCA, BANKSY, and SpatialLeiden (v1, v2). ARI values above each map quantify agreement with ground truth. (C) Spatially variable gene (SVG) examples for the MERFISH 0.19 sample. Spatial expression patterns align with anatomical boundaries and support the cluster assignments in (A). (D) Stereo-seq mouse embryo (E9.5E2S2). Ground-truth annotations (left) and spatial expression of representative embryo marker genes (right) show anatomically consistent localization. (E) SVGs for the Vizgen MERFISH mouse brain S2R3 sample, highlighting region- and cell-type–specific expression patterns. (F) 10x Visium human dorsolateral prefrontal cortex (DLPFC) sample 151673. Ground-truth cortical layers (left) and layer-enriched marker genes (right) demonstrate accurate laminar recovery.

To quantify cross-sample robustness relative to recently proposed methods, we performed paired two-sided Wilcoxon signed-rank tests comparing Spartan (v1 and v2) against NichePCA and BANKSY across MERFISH and Vizgen MERFISH samples, and against both SpatialLeiden configurations across MERFISH samples. P-values were adjusted using the Benjamini–Hochberg procedure within each dataset and metric. Median paired differences and adjusted P-values are summarized in Supplementary Fig. 14.

Across MERFISH, Spartan (v2) exhibits consistently positive median paired differences across NMI, HOM, and COM relative to competing methods, with several comparisons remaining significant after FDR correction. In Vizgen MERFISH, improvements are directionally consistent, while statistical significance varies across metrics, reflecting the limited number of paired samples. No systematic degradations relative to competing methods were observed. Together, these results indicate that Spartan’s performance remains stable across independent samples and is not driven by isolated dataset-specific effects.

Taken together, across both SDMBench and recently proposed methods, Spartan demonstrates stable cross-sample performance and competitive results with statistically supported improvements across multiple technologies in several datasets. The paired statistical analyses confirm that observed gains reflect systematic effects rather than isolated dataset-specific instances. Notably, Spartan (v2), which employs a fixed-degree KNN spatial graph, exhibits the most stable and reproducible improvements across metrics and datasets, suggesting that controlled neighborhood regularization enhances multiplex activation-aware integration in heterogeneous spatial contexts. Under the evaluation protocols applied here, these results suggest that Spartan provides a robust and generalizable framework for spatial domain identification across diverse spatial transcriptomics technologies.

### Spartan identifies cell-specific, layer-specific, and region-specific SVGs

In the preoptic and hypothalamic regions of the adult mouse brain sample, obtained from the MERFISH technology, Spartan effectively identifies spatially variable genes (SVGs) that delineate distinct anatomical and cellular domains (Fig. 4C; Supplementary Fig. 15A). The inhibitory neuron marker *Gad1* is detected across nearly all gray matter structures, including the bed nuclei of the stria terminalis (BST), medial preoptic area (MPA), medial preoptic nucleus (MPN), periventricular hypothalamic nucleus (PV), and paraventricular hypothalamic nucleus (PVH), but is absent in the fornix (fx) and paraventricular thalamic nucleus (PVT). This pattern is consistent with the known distribution of GABAergic neurons [18, 19], as the fx and PVT are primarily white matter and glutamatergic, respectively. Conversely, *Mbp* is strongly expressed only in the fx, highlighting oligodendrocyte-rich white matter tracts [20], whereas *Nnat* is restricted to the V3 ventricular zone, marking ependymal cells [21]. The calcium-regulatory gene *Sln* is present in most neuronal and ependymal regions [22] but absent in fx, and the neuropeptide gene *Oxt* is highly specific to the PVH, corresponding to magnocellular neurosecretory neurons [23]. Collectively, these SVGs reveal a coherent molecular anatomy, distinguishing inhibitory neurons, white matter, ependymal cells, and neuroendocrine centers.

In the Stereo-seq developing mouse embryo sample, Spartan captures region- and tissue-specific gene expression patterns that reflect early organogenesis (Fig. 4D; Supplementary Fig. 15B). Cardiac markers such as *Actc1, Myh7*, and *Myl7* are detected exclusively in the heart, with *Nppa* showing highly lateralized expression restricted to the left side of the heart, consistent with left–right patterning [24]. Hemoglobin genes (*Hba-a1, Hba-a2*) are expressed in the heart, liver, and fluid-filled cavity regions [25], including areas between the sclerotome and lung primordium, highlighting early erythropoietic activity. Other genes, such as *Gm42418*, are confined to cavity regions, while *Cdk8* and *Il31ra* are absent from specific tissues like the surface ectoderm, spinal cord, and brain regions, reflecting selective tissue differentiation. These findings demonstrate Spartan’s ability to detect localized embryonic expression patterns consistent with developmental biology [26, 27].

The Vizgen MERFISH mouse brain dataset further highlights Spartan’s ability to capture regionally specialized neuronal and glial populations across the adult brain (Fig. 4E; Supplementary Fig. 15C). The excitatory neuron marker *Slc17a7* [28] is strongly expressed in the cerebral cortex, hippocampus, and thalamus, whereas the astrocytic marker *Gfap* is enriched in the hippocampal and hypothalamic regions. *Chat* and *Gpr151* [29] show selective expression in the medial and lateral habenula of the epithalamus, while *Gpr88* is enriched in the striatum of the cerebral nuclei, reflecting medium spiny neuron populations [30]. The muscarinic acetylcholine receptor *Chrm1* is present throughout the brain but with lower expression in the thalamus and hypothalamus, highlighting regionally modulated cholinergic signaling [31]. Collectively, these patterns reflect both cortical and subcortical specialization, including excitatory, inhibitory, and modulatory neuronal circuits, as well as astrocytic distribution.

In the 10x Visium human dorsolateral prefrontal cortex (DLPFC) sample, Spartan identifies layer- and white matter-specific SVGs that recapitulate canonical cortical architecture (Fig. 4F; Supplementary Fig. 15D). *MBP* is highly expressed in the white matter (WM), marking oligodendrocyte populations [20], while *GFAP* is enriched in both WM and layer 1, highlighting astrocytic distribution [32] across superficial cortex and subcortical tracts. Layer-specific genes such as *COX6C* and *CST3* exhibit selective expression in combinations of deep and superficial layers (layers 1, 2, 5, and 6), whereas *FBXL16* is detected across all cortical layers except WM, indicating broad gray matter enrichment. *SCGB2A2*, although widely expressed, shows absence in portions of the left upper layers 1–4, reflecting subtle spatial heterogeneity. These findings indicate that Spartan resolves both classical layer-specific markers and fine-scale lateralized expression patterns in the human cortex.

Across all four datasets, Spartan consistently identifies genes with spatial specificity corresponding to known cell types, tissue structures, and functional domains. In the MERFISH mouse hypothalamus sample, the distinction between GABAergic nuclei, white matter tracts, and ependymal zones is clearly captured. In the Stereo-seq mouse embryo sample, tissue-specific expression reflects organogenesis and left–right patterning. In the 10x Visium human DLPFC sample, cortical layer specificity and white matter enrichment are resolved, while in the Vizgen MERFISH mouse brain dataset, regional specialization of neurons and astrocytes is detected across cortical, hippocampal, thalamic, and subcortical structures.

Notably, Spartan also captures subtle spatial heterogeneity and lateralization. Examples include *Nppa* restricted to the left heart, partial absence of *SCGB2A2* in upper cortical layers, and lateralized expression of *COX6C* in DLPFC layers. Such fine-scale patterns illustrate Spartan’s sensitivity to both broad regional patterns and localized deviations.

Overall, Spartan’s detection of spatially variable genes across multiple organs and developmental stages demonstrates robust spatial pattern detection consistent with known biology. It accurately resolves canonical markers, layer- and region-specific genes, and subtle spatial heterogeneities, making it a powerful platform for mapping molecular architecture in complex tissues.

#### Spartan preserves transferability of detected SVGs

Next, we evaluated Spartan’s ability to preserve the transferability of spatially variable genes (SVGs). A gene identified as spatially variable in one SRT sample should exhibit a similar spatial expression pattern in other samples from the same tissue type [14]. To test this, two different MERFISH samples representing the preoptic region of the mouse hypothalamus were used. The SVGs detected in one sample, *Nnat, Oxt, Gad1, Sln*, and *Mbp*, exhibited similar non-random spatial expression patterns in the other sample (Supplementary Fig. 16A-B). Importantly, SVGs that defined spatial domains in one sample also defined the same spatial domains in the other sample. For instance, *Mbp* showed high expression in the fx region in both samples, while *Sln* was restricted to the V3, MPN, BST, and PVT domains in both. Similarly, the presence of *Oxt* in the PVH region was preserved across samples. These results highlight Spartan’s ability to identify reproducible SVGs and accurately capture non-random spatial patterns.

### Visium HD analysis: Spartan resolves anatomical layering and functional compartmentalization in the developing esophagus and stomach

To evaluate Spartan in a novel and biologically complex setting, we applied it to an in-house high-resolution Visium HD section of the developing human esophagus and stomach at 10 weeks of gestation. This tissue comprises multiple epithelial cell types, concentric smooth muscle layers, heterogeneous stromal compartments, and vascular niches arranged along curved and continuous anatomical axes. The gastroesophageal junction (GEJ) represents a particularly challenging transitional region in which epithelial identity and stromal programs shift gradually rather than forming sharply demarcated boundaries. This structured resolution of a gradual transition zone suggests that activation-aware integration can partition gradual molecular gradients into spatially coherent domains, while retaining fine-scale substructure that is often lost under purely spatial smoothing.

Spartan resolved spatial domains across the entire specimen that aligned with known anatomical organization, including precise delineation of the GEJ (Fig. 5A). For this analysis, parameters were configured to emphasize the LSA component (**L**), while retaining spatial contiguity (**S**) and expression similarity (**G**), enabling detection of spatially coherent transcriptional programs within fine-grained tissue heterogeneity. Multiple domains were supported by domain-restricted spatially variable genes (SVGs), whereas additional domains reflected anatomically meaningful microstructures captured through local transcriptomic sensitivity. Notably, Spartan identified the GEJ, outer muscularis layer, esophageal mesenchyme compartment, esophageal luminal epithelium, gastric mucosal epithelium, and three quantitatively stratified gastric stromal compartments characterized by graded enrichment of extracellular matrix, glycosaminoglycan, vascular, and immune-associated programs as coherent spatial domains. Compared to NichePCA (Fig. 5B), Spartan preserved anatomically consistent boundaries and spatial continuity along curved tissue structures, consistent with more localized representation of regionally enriched programs.

**Figure 5.**
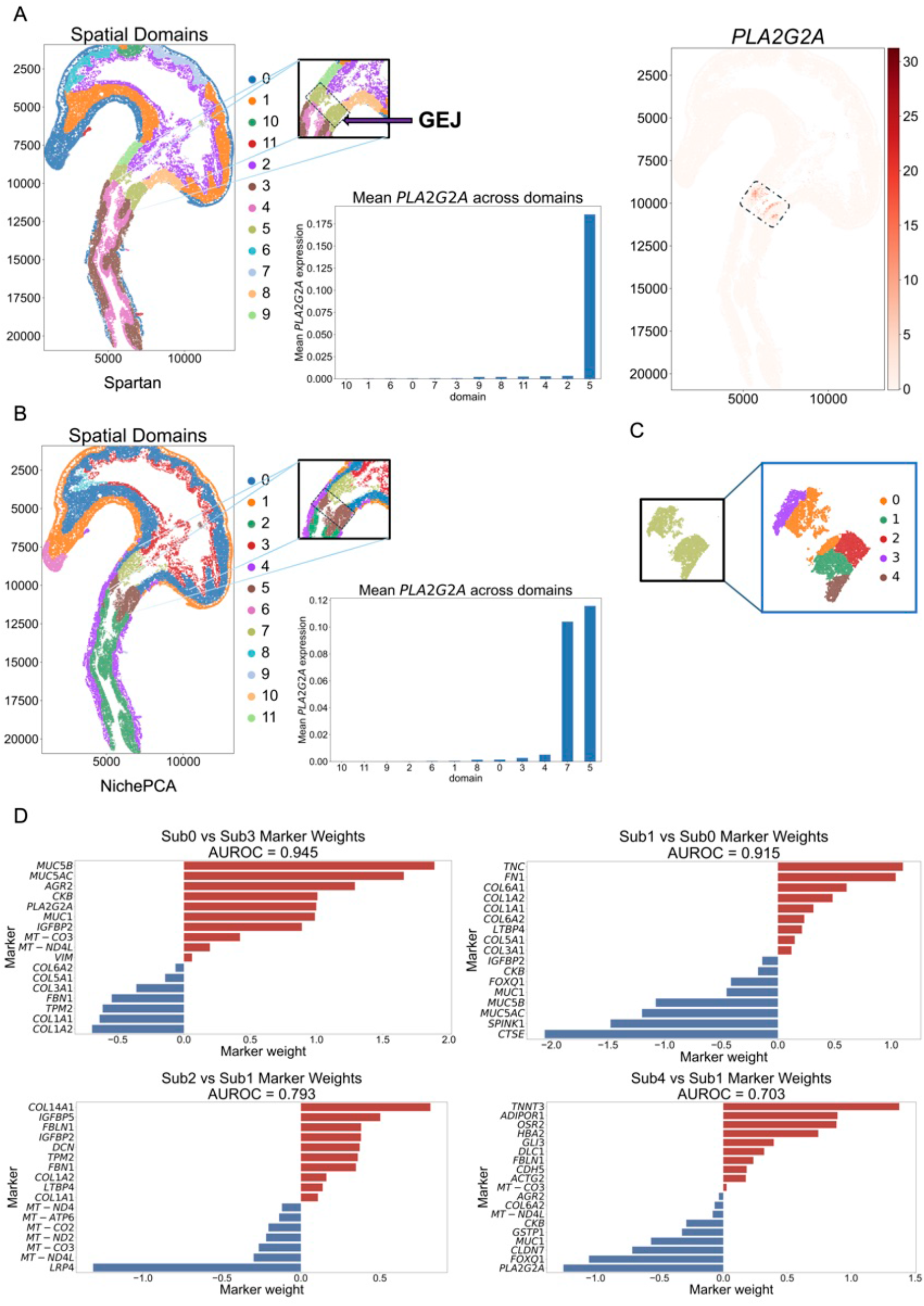
High-resolution domain reconstruction and GEJ subdomain stratification in Visium HD. (A–B) Spatial domains identified by Spartan (A) and NichePCA (B) on a Visium HD section of the developing human esophagus and stomach using 12 domains. Spartan preserves spatially coherent domains in anatomically complex regions, including the gastroesophageal junction (GEJ; Domain 5). Insets highlight the GEJ and spatial expression of *PLA2G2A*, with corresponding mean expression across domains. Domain colors are for visualization only and are not comparable across methods. (C–D) Supervised subdomain analysis within the GEJ. Subdomains were identified by reapplying Spartan to Domain 5 only. Marker weights correspond to logistic-regression coefficients from pairwise subdomain classification, and AUROC values (0.703–0.945) quantify subdomain separability.

Spartan resolved the GEJ as domain 5 (Fig. 5A), defined by the presence of SVGs including *PLA2G2A*, gastric-associated mucins (*MUC5AC, MUC5B*), and squamous epithelial-associated markers such as the transcription regulator *ELF3*, and *TACSTD2* [33, 34] (Supplementary Fig. 17A-G). Co-expression of gastric-associated mucins alongside esophageal markers like *TACSTD2* supports a mixed transcriptional identity consistent with a junctional transition zone. Notably, *PLA2G2A* exhibited selective enrichment within this domain, whereas NichePCA distributed the signal across multiple adjacent partitions (Fig. 5A-B), further supporting more localized representation of regionally enriched programs by Spartan. *PLA2G2A* has been reported to function as an antimicrobial effector and modulator of gut microbiota, supporting epithelial barrier associated functions in the gastrointestinal tract [35]. Together, these findings are consistent with activation-aware integration contributing to stable partitioning at transitional interfaces.

To further characterize spatial structure within the GEJ, we recursively applied Spartan within domain 5 to identify subdomains capturing localized transcriptional programs (Fig. 5C). Adjacent subdomains were differentiated using pairwise differential expression (DE) analysis, and discriminative marker sets were evaluated using logistic regression, yielding AUROC scores ranging from 0.703 to 0.945, indicating robust separability.

Subdomain 0 (Sub0) represented a secretory gastric epithelial-like compartment oriented towards the lumen, reflected by mucin-associated expression patterns (Supplementary Fig. 17B-C). Pairwise comparison against a left-side stromal subdomain (Sub3) yielded high discriminative performance (AUROC=0.945), consistent with distinct epithelial and collagen-rich stromal molecular states (Fig. 5D). On the right side of the GEJ, a remodeling-associated stromal compartment (Sub1) was distinguished by extracellular matrix reorganization programs (AUROC=0.915). Additional right-side subdomains captured deeper stromal niches with separable signaling and infrastructure associated transcriptional profiles (AUROC range 0.703–0.793)(Fig. 5D), illustrating the ability of Spartan to resolve nested spatial microenvironments within a continuous transition zone. This analysis revealed asymmetry across the junction, with a coherent mucosal and stromal interface on the left side, and a multi-layered arrangement with distinct matrix remodeling and signaling associated compartments on the right side.

Beyond the GEJ, Spartan resolved the concentric multi-laminar architecture of the fetal esophagus (Fig. 6A), identifying an outer muscularis layer (domain 0), an intermediate fibroblast-rich mesenchymal stroma (domain 3), and an inner luminal epithelium (domain 4) (Fig. 6B). In contrast, NichePCA grouped the luminal epithelium and intermediate stroma into a single domain. The muscularis externa (domain 0) formed a coherent contractile scaffold defined by smooth muscle associated SVGs, anchored by *TAGLN* [36] (Fig. 6B-D; Supplementary Fig. 18A-E). Quantitative analysis confirmed maximal enrichment within domain 0 and reduced enrichment at the GEJ, consistent with distinct molecular prioritization at the junction relative to surrounding muscle (Fig. 6C; Supplementary Fig. 18F-G). Domain 3 was supported by extracellular matrix associated stromal programs anchored by *FN1*, consistent with a fibroblast-rich supportive compartment (Fig. 6B-D; Supplementary Fig. 19). Domain 4 was supported by epithelial stratification associated expression patterns, including *KRT5*, a canonical basal epithelial marker in stratified epithelia [37] (Fig. 6B-D; Supplementary Fig. 20).

**Figure 6.**
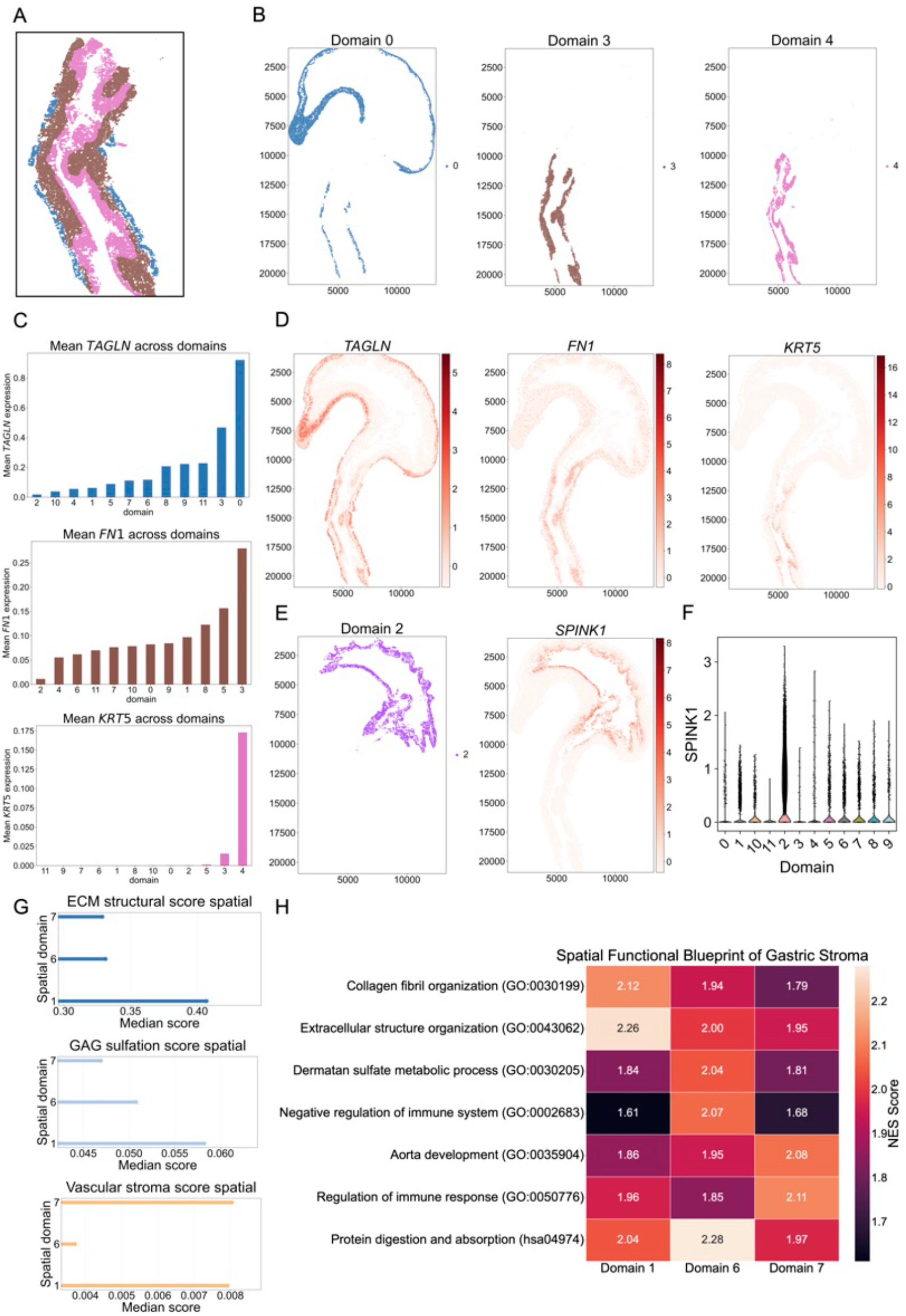
Reconstruction of esophageal laminar architecture and stromal functional programs using Visium HD. (A) Spartan resolves the concentric laminar organization of the developing esophageal wall, delineating epithelial, stromal and muscular compartments. (B) Spatial domains corresponding to outer muscularis (Domain 0), stromal interface (Domain 3), and luminal epithelium (Domain 4). (C) Mean expression of canonical markers across domains. *TAGLN* is enriched in Domain 0, *FN1* in Domain 3, and *KRT5* in Domain 4. (D) Spatial expression maps of *TAGLN, FN1*, and *KRT5* align with domain boundaries. (E) Gastric mucosal epithelium (Domain 2) with localized *SPINK1* expression. (F) Violin plot confirming *SPINK1* enrichment in Domain 2. (G) Median spatially smoothed module scores across stromal domains (Domains 1, 6, 7) reveal continuous functional gradients across adjacent compartments. (H) Gene set enrichment heatmap summarizing domain-associated programs: Domain 1 shows extracellular matrix enrichment, Domain 6 sulfated proteoglycan and immune-regulatory pathways, and Domain 7 vascular-associated signaling.

Spartan also identified the gastric mucosal epithelium (domain 2), characterized by a spatially coherent secretory program and epithelial identity (Fig. 6E). This compartment was supported by enrichment of a set of gastric epithelial marker genes including the gene encoding the mucosal protective protein *SPINK1* [38, 39] (Fig. 6E-F; Supplementary Fig. 21).

Surrounding this epithelium, Spartan resolved the gastric stroma into three specialized spatial domains, capturing regional heterogeneity that NichePCA grouped as a single domain. Domain 1 represents the structural framework of the gastric corpus, exhibiting the highest median extracellular matrix (ECM) and glycosaminoglycan (GAG) sulfation scores (Fig. 6G-H). Domain 6 represents the antral reservoir, showing peak enrichment for protein digestion and the dermatan sulfate metabolic process, while also contributing to immune regulation (Fig. 6G-H). Domain 7 is associated with deeper submucosal layers and is defined by programs linked to vascular development and immune response regulation (Fig. 6G-H).

Beyond coarse domain recovery, Spartan identifies localized stromal microstructures that provide a mechanistic justification for sub-compartmentalization. Domain 10, a spatially restricted niche embedded within gastric stroma, exhibits strong enrichment for erythroid-associated genes such as *SLC4A1* and *HBA2* (Supplementary Fig. 22A-D). The corresponding functional enrichment patterns highlight oxygen transport and heme-related biology (Supplementary Fig. 22E) supporting the interpretation that this region represents a blood-associated microenvironment rather than a generic fibroblast compartment. Stromal regions are frequently grouped into broad partitions by neighborhood-based approaches, whereas activation-aware integration supports separation of smaller but biologically coherent niches.

Similarly, the analysis of Domains 8 and 9 supports Spartan’s ability to distinguish closely adjacent stromal subregions on either side of the lumen at the entrance to the stomach following the gastroesophageal junction (Supplementary Fig. 23A). While both domains belong to a broadly stromal context, their differential signatures suggest that they encode distinct microenvironments rather than noisy fragmentation (Supplementary Fig. 23B). Domain 8 shows strong enrichment of extracellular matrix-associated markers, including fibrillar collagens and matrix remodeling genes, exemplified by *FBLN1*, consistent with a structural stromal compartment (Supplementary Fig. 23B and Supplementary Fig. 23C). In contrast, Domain 9 is characterized by reduced levels of these markers but moderately enriched vascular-associated transcripts such as *PECAM1, CLDN5, PLVAP*, and *EGFL7*, supporting a vessel-associated stromal niche (Supplementary Fig. 23B and Supplementary Fig. 23D). Logistic regression based discrimination provides a quantitative measure of separability between these domains, further strengthening the argument that Spartan is resolving functional microstructure rather than producing arbitrary clusters (Supplementary Fig. 24). Together, these results demonstrate that Spartan can resolve spatially coherent anatomical layers and functionally distinct stromal compartments in a complex developing tissue.

To further assess whether the gene programs underlying the spatial domains identified by Spartan reflect meaningful spatial structure, we evaluated the spatial properties of SAQ-ranked genes within this dataset.

#### Validation of SAQ-based SVGs in Visium HD sample

To validate the spatial relevance of SAQ-based gene ranking, we evaluated whether genes with high SAQ scores exhibit stronger spatial autocorrelation than background genes (Supplementary Methods; Supplementary Fig. 25). Moran’s I statistics were computed for all genes using the spatial neighbor graph. The mean Moran’s I for top-ranked genes was 0.3048, whereas candidate background genes exhibited a mean Moran’s I of 0.0153, indicating stronger spatial autocorrelation among SAQ-ranked genes.

The distribution of Moran’s I values for top-ranked SAQ genes was compared with that of candidate background genes using a Mann–Whitney U test (Supplementary Fig. 25), yielding a highly significant difference (*p* ≈ 1.2×10^−19^). Enrichment of significantly spatially autocorrelated genes among the top SAQ genes was further evaluated using Fisher’s exact test (p-value = 5.68 × 10^−7^, odds ratio = ∞) (Supplementary Fig. 25).

Finally, spatial expression maps of the top six SAQ-ranked genes illustrate localized expression patterns consistent with spatial structure (Supplementary Fig. 26).

## Discussion and conclusion

Spatial transcriptomics is rapidly transitioning from coarse tissue maps to high-resolution profiling, where tens of thousands of spatial locations capture subtle biological structure along continuous anatomical axes. This shift exposes a computational limitation: many clustering approaches separate broad compartments but blur tissue boundaries, merge transitional zones, or miss localized microstructures that are biologically meaningful, particularly in developmental tissue where programs vary gradually yet change sharply at interfaces. Spartan is designed to address this challenge through an aggregated graph formulation that integrates spatial topology, Local Spatial Activation (LSA), an activation-aware component designed to capture localized transcriptomic deviation relative to neighborhood context, complementing spatial topology and gene expression connectivity.

Conceptually, LSA draws inspiration from local spatial statistics in that it evaluates neighborhood context; however, unlike classical autocorrelation measures, LSA is formulated as a non-negative graph signal in PCA-reduced space, enabling direct integration into weighted community detection frameworks. Within transcriptionally coherent domains, LSA values tend to be low because neighboring locations exhibit minimal deviation from their local context. At spatial transitions or nested microstructures, LSA increases when joint deviation from the neighborhood exceeds background variance. When integrated with gene expression connectivity, this activation-aware signal contributes additional contrast at localized transitions, influencing community detection toward partitions that better respect anatomical interfaces.

Spartan remains computationally practical as datasets increase in resolution and gene panel size (Supplementary Figs. 27–28). Runtime scales approximately linearly with the number of spatial locations, and high-resolution datasets such as Visium HD are processed within minutes on CPU. Efficiency derives from sparse local graphs and linear-time community detection, avoiding dense pairwise computations or model training. These properties position Spartan as a computationally efficient and scalable framework that supports high-resolution datasets without reliance on model training or specialized hardware.

A practical advantage of Spartan is that it does not require ground-truth labels. Ground truth was used primarily for benchmarking and ablation studies. In unlabeled datasets, Spartan can be applied by varying the Leiden resolution to obtain partitions at the desired level of biological granularity. Optional stability-based pruning metrics may further constrain the search space but should be viewed as workflow guidance rather than definitive unsupervised proof of a universally optimal partition.

Spartan was designed for robustness to parameterization. Clustering performance varies smoothly across a broad range of multiplex weight allocations (*β*_1_, *β*_2_) without sharp optima (Supplementary Fig. 29), indicating that improvements are not driven by fragile tuning. For benchmarking, a single dataset-level *α* was selected using a cross-sample consensus procedure and subsequently fixed, avoiding per-sample optimization (Supplementary Fig. 30); performance remains stable across a wide *α* range. The smooth dependence on multiplex weights clarifies the role of LSA: its contribution can be increased without destabilizing clustering, which is particularly advantageous in high-resolution datasets with sharp spatial transitions, such as Visium HD and Vizgen MERFISH, where increased activation weighting enhances sensitivity to localized transitions while preserving spatial coherence. In this framework, LSA provides a structured signal that highlights localized transcriptional transitions within spatial neighborhoods.

We demonstrate Spartan’s biological relevance using a cell-level Visium HD section of developing human esophagus and stomach at 10 weeks of gestation, containing layered smooth muscle, fibroblast-rich stroma, glandular epithelium, vascular niches, and the complex gastroesophageal junction (GEJ). Spartan resolves spatial domains across the specimen and separates the GEJ as a distinct transitional territory consistent with its mixed molecular identity. Recursive application within the GEJ reveals asymmetrically organized subdomains, supporting the interpretation that activation-aware graph integration enhances sensitivity to nested microenvironments in continuous tissue regions relative to expression-only or topology-only integration.

To assess robustness to clustering granularity, we varied the Leiden resolution for the Visium HD sample to obtain cluster counts spanning coarse to fine partitioning (*R* = 7–20) while fixing all other hyperparameters (Supplementary Figs. 31-32). Spartan exhibits stable behavior across this range. At low resolution (*R* = 7), it collapses into spatially contiguous macro-domains reflecting global tissue organization while preserving the GEJ as a coherent transition region. At higher resolution (*R* = 16–20), it introduces local subdomains through hierarchical refinement without disrupting anatomical continuity or fragmenting the GEJ. In contrast, NichePCA shows reduced anatomical alignment at low resolution and increasingly fragmented or stripe-like partitions at higher cluster counts.

Finally, Spartan extends beyond domain discovery to spatially variable gene detection. By leveraging activation-aware structure, Spartan identifies genes associated with localized transitions and domain-restricted programs, supporting interpretability and enabling mechanistic follow-up. Spartan therefore functions not only as a clustering method but also as a framework for investigating how spatial organization relates to localized transcriptional programs.

In conclusion, Spartan introduces an activation-aware multiplex graph framework that integrates spatial topology, Local Spatial Activation, and gene expression connectivity. The method is robust to parameter variation while permitting adaptive weighting in high-resolution settings where microdomain boundaries are critical. By linking Local Spatial Activation to principles of local spatial autocorrelation in geostatistics, Spartan provides an interpretable and extensible framework for spatial systems-level analysis, with potential applications in cell–cell communication, developmental tissue remodeling, and spatial multi-omics integration.

## Methods

### Data preprocessing

Raw SRT data were processed using standard workflows described previously [40, 41]. Gene expression matrices were library-size normalized using Scanpy’s *pp*.*normalize total* and log-transformed using *pp*.*log1p* [40]. The resulting filtered log-normalized matrix is denoted as **X** ∈ ℝ^*N*×*M*^, where *N* represents the number of spots (or cells) and *M* the number of genes. For sequencing-based technologies such as 10x Visium, Visium HD, and Stereo-seq, highly variable genes (HVGs) were selected using Scanpy’s *pp*.*highly variable genes* functionality [40]. Throughout this work, each row **X**_*i*_ is referred to as a spot; the formulation applies equally when **X**_*i*_ represents a cell (e.g., MERFISH).

Principal component analysis (PCA) was applied to **X** to obtain a lower-dimensional representation of spots in principal-component space, **Y**^**pca**^ ∈ ℝ^*N*×*P*^, where *P* ≪ *M* is the number of principal components. The feature matrix is represented as

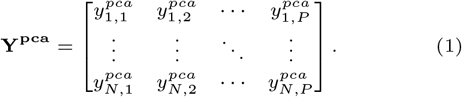

Equivalently, **Y**^**pca**^ can be expressed in terms of row vectors:

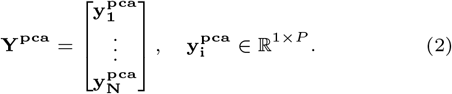

Spartan is also implemented within the `SpatialData` framework [42], enabling standardized multimodal I/O and interoperability across spatial omics data types.

### Spatial graph

The spatial graph is expressed as *gr*^s^ = (*V*_s_, *E*_s_) where *V*_s_ = {1, 2, ···, *N*} and *E*_s_ is represented in the form of a binary adjacency matrix, **S** ∈ ℝ^*N*×*N*^. Spatial neighbor graphs were constructed using Squidpy [41]. Each element of the matrix **S**, *s*_*ij*_, represents:

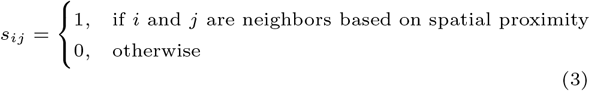

The spatial proximity is determined by constructing either: (i) a *k*-nearest neighbor (KNN) graph based on Euclidean distance in 2D space using spatial coordinates, where for each spot *i* its *k* nearest spatial neighbors are identified. The KNN construction yields a directed adjacency by default; we optionally symmetrized it depending on the `Squidpy` coordinate setting. Alternatively, (ii) a Delaunay graph is constructed, in which spot *i* shares edges with neighboring spots defined by the Delaunay triangulation of the spatial coordinates. The Delaunay construction yields an undirected planar graph. Accordingly, **S** is generally asymmetric under the directed KNN construction and symmetric under the Delaunay construction. The matrix **S** can be represented in terms of row vectors:

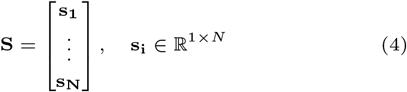

Each vector **s**_**i**_ = (*s*_*i*1_, ···, *s*_*iN*_) represents the spatial neighborhood of spot *i* where each *s*_*ij*_ is defined according to Eq. 3. Each row **s**_**i**_ contains nonzero entries corresponding to the spatial neighbors of spot *i* (with out-degree *K*_*i*_ in the directed KNN case), and the matrix is stored in sparse format.

#### Row-normalized spatial weights matrix

The spatial weights matrix, **W** ∈ ℝ^*N*×*N*^, is constructed where each element of the matrix *w*_*ij*_ represents the measure of spatial proximity between spot *i* and spot *j*. The spatial weight *w*_*ij*_ is calculated as:

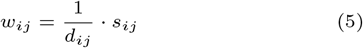

where *d*_*ij*_ is the Euclidean distance between spatial coordinates of spots *i* and *j*. To ensure numerical stability, distances were offset by *ϵ* = 10^−3^, i.e., 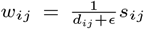. We row-normalized **W** to obtain 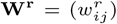, where 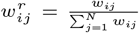. The matrix **W**^**r**^ is stored as a sparse matrix. Distances *d*_*ij*_ were computed only for graph-connected pairs (*i, j*) using the neighbor graph construction utilities in `Squidpy` [41]. The **S** provides a binary connectivity structure based on raw spatial proximity, while the row-normalized weight matrix **W**^**r**^ refines this by accounting for distance. We use the row-normalized matrix **W**^**r**^ to construct the matrix **L** in the next section.

### Local Spatial Activation (LSA) graph

The LSA graph is expressed as *gr*^lsa^ = (*V*_lsa_, *E*_lsa_) where *V*_lsa_ = {1, 2, ···, *N*} and *E*_lsa_ is represented as the weighted adjacency matrix, **L** ∈ ℝ^*N*×*N*^. **L** captures neighborhood-relative transcriptional deviation between spatially adjacent spots. Each element *l*_*ij*_ gives the LSA weight associated with the spatial edge from spot *i* to spot *j*.

We divide the calculation of the LSA weight *l*_*ij*_ for a spatial edge *i* → *j* into three components:

#### Summary statistics of spot i

We calculate two summary statistics: the mean and variance of the closed neighborhood 𝒩_*i*_ of spot *i*. The closed neighborhood 𝒩_*i*_ consists of spot *i* and its *K*_*i*_ spatial neighbors, as determined by the matrix **S**.

We calculate the mean feature vector of N_*i*_ as:

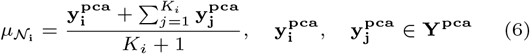

where 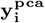 and 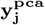 are the feature vectors of spot *i* and spot *j*, respectively.

We calculate the local variance and the local variance norm of the neighborhood 𝒩_*i*_ as follows.

First, define the deviation vectors:

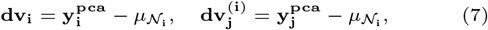

where **dv**_**i**_ and 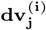 denote deviations from the neighborhood mean.

The neighborhood variance vector is then computed using elementwise products:

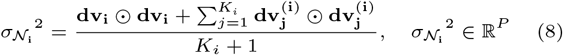

where ⊙ denotes elementwise (Hadamard) multiplication.

The local variance norm of the neighborhood 𝒩_*i*_ is defined as

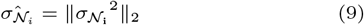

The mean feature vector, 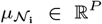, summarizes the transcriptional context of the surrounding tissue (neighborhood) and provides a smooth representation of the local expression trends. The local variance norm of the neighborhood 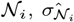, is defined as the Euclidean norm (‖ · ‖_2_) of the neighborhood variance vector 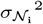. This *ℓ*_2_ aggregation provides an isotropic summary of multivariate variance across principal components and is invariant to orthogonal rotations of the PCA coordinate system.

#### Attribute deviation

We measure the attribute deviation of spot *i* from its surroundings, 𝒩_*i*_:

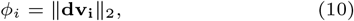

The quantity *ϕ*_*i*_ is the attribute deviation norm, measuring how much spot *i* deviates from its neighborhood in PCA-reduced expression space. Lower values indicate that spot *i* is transcriptionally similar to its neighborhood mean, whereas higher values indicate stronger local deviation.

Similarly, we calculate the attribute deviation of neighbor *j* of spot *i*, from 𝒩_*i*_ as:

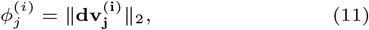

Note that 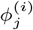 is computed with respect to 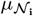, making **L** asymmetric by construction.

#### Local Spatial Activation (LSA)

The LSA weight associated with the spatial edge *i* → *j* is defined as:

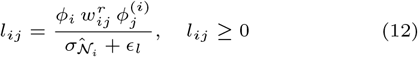

Here, *l*_*ij*_ consists of three key components: the quantities *ϕ*_*i*_ and 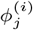 measure the magnitude of transcriptional deviation of spots *i* and *j* relative to the local neighborhood of spot *i*, while 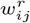 denotes the row-standardized spatial proximity weight between spots *i* and *j*. The small quantity *ϵ*_*l*_ ensures numerical stability. The local variance norm 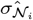 normalizes LSA values relative to local neighborhood variability. LSA is formulated as a non-negative activation signal on the spatial neighbor graph rather than as a statistical test of spatial autocorrelation. Thus, *l*_*ij*_ encodes a variance-normalized, spatially weighted joint transcriptional deviation relative to the local neighborhood context of spot *i* within *gr*^lsa^. The resulting matrix **L** is therefore an asymmetric weighted adjacency matrix and serves as a key component integrated into the aggregated graph (Eq. 15), where it is balanced against gene expression connectivity and spatial proximity.

### Gene expression connectivity graph

We construct a gene expression connectivity graph in the form of a KNN graph in the PCA space, *gr*^g^. The *gr*^g^ is expressed as (*V*_g_, *E*_g_) where *V*_g_ = {1, 2, ···, *N*} and *E*_g_ is represented by the binary adjacency matrix, **G** ∈ ℝ^*N*×*N*^. The *gr*^g^ encodes the neighborhood relationship between two spots based on their transcriptomic similarity. The transcriptomic distance in PCA space between spots *i* and *j*, 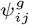, was computed as:

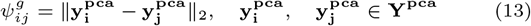

The adjacency matrix **G** encodes directed KNN neighborhood connectivity. Each entry of **G**, denoted *g*_*ij*_, is defined as:

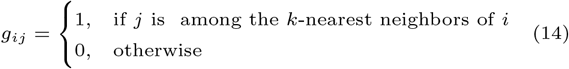

Here, 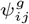 is used to identify the *k* nearest neighbors for each spot in PCA space. The adjacency matrix **G** forms the gene expression connectivity component of the aggregated graph (Eq. 15), complementing LSA and raw spatial proximity.

### Aggregated graph

The aggregated graph *gr*^mul^ is a directed graph that combines three components: *gr*^s^, *gr*^lsa^, and *gr*^g^. The aggregated graph, *gr*^mul^, is expressed as (*V*_mul_, *E*_mul_) where *V*_mul_ = {1, 2, ···, *N*} and *E*_mul_ is represented by an asymmetric weighted adjacency matrix **J** ∈ ℝ^*N*×*N*^ and is formed as:

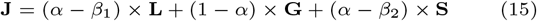

where *α* ∈ (0, 1) and *β*_1_ + *β*_2_ = 0.5 with *β*_1_, *β*_2_ ≤ *α*. In this formulation, the matrix **J** balances three complementary sources of structure: **L** that encodes LSA information of all spots, the gene expression connectivity matrix **G**, and the raw spatial adjacency matrix **S**. An entry *j*_*ij*_ of the matrix **J** is high when spot *j* belongs to the neighborhood N_*i*_ (the neighborhood of spot *i*) and is supported either by high transcriptomic similarity or strong joint deviation relative to the neighborhood N_*i*_. The parameter *α* regulates the overall trade-off between spatial and expression-derived connectivity. At the same time, the correction terms *β*_1_ and *β*_2_ ensure that the combined weight of the structural terms **L** and **S** equals 2*α* − 0.5, thereby controlling the total structural contribution relative to the expression-based connectivity term **G**. We set *β*_1_ +*β*_2_ = 0.5 so that *α* controls the overall structural−transcriptomic trade-off while *β*_1_ and *β*_2_ set the relative allocation between **L** and **S**. With *β*_1_ and *β*_2_ fixed within each experimental configuration, *α* is the sole free parameter governing this trade-off. This yields a balanced integration of spatial and expression-derived information. No additional per-layer rescaling was applied. Since *s*_*ij*_, *g*_*ij*_ ∈ {0, 1} and *l*_*ij*_ ≥ 0, **J** is non-negative, satisfying the requirements of modularity-based community detection.

### Spatial biological partitioning

We use the Leiden clustering algorithm to partition the aggregated graph *gr*^mul^ = (*V*_mul_, *E*_mul_). We use the RBConfigurationVertexPartition quality function (leidenalg), which supports directed graphs. The objective *Q*(**J**, *γ*) uses the weighted adjacency matrix **J** and resolution parameter *γ*. It partitions *V*_mul_ into *R* non-overlapping clusters {*C*_1_, *C*_2_, ···, *C*_*R*_}. The quality function *Q*(**J**, *γ*) maximizes the total edge weight within clusters relative to a configuration-model null. In this case, we refer to these clusters as spatial domains.

In practice, the resolution parameter *γ* controls the granularity of spatial domains. Together with the layer parameters (*α* − *β*_1_, 1 − *α, α* − *β*_2_), it allows flexible adjustment across SRT datasets of varying spatial scales and technologies. This approach allows the algorithm to leverage the connectivity information encoded by **S** and **G**, which provide neighborhood guidance, and the matrix **L**, which introduces neighborhood-relative activation structure not captured by spatial proximity or expression-derived connectivity alone. Modularity optimization on **J** therefore favors partitions that concentrate activation-weighted and connectivity-supported interactions within clusters while penalizing spurious cross-domain links. This formulation enables the detection of spatial domains that reflect both spatial cohesion and localized transcriptional heterogeneity.

### *α*-selection strategy

We formulated a hierarchical *α*-selection strategy for determining the multiplex weighting parameter *α* of the matrix **J**. This *α*-selection procedure was used only in benchmarking and ablation on annotated datasets, and was not used in unlabeled biological analyses. Annotations were used only to set dataset-level hyperparameters for evaluation.

For a given multi-sample SRT dataset *D* = {Sm_1_, ···, Sm_*T*_}, where Sm_*i*_ denotes the *i*^th^ sample and *T* is the total number of samples, the *α*-selection strategy consists of four stages:

1. **Range selection**. For each sample Sm_*i*_ in *D*, we defined the search ranges of the multiplex weighting parameter *α* ∈ [*α*_*l*_, *α*_*r*_] and the resolution parameter *γ* ∈ [*γ*_*l*_, *γ*_*r*_]. For every candidate *α*_*i*_ ∈ [*α*_*l*_, *α*_*r*_], we performed Leiden clustering across all *γ* values, yielding a set of clustering assignments 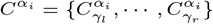.
2. **Cluster-count alignment**. For benchmarking datasets with curated annotations, we retained clustering assignments whose number of clusters matched the annotated domain count, forming 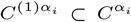. This cluster-count alignment follows standard SDMBench benchmarking conventions. For each 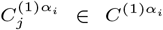, we computed the evaluation metrics ARI, NMI, HOM, COM, together with nLSAS.
3. **Spatial-stability prior**. We further refined 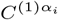 by retaining only clustering assignments whose nLSAS values lay within the central stability band (40^th^–70^th^ percentiles by default). The nLSAS score is fully unsupervised and derived solely from the dataset-specific LSA graph and the predicted partition structure. This percentile-based filtering removes spatially fragmented partitions (low nLSAS) and over-smoothed partitions (high nLSAS), thereby favoring configurations that balance activation coherence with domain resolution. The resulting pruned set 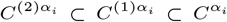 was used to compute the median of each evaluation metric across all 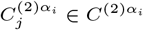. For each sample Sm_*i*_, we thus obtained a collection of candidate *α* values and their corresponding median performance summaries, stored in a table 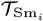, where each row represents one *α*_*i*_ and its associated summaries. Each table 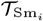 was ranked by median NMI, and only the top *K* candidates were retained to mitigate the influence of outliers (default *K*=50). The resulting table 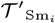 therefore contains high-performing structural weight configurations for that sample. Although ranking was performed using median NMI for stability, final performance evaluation was conducted using multiple metrics, as described in the benchmarking Results section Spartan demonstrates competitive and statistically supported performance on public SRT datasets. Repeating Steps 1–3 for all samples in *D* yields a collection of tables 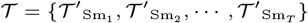.
4. **Cross-sample consensus**. Let 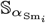 denote the set of *α* values contained in 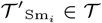. We computed the intersection of all such sets to obtain a consensus set of *α* values, 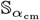, thereby enforcing cross-sample generalizability by restricting selection to *α* values that were consistently stable across samples (when the intersection was non-empty). In cases where the consensus intersection is empty, the union of top-*K* candidates is used with cross-sample averaging.

For each 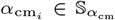, we extracted its median performance summaries from all 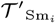 tables and calculated the mean of the median NMI values across samples. The 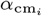 yielding the highest mean of the per-sample median NMI values under this criterion was designated as *α*_best_.

The resulting *α*_best_ represents a dataset-level structural weighting parameter for the matrix **J**. Once selected, *α*_best_ was fixed for all samples within that dataset and not reoptimized per sample or per resolution.

### Parameter selection

#### Ablation study

The *k* value for the KNN graph was set to 10 for constructing the spatial graph and to 15 for constructing the gene expression connectivity graph. The parameter *α*, which balances spatial and transcriptomic contributions, was varied between 0.5 and 0.9. Two additional parameters, *β*_1_ and *β*_2_, controlled the relative weighting of the LSA and spatial neighborhood connectivity components; these were fixed at 0.26 and 0.24, respectively, to emphasize spatial connectivity slightly more strongly. The resolution parameter *γ* for the Leiden algorithm was varied between 0.5 and 0.9. For all datasets, the number of ground truth clusters was provided as input for evaluation.

#### Benchmarking on SDMBench

The benchmarking parameters for Spartan were configured as follows. In feature selection and graph construction, the number of principal components (*P*) was fixed at 30 for all datasets. A total of 3,000 highly variable genes (HVGs) were selected, following [10], with HVGs required only for the Stereo-seq dataset. The *k* value for constructing the *gr*^*s*^ KNN graph was set to 10, and the *gr*^*g*^ KNN graph was constructed using 15 nearest neighbors. Except for the Stereo-seq dataset, the number of immediate neighbors was set to 4 and the ring size was set to 1.

Parameter ranges were defined as described in Section *α*-selection strategy. The weight parameter *α* was bounded between 0.5 and 0.9 across datasets. The parameters *β*_1_ and *β*_2_ were fixed at 0.26 and 0.24, respectively. The resolution parameter *γ* was varied between 0.5 and 0.9 for most datasets but extended to the range [0.5, 1.2] for osmFISH and Stereo-seq. The percentile thresholds for nLSAS were fixed at the 40 and 70 percentiles for all datasets, except for STARmap where they were fixed at 40 and 80.

#### Benchmarking against recently proposed methods

We used the same benchmarking pipeline for Spartan to evaluate its performance on five SDMBench datasets as discussed above. To evaluate NichePCA and BANKSY on SDMBench datasets, we adopted a similar benchmarking pipeline as detailed in [9]. Specifically, for each dataset, the number of nearest neighbors was varied between 5 and 20, and the value yielding the highest mean NMI across samples within that dataset was selected. For SpatialLeiden, we chose their best-performing configuration as reported in their SDMBench benchmarking, namely the configuration that uses a KNN graph to represent spatial connectivity. We adopted their technology-specific layer weight ratios as used in their SDMBench benchmarking.

For the Vizgen MERFISH dataset, we employed a different benchmarking workflow similar to that used in NichePCA. Specifically, we restricted the maximum number of clusters to 60, consistent with the evaluation protocol used in NichePCA. The range selection step was similar to Section Benchmarking on SDMBench, but here *α* was varied between 0.77 and 0.87, *γ* was varied between 2.0 and 4.0 for the Delaunay configuration and between 1.0 and 2.0 for the KNN configuration. The parameters *β*_1_ and *β*_2_ were fixed at 0.20 and 0.30, reflecting the higher spatial resolution characteristic of imaging-based datasets. For a given configuration (KNN/Delaunay), we selected a single dataset-level *α* by, for each candidate *α*, taking the maximum NMI achieved over the predefined resolution sweep (subject to the cluster cap), and then selecting the *α* that maximized this value consistently across samples. Similarly, for NichePCA and BANKSY, we varied the number of nearest neighbors between 5 and 29 and the *γ* between 1.0 and 2.0 for NichePCA, 2.0 and 4.0 for BANKSY, and selected the dataset-level *k* yielding the highest NMI across samples under the same resolution sweep. This applied dataset-level parameter selection across methods.

For all benchmarking comparisons, parameter ranges for competing methods were explored within the bounds recommended by their respective publications, and configurations were selected according to their reported benchmarking strategies. No additional dataset-specific supervision was introduced beyond what was described in the respective original studies. This ensures that cross-method comparisons reflect methodological design rather than asymmetric tuning procedures.

### Spatial Activation Quotient

To identify genes with non-random spatial expression patterns [spatially variable genes (SVGs)], we introduce the *Spatial Activation Quotient* (SAQ), denoted by **v**^**saq**^ ∈ ℝ^*M*^. This metric ranks genes according to the degree to which their expression profiles exhibit localized spatial structure rather than random variation.

We begin with the filtered gene expression matrix **X** ∈ ℝ^*N*×*M*^, where *N* is the number of spots and *M* is the number of genes. Each gene vector is first column-wise mean-centered to remove global expression offsets:

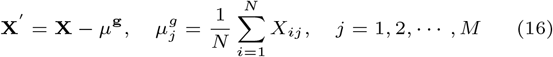

where **X**′ ∈ ℝ^*N*×*M*^ is the mean-centered expression matrix and *μ*^**g**^ ∈ ℝ^*M*^ is the gene-wise mean expression vector.

Next, we propagate the mean-centered expression through the LSA graph to obtain an LSA-weighted neighborhood representation of the expression for each gene:

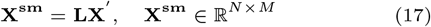

Here, **X**^*sm*^ represents an LSA-weighted neighborhood aggregation (with weights given by **L**). **L** is the weighted adjacency matrix of the LSA graph. The product **LX′** represents a spatially weighted smoothing of expression values, such that each spot’s value reflects its local neighborhood context. For SAQ scoring, **L** is treated as a fixed spot–spot operator constructed from PCA-reduced spot embeddings, whereas gene scores are computed in the original mean-centered gene expression space.

We then quantify the alignment between the original expression profile of a gene and its spatially smoothed profile. For gene *j*, this is given by:

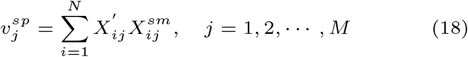

Equivalently, if 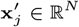 denotes the mean-centered expression vector of gene *j* across all spots (i.e., the *j*^th^ column of **X**′), then

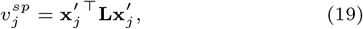

highlighting that SAQ corresponds to a graph-weighted quadratic form measuring spatial coherence of gene *j*. This quadratic-form interpretation also motivates the permutation-based null model used for SVG identification, described below.

To normalize for gene-specific variability and expression magnitude, we compute the squared *ℓ*_2_ norm of each gene’s mean-centered profile:

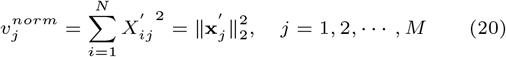

Finally, the spatial activation quotient is defined as the ratio:

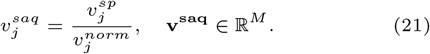

This normalization rescales the graph-weighted quadratic form by total gene variance, yielding a dimensionless measure of relative spatial coherence across genes.

### Spatially variable gene identification

We computed the Spatial Activation Quotient (SAQ) for each gene (*M* total) using the weighted adjacency matrix **L**, which represents the LSA graph of a given sample. The SAQ can be interpreted as a normalized graph quadratic form, analogous to a Rayleigh quotient [43], providing a measure of how strongly a gene’s expression pattern aligns with the spatial neighborhood structure encoded by **L**. Genes with higher SAQ values exhibit stronger non-random spatial organization and are therefore candidates for spatially variable genes (SVGs).

To assess statistical significance, we performed permutation-based calibration under a spatial-randomness null model. Specifically, for each permutation we randomly permuted the spot indices within each mean-centered gene vector 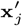, disrupting its alignment with the fixed graph operator **L** while preserving the marginal expression distribution. For each permuted instance, the SAQ statistic was recomputed to generate a gene-wise null distribution reflecting the range of values expected under spatial randomness.

Rather than estimating empirical tail probabilities, we estimated the null mean and variance of the SAQ from permutation scores and computed a standardized *z*-score for each observed value. One-sided *p*-values were obtained from the upper tail of the standard normal distribution, reflecting enrichment for positive spatial coherence under **L**.

Finally, we applied False Discovery Rate (FDR) correction across all genes using the Benjamini–Hochberg procedure [44]. Genes passing the threshold (FDR *<* 0.05) were considered significantly spatially variable genes (SVGs).

### Evaluation metrics

We evaluated spatial domain identification using the following metrics:

#### Adjusted Rand Index

The Adjusted Rand Index (ARI) measures the agreement between a predicted clustering and a ground truth partition while correcting for random label assignments. Given two clustering assignments: *U* = {*U*_1_, *U*_2_, ···, *U*_ℛ_} being the ground truth partition and *V* = {*V*_1_, *V*_2_, ···, *V*_𝒞_} being the predicted clustering of Spartan, of *N* spots. Let *n*_*ij*_ denote the number of spots simultaneously in cluster *U*_*i*_ and *V*_*j*_.

Let *a*_*i*_ = Σ_*j*_ *n*_*ij*_ and *bj* = Σ_*i*_ *n*_*ij*_ represent the cluster sizes in *U* and *V* respectively. The ARI is defined as:

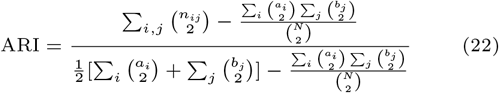

The numerator measures the excess agreement between the two clusterings relative to that expected by random chance, while the denominator normalizes the score so that random labelings yield an expected value of zero and identical clusterings yield a value of one. The ARI is invariant to label permutations and robust to differences in cluster sizes. In this work, we use the ARI as implemented in scikit-learn [45].

#### Normalized Mutual Information

The Normalized Mutual Information (NMI) quantifies the mutual dependence between a predicted clustering and a ground truth labeling. Given two clustering assignments: *U* = {*U*_1_, *U*_2_, ···, *U*_ℛ_} being the ground truth partition and *V* = {*V*_1_, *V*_2_, ···, *V*_𝒞_} being the predicted clustering of Spartan, of *N* spots.

Let *n*_*ij*_ = |*U*_*i*_ ∩ *V*_*j*_ |, *a*_*i*_ = Σ_*j*_ *n*_*ij*_ and *b*_*j*_ = Σ_*i*_ *n*_*ij*_. Define the probability distributions 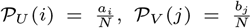, and joint-probability 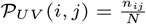. The mutual information is:

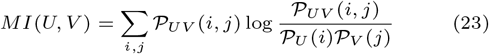

The entropies of the two clustering assignments are:

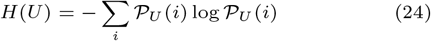

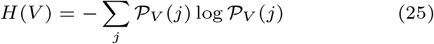

The NMI is defined as:

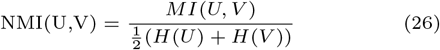

The NMI(U,V) ∈ [0,1]. A value of 1 indicates perfect agreement, and a value near 0 indicates essentially independent assignments. In our experiments, we compute NMI using scikit-learn [45].

#### Homogeneity and Completeness

Homogeneity (HOM) and Completeness (COM) are information-theoretic metrics that assess the quality of clustering with respect to known true cluster labels. Homogeneity (HOM) measures whether each predicted cluster contains only spots of a single ground truth cluster, while completeness assesses whether all spots of a given ground truth cluster are assigned to the same predicted cluster.

Formally, given ground truth labels *U* = {*U*_1_, *U*_2_, ···, *U*_ℛ_} and predicted clusters *V* = {*V*_1_, *V*_2_, ···, *V*_𝒞_}, the conditional entropies are defined as:

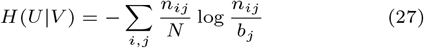

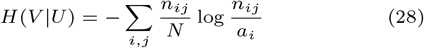

where *n*_*ij*_ = |*U*_*i*_ ∩ *V*_*j*_ |, *a*_*i*_ = Σ_*j*_ *n*_*ij*_, *b*_*j*_ = Σ_*i*_ *n*_*ij*_, and *N* is the total number of spots.

The entropy of true and predicted labels are 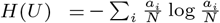 and 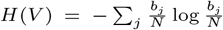. HOM and COM are then defined as:

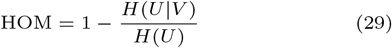

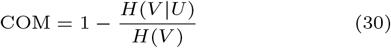

We compute both scores using scikit-learn’s implementation [45].

#### R score

While individual clustering metrics such as NMI, homogeneity (HOM), and completeness (COM) each capture distinct aspects of clustering quality, they can vary in sensitivity depending on cluster purity or fragmentation. To obtain a balanced, single-value summary of clustering performance across these complementary information-theoretic measures, we define the R score [46] as their arithmetic mean:

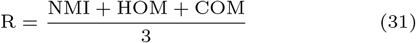

A higher R score indicates simultaneously high mutual information, cluster purity, and label completeness, thus providing a robust overall indicator of alignment between predicted and ground truth structures.

#### Normalized Local Spatial Activation Score (nLSAS)

The normalized Local Spatial Activation Score (nLSAS) quantifies the distribution of LSA-derived activation mass across clusters. It measures how activation mass derived from the LSA graph is distributed among cluster-specific regions, thereby reflecting structural consistency and fragmentation characteristics of a clustering solution.

Let *V* = {*V*_1_, *V*_2_, ···, *V*_𝒞_} denote the set of clusters, where 𝒞 is the total number of clusters. For each spot *i* ∈ {1, 2, ···, *N*}, we define its LSA as:

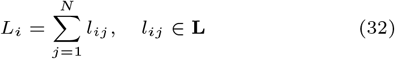

where *l*_*ij*_ represents the LSA of spot *i* with respect to spot *j*, derived from the LSA matrix **L**. Since **L** is asymmetric, *L*_*i*_ corresponds to the row-sum (outgoing activation mass) of spot *i*.

For each cluster *V*_*k*_ ∈ *V*, the cluster-level activation mass is computed as:

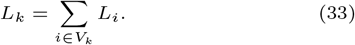

We then determine the maximum and minimum activation mass across all clusters:

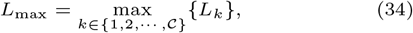

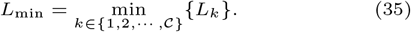

To ensure comparability across clusters and datasets, we normalize the cluster activation masses to the range [0, 1] as:

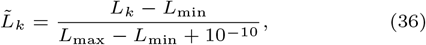

where a small constant (10^−10^) avoids division by zero when *L*_max_ = *L*_min_ and ensures numerical stability.

Finally, the normalized Local Spatial Activation Score (nLSAS) is defined as the average normalized activation across all clusters:

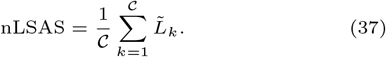

nLSAS summarizes how LSA-derived activation mass is distributed across clusters and provides a dataset-normalized measure of structural heterogeneity. nLSAS was used as an unsupervised stability prior during *α*-selection.

### Datasets

To assess Spartan’s performance in spatial domain identification and spatially variable gene (SVG) detection, we have evaluated Spartan on a diverse group of SRT datasets, coming from many different technologies and tissue types. Seven of those datasets have been used for spatial domain identification and four of them for SVG detection. Across the manuscript, we use the term sample to describe a single slice or a tissue section of the SRT. We use dataset to describe a set of SRT samples. The description of each dataset are given below:

#### Stereo-seq

The Stereo-seq dataset is sequencing-based SRT dataset obtained by high-resolution technology that spans the full transcriptome. The dataset consists of nine samples from mouse embryo depicting organogenesis [11]. The samples of the dataset [47] have the following number of spots: 5,913 (E9.5E1S1), 5,292 (E9.5E2S1), 4,356 (E9.5E2S2), 5,059 (E9.5E2S3), 5,797 (E9.5E2S4), 18,408 (E10.5E1S1), 18,647 (E10.5E1S2), 18,670 (E10.5E1S3), and 8,494 (E10.5E2S1). The corresponding number of genes: 25,568 (E9.5E1S1), 23,756 (E9.5E2S1), 24,107 (E9.5E2S2), 24,238 (E9.5E2S3), 23,398 (E9.5E2S4), 25,201 (E10.5E1S1), 25,244 (E10.5E1S2), 25,647 (E10.5E1S3), and 22,385 (E10.5E2S1).

#### BaristaSeq

The BaristaSeq dataset is a SRT dataset obtained by barcoded BarisTaSeq technology [48] which allows faster image acquisition. The dataset [47] consists of three samples from the mouse primary cortex. The number of cells in the samples: 1,525 (Slice 1), 2,042 (Slice 2), and 1690 (Slice 3). The size of the gene panel is 79.

#### MERFISH

The MERFISH dataset is SRT dataset obtained from imaging-based MERFISH technology. The dataset represents the preoptic region of the mouse hypothalamus [49]. The dataset [47] consists of five samples with number of cells: 5,488 (Bregma-0.04), 5,557 (Bregma-0.09), 5,926 (Bregma-0.14), 5,803 (Bregma-0.19), and 5,543 (Bregma-0.24). The gene panel size is 155. The regional annotations of the dataset is published in [12].

#### osmFISH

The osmFISH dataset is an imaging-based SRT dataset obtained by nonbarcoded osmFISH technology [50]. The dataset [47] consists of one sample representing the mouse somatosensory cortex. The gene panel is made up of 33 marker genes. The sample consists of 4,839 cells. The regional annotations of the dataset is published in [51].

#### STARmap

The STARmap dataset is an imaging-based SRT dataset obtained by in-situ RNA-sequencing STARmap technology [52]. The dataset [47] consists of three samples from the high cognitive region of the brain, the medial prefrontal cortex. The samples consist of the following number of cells: 1,049 (BZ5), 1,053 (BZ9), and 1,088 (BZ14). The gene panel size is 166. All samples are annotated to four distinct layer structures [12].

#### STARmap*

The STARmap* is imaging-based SRT dataset, version of the STARmap dataset but has 1020 genes. The dataset [47] consist of one sample with 1207 cells. The dataset represents the visual cortex with regions enriched with excitatory, non-neuronal, and inhibitory cell types, expertly annotated [52].

#### Vizgen MERFISH

The Vizgen MERFISH is imaging-based SRT dataset obtained by the MERFISH technology. The dataset contains a gene panel of 483 genes [53]. This includes canonical cell type markers, nonsensory G-protein coupled receptors (GPCRs), and Receptor Tyrosine Kinases (RTKs). Cell type markers are mapped using Vizgen’s own pipeline. The dataset consists of four samples derived from the coronal section of the mouse brain. The number of cells in the samples are:78,329 (S1R1), 88884 (S1R2), 83,546 (S2R1), and 85,958 (S2R3). The samples are regionally annotated by [9].

#### 10x Visium

The 10x Visium is spot sequencing-based SRT dataset. The dataset consists of 12 samples from the human dorsolateral pre-frontal cortex (DLPFC) region [54]. Each sample spans six neuronal layers and the white matter. The number of spots in the samples [47]: 4,226 (151507), 4,384 (151508), 4,789 (151509), 4,634 (151510), 3,661 (151669), 3,498 (151670), 4,110 (151671), 4,015 (151672), 3,639 (151673), 3,673 (151674), 3,592 (151675), and 3,460 (151676). The total number of genes in the dataset is 33,538. The dataset is expertly annotated with six neuronal layers and the white matter [55].

#### Visium HD dataset

High-resolution spatial transcriptomic profiling was conducted on a 10-week post-conception (wpc) human fetal gastroesophageal specimen using the Visium HD Spatial Gene Expression platform (10x Genomics). Developmental staging was determined by foot length (8 mm) in combination with ultra-sonographic assessment.

##### Human tissue collection and processing

Human embryonic material was obtained from elective terminations of pregnancy under ethical approval from the North West Research Ethics Committee (23/NW/0039), with written informed consent from participants and in accordance with the Human Tissue Authority Codes of Practice.

The esophagus, stomach, and proximal duodenum were dissected and fixed within 1 h post-collection in 4% paraformaldehyde under RNase-free conditions. Tissue was processed and paraffin-embedded using standard protocols. Sagittal sections (5 μm thickness) were obtained, with haematoxylin and eosin (H&E) staining performed on every eighth section for anatomical reference.

Sections selected for spatial transcriptomic profiling were processed following the Visium HD FFPE Tissue Preparation Handbook (CG000684) and the Visium HD Spatial Gene Expression Reagent Kits User Guide (CG000685). Sections were deparaffinised, destained, de-crosslinked, and permeabilised prior to overnight probe hybridisation. Following probe ligation and washing, probe transfer was performed using the 10x Genomics CytAssist instrument. Libraries were prepared using the Visium HD Spatial Gene Expression Reagent Kit according to the manufacturer’s protocol and sequenced on an Illumina NovaSeq 6000 platform (SP flow cell, 100-cycle configuration).

##### Data preprocessing and cell-level binning

Raw sequencing data were processed using the Space Ranger pipeline (10x Genomics, version 3.1.2) with alignment to the GRCh38 reference genome (release 2020-A).

To achieve cell-level resolution, histological images were analysed using QuPath (ersion 0.5.1-arm64). Nuclei were detected based on intensity and morphological parameters derived from H&E staining, followed by watershed-based expansion of nuclear boundaries to approximate whole-cell areas. Segmentation parameters included intensity threshold of 0.1, background radius 5 *μ*m, and a minimum cell area of 10 *μ*m^2^.

Custom cell-specific bins were generated by assigning Visium HD barcodes to segmented cell polygons based on centroid overlap between barcode spatial coordinates and cell boundaries. For each segmented cell, gene-wise UMI counts from overlapping barcodes were aggregated to generate a gene-by-cell count matrix.

Cells were filtered to retain those with ≥20 UMIs and a maximum cell area of 2000 *μ*m^2^.

The final preprocessed dataset comprised 84,732 segmented cells and 18,085 detected genes following quality control filtering.

## Data availability

The authors analyzed eight publicly available and one in-house SRT datasets. The data were acquired from the following websites or accession numbers: (1) Stereo-seq, BaristaSeq, MERFISH, osmFISH, STARmap, STARmap*, 10x Visium datasets with regional annotations were downloaded from http://sdmbench.drai.cn/; (2) Vizgen MERFISH dataset was downloaded from https://console.cloud.google.com/storage/browser/public-datasets-vizgen-merfish;tab=objects?prefix=&forceOnObjectsSortingFiltering=false; (3) The Visium HD dataset generated in this study is available in ArrayExpress under accession number E-MTAB-16671.

## Code availability

An open-source implementation of Spartan can be downloaded from https://github.com/MohammadFaizIqbalFaiz/spartan-st, where code and scripts for reproducing the results and figures are provided.

## Acknowledgments

The authors thank all the contributors of the public data which was used in this study. Mohammad Faiz Iqbal Faiz was funded by BBSRC DTP award (BB/T008725/1).

