## Supplementary Figures for "Spartan: activation-aware framework for spatial domain and variable gene discovery"

#### Supplementary Methods and Benchmarking

This supplementary document presents additional analyses supporting the spatial domain identification capabilities of Spartan across multiple spatial transcriptomics technologies and datasets. It includes supplementary methods, SVG validation, runtime and scalability analysis, parameter sensitivity, $\alpha$-selection strategies, and comprehensive benchmarking on SDMBench datasets as well as comparisons against recently proposed methods.

##### 0.1 Supplementary methods

##### 0.2 Validation of Local Spatial Activation (LSA) graph structure

To evaluate whether the Local Spatial Activation (LSA) graph captures biologically meaningful spatial structure, we analyzed the distribution of LSA weights with respect to ground truth spatial domains. The analysis was performed using the spatial neighbor graph of each dataset together with the LSA matrix $\mathbf{L}$ produced by Spartan.

**Edge-level analysis of LSA weights** For each spatial neighbor pair $(i, j)$ in the spatial adjacency matrix $\mathbf{S}$, the corresponding LSA edge weight $l_{ij}$ was obtained from the LSA adjacency matrix $\mathbf{L}$. Using the ground truth labels, edges were categorized as either intra-domain edges (spots belonging to the same domain) or inter-domain edges (spots belonging to different domains).

The distributions of $l_{ij}$ values for intra-domain and inter-domain edges were compared to evaluate whether LSA preferentially assigns stronger weights to relationships within spatial domains. Median LSA weights were computed for both categories, and a Mann–Whitney U test was used to assess whether the distributions differed significantly.

To quantify effect size, the rank-biserial correlation (RBC) was computed based on the Mann–Whitney U statistic. The RBC provides a measure of the degree to which LSA weights preferentially favor intra-domain edges over inter-domain edges.

**Discriminative analysis using ROC curves** To evaluate whether LSA weights provide discriminative information about spatial organization, receiver operating characteristic (ROC) analysis was performed.

For edge-level analysis, the task was to distinguish intra-domain edges from inter-domain edges using the LSA edge weight $l_{ij}$ as the predictor variable. The area under the ROC curve (AUROC) was computed to quantify discriminative performance.

For boundary analysis, local LSA magnitudes were used to distinguish boundary spots from interior spots, and AUROC values were similarly computed.

**Permutation-based validation** To confirm that the observed differences in LSA distributions were not attributable to random spatial arrangements, a permutation analysis was performed. Ground truth spatial domain labels were randomly permuted while preserving the spatial graph structure, and the difference between median intra-domain and inter-domain LSA weights was recomputed for each permutation. Repeating this procedure generated a null distribution of median differences under random spatial labeling.

The observed median difference was then compared against the null distribution to compute an empirical permutation p-value, thereby assessing whether the observed LSA-domain association exceeded that expected under spatial randomness.

##### 0.2.1 Validation of SAQ-based spatially variable genes

For validating the SVGs identified by Spartan, we created a list of candidate genes representing a biologically relevant background set. First, we selected the top 1000 highly variable genes (HVGs), ranked by the metric *dispersions_norm*, denoted as $U_{1000}$, using `Scanpy`'s *pp.highly_variable_genes* function. We used the *seurat* flavor for identifying highly variable genes. Genes considered undesirable or potentially confounding (e.g., mitochondrial genes, ribosomal genes, and other non-informative transcripts) were removed from $U_{1000}$. The remaining genes were further filtered to retain genes detected in at least 0.25% of the cells, ensuring sufficient spatial signal for reliable spatial statistics. Finally, the curated list, $U_{\text{final}}$, contained 195 candidate genes.

Next, we applied Spartan to compute SAQ scores for all genes in the gene panel (18,085 genes). The top 100 genes according to their SAQ scores were selected and denote this set of genes as $G_{SAQ}$. We then constructed two gene sets for validation. The first set, $G_{top} = G_{SAQ} \cap U_{\text{final}}$, contains 33 genes that are both highly ranked by SAQ and belong to the candidate background pool. The second set, $G_{bg} = U_{\text{final}} \setminus G_{SAQ}$, contains the remaining 162 background genes from the candidate pool. This design ensures that comparisons are performed among genes with comparable expression variability, reducing potential confounding effects due to gene expression dispersion.

We computed Moran's I statistic for all genes in the gene panel using `Squidpy`'s *gr.spatial_autocorr* function based on the spatial neighbor graph of the Visium HD dataset. Moran's I values of genes in the lists $G_{top}$ and $G_{bg}$ were then compared. To evaluate whether SAQ-ranked genes exhibit stronger spatial structure, we performed a non-parametric Mann–Whitney U test to determine whether Moran's I values of genes in $G_{top}$ are systematically larger than those of genes in $G_{bg}$.

Finally, to test whether genes identified by SAQ are enriched for statistically significant spatial autocorrelation, we evaluated enrichment of Moran-significant genes within $G_{SAQ}$. Genes were considered Moran-significant if they satisfied a false discovery rate threshold of FDR $< 0.05$ during Moran's I computation. Enrichment of Moran-significant genes in $G_{SAQ}$ relative to the background pool $U_{\text{final}}$ was assessed using Fisher's exact test. P-values were computed for both the Mann–Whitney U test and Fisher's exact test, and the odds ratio was reported for the enrichment analysis.

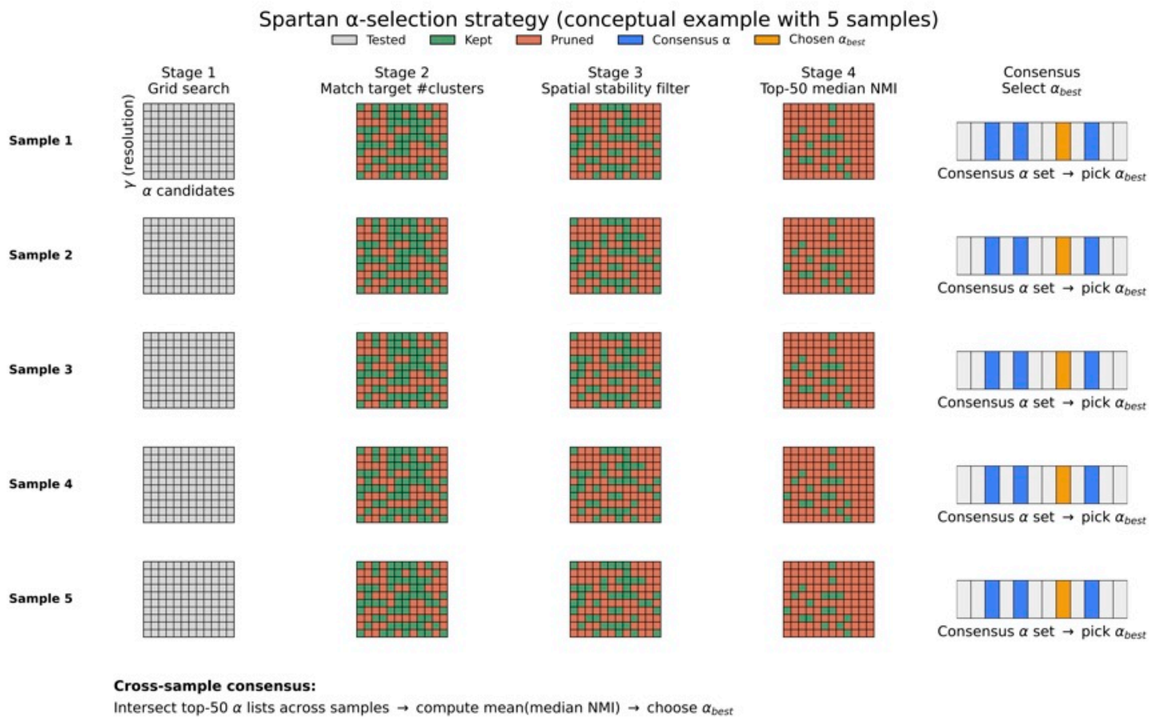


**Supplementary Figure 1.** Spartan $\alpha$-selection strategy based on multi-stage pruning and cross-sample consensus. Conceptual illustration of Spartan's $\alpha$-selection workflow using five representative samples. Stage 1 (Grid search): A grid of candidate $\alpha$ values is evaluated for each sample across a range of clustering resolutions. Stage 2 (Target cluster matching): Candidate $\alpha$ values that recover the target number of spatial domains are retained. Stage 3 (Spatial stability filtering): Candidates are further filtered based on spatial stability criteria, removing $\alpha$ values that produce unstable or fragmented spatial domains. Stage 4 (Performance ranking): Remaining candidates are ranked by median Normalized Mutual Information (NMI), and the top-performing $\alpha$ values are retained for each sample. Consensus selection: A consensus set of $\alpha$ values is constructed across samples, and a single dataset-level $\alpha_{\text{best}}$ is selected. Colors indicate tested, retained, and pruned $\alpha$ values at each stage. This strategy enables robust $\alpha$ selection by integrating biological priors, spatial stability, and clustering performance while avoiding exhaustive parameter tuning on individual samples.

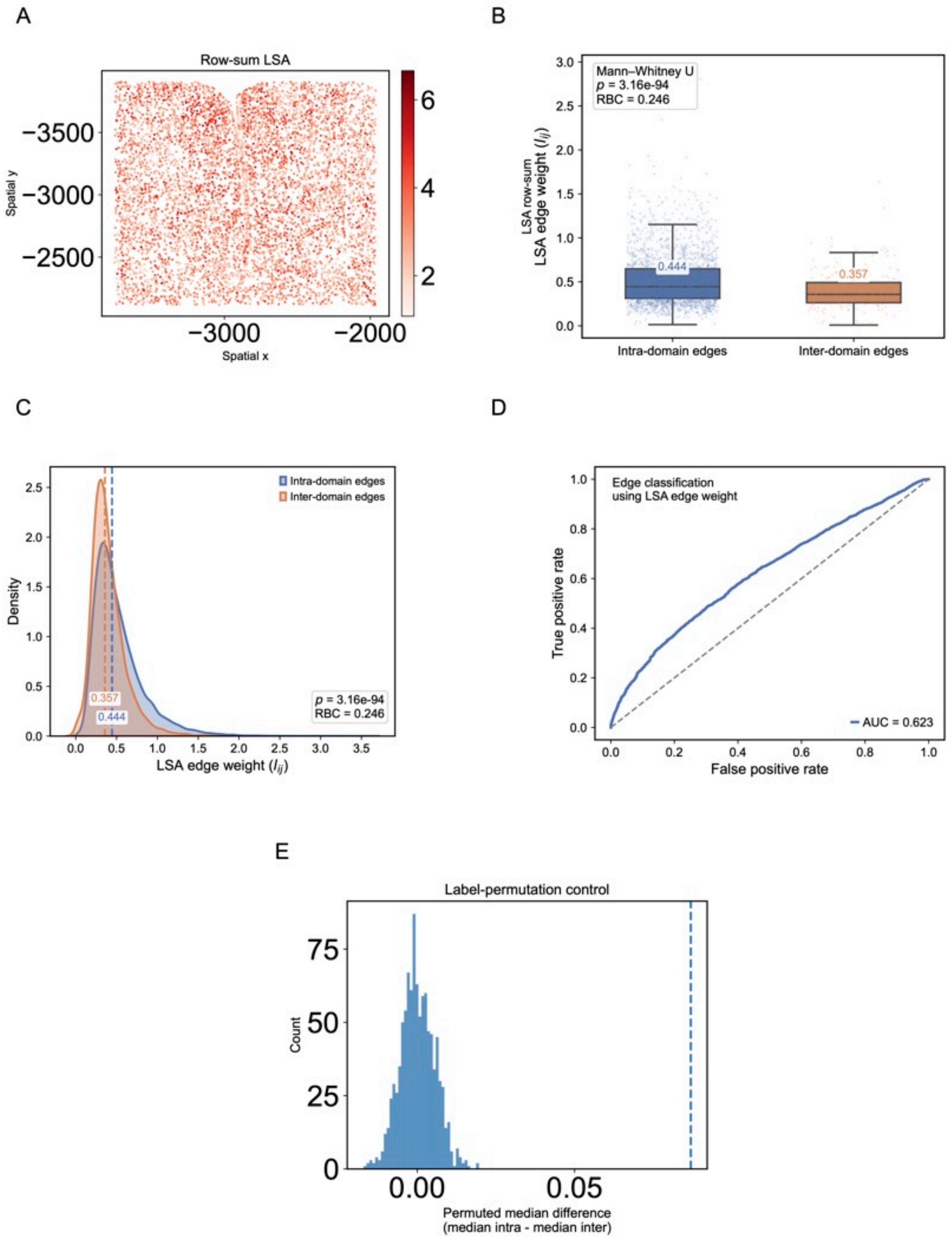


**Supplementary Figure 2.** Validation of the Local Spatial Activation (LSA) graph on the MERFISH 0.24 sample. (A) Spatial distribution of row-sum LSA values, showing spatially coherent patterns across the tissue. (B) Distribution of LSA edge weights ($l_{ij}$) for spatial neighbor pairs within the same ground-truth domain (intra-domain) and across different domains (inter-domain). Intra-domain edges exhibit higher LSA weights, indicating preferential strengthening of connections within spatial domains (median intra-domain: 0.444; inter-domain: 0.357; Mann–Whitney U test $p = 3.16 \times 10^{-94}$; RBC = 0.246). (C) Density distributions of LSA edge weights for intra- and inter-domain edges, illustrating clear separation between the two categories. (D) Receiver operating characteristic (ROC) analysis using LSA edge weights to distinguish intra-domain from inter-domain edges. The area under the curve (AUROC = 0.623) indicates that LSA weights provide discriminative information about domain structure. (E) Permutation-based control showing the null distribution of median intra- versus inter-domain LSA differences under random label assignments. The observed difference (dashed line) lies far from the null distribution, indicating that the LSA-domain association is not explained by random spatial organization. Together, these results demonstrate that the LSA graph captures biologically meaningful spatial organization and preferentially reinforces relationships within annotated spatial domains.

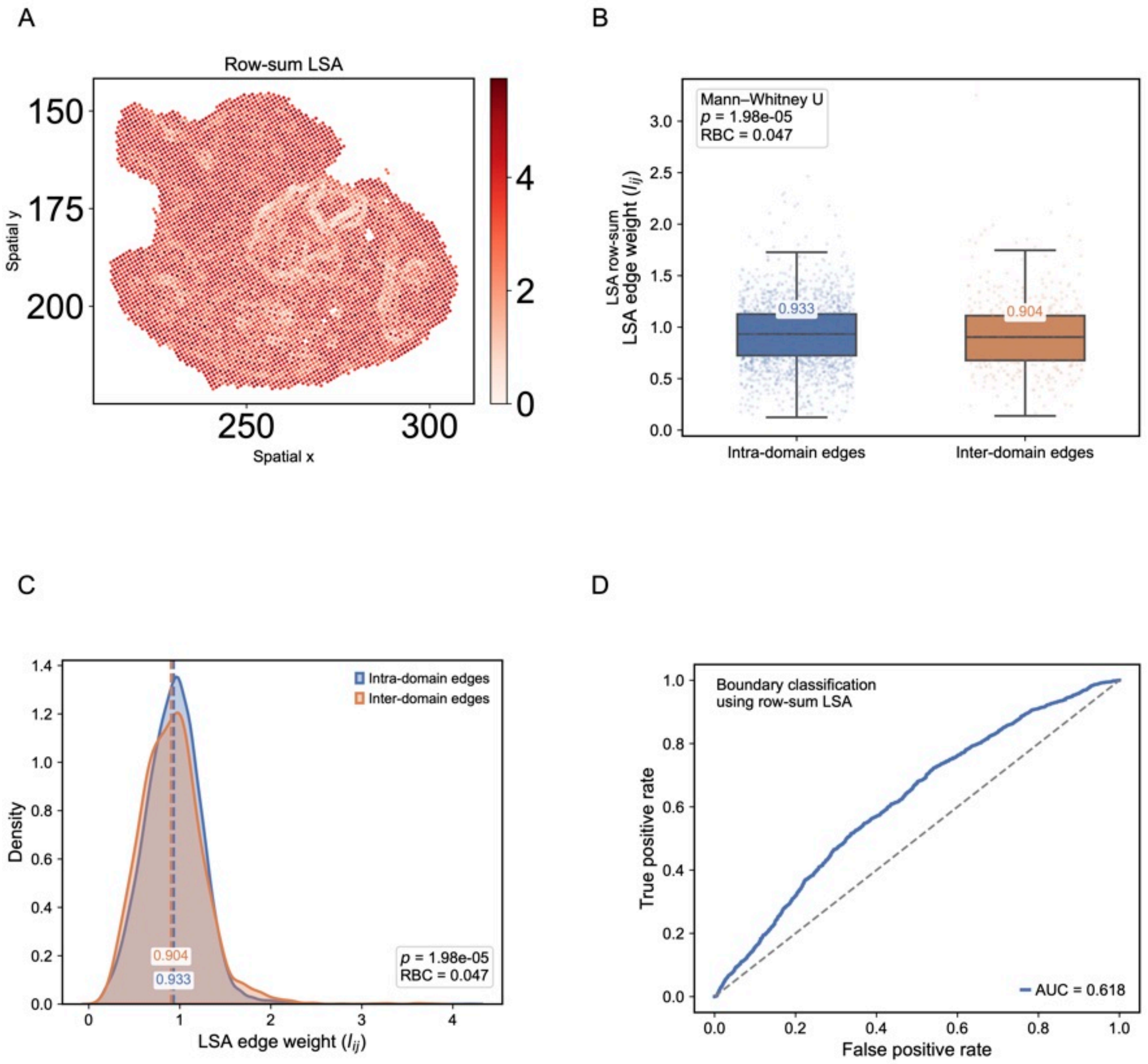


**Supplementary Figure 3.** Validation of the Local Spatial Activation (LSA) graph on the Stereo-seq E2S3 sample. (A) Spatial distribution of local LSA magnitude (row-sum of $l_{ij}$), showing coherent spatial patterns across the tissue. (B) Distribution of LSA edge weights ($l_{ij}$) for spatial neighbor pairs within the same ground-truth domain (intra-domain) and across different domains (inter-domain). Intra-domain edges exhibit slightly higher LSA weights, consistent with within-domain spatial coherence (median intra-domain: 0.933; inter-domain: 0.904; Mann–Whitney U test $p = 1.98 \times 10^{-5}$; RBC = 0.047). (C) Kernel density estimates of LSA edge weights for intra- and inter-domain edges, showing modest but consistent separation between the two distributions. (D) Receiver operating characteristic (ROC) analysis using local LSA magnitudes (row-sum LSA) to distinguish boundary from interior spots. The area under the curve (AUROC = 0.618) indicates that LSA captures informative signals at spatial domain boundaries. These results indicate that the LSA graph encodes spatial activation patterns that reflect both within-domain coherence and transitions across domain boundaries.

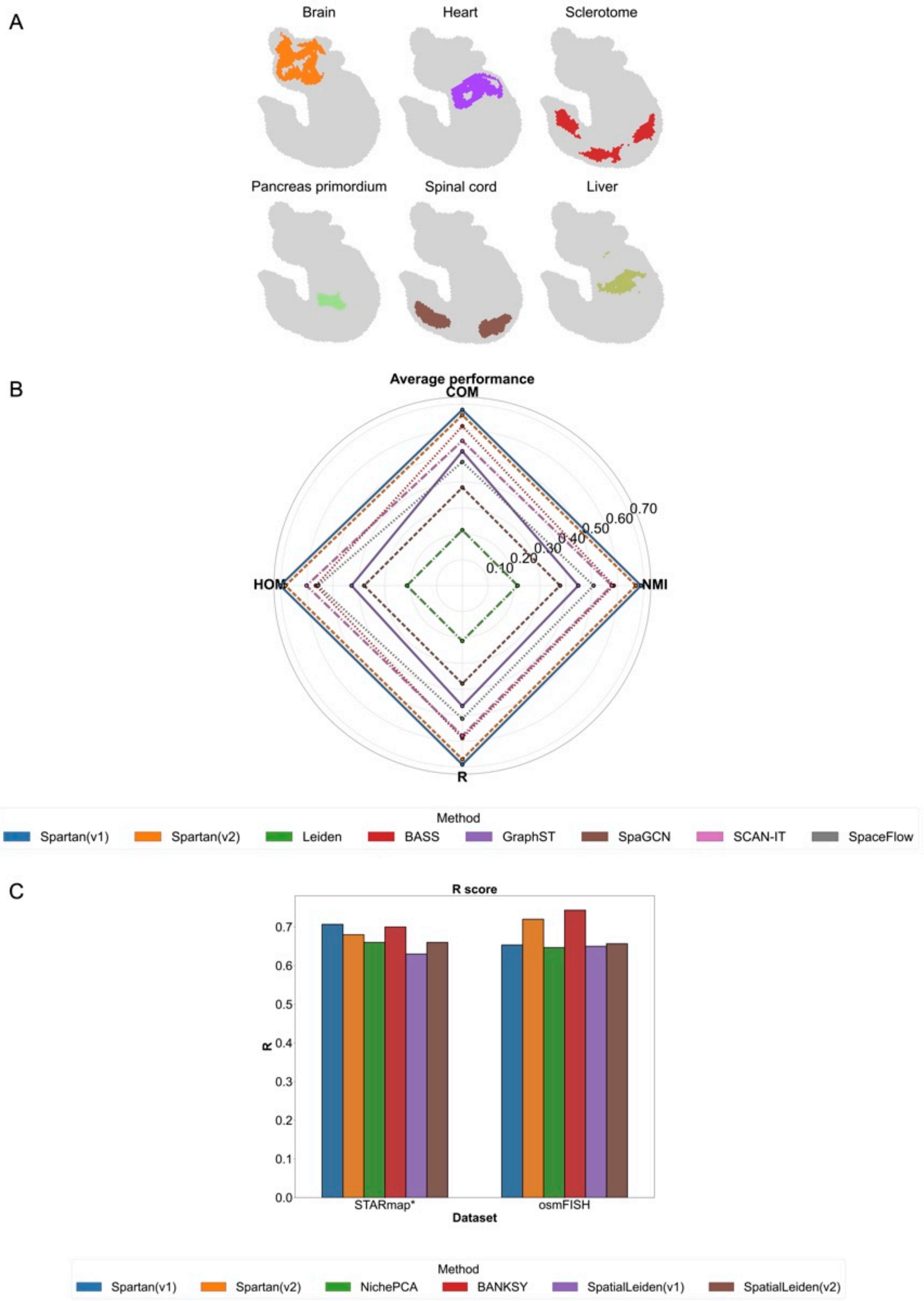


**Supplementary Figure 4.** (A) Spatial domains identified on the Stereo-seq E9.5E2S2 embryo sample, resolving anatomically coherent tissue compartments, including brain, heart, sclerotome, liver, pancreas primordium, and spinal cord. (B) Radar chart summarizing average performance across multiple clustering metrics (COM, HOM, NMI, and R score), benchmarked against established methods within the SDMBench framework. Larger polygons indicate stronger and more balanced multi-metric recovery. (C) Single-sample performance comparison using the R score on representative datasets.

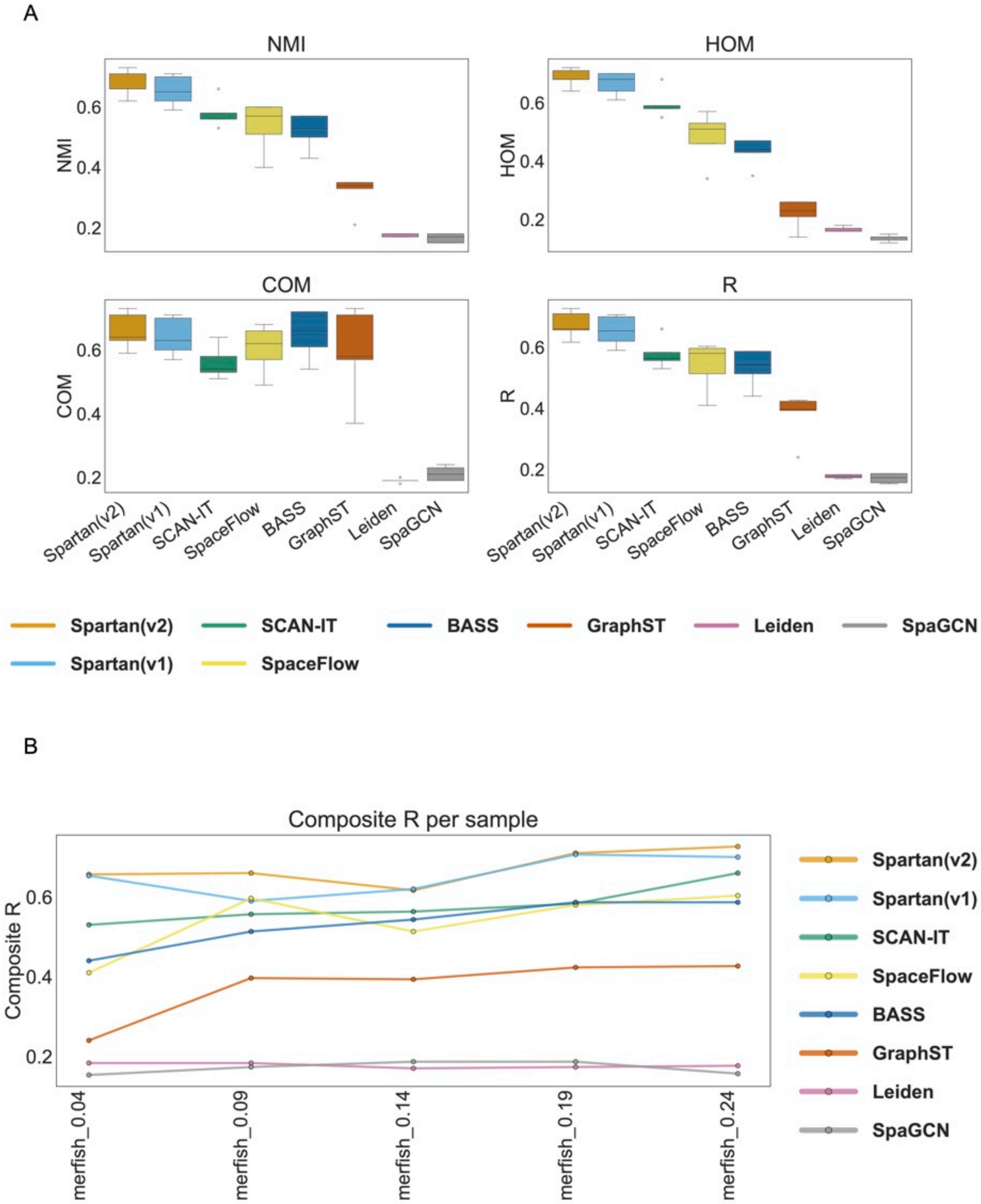


**Supplementary Figure 5.** Multi-metric benchmarking of spatial domain identification methods on the MERFISH dataset from SDMBench. (A) Distribution of clustering performance metrics across five MERFISH samples from the SDMBench benchmark for all evaluated methods. Boxplots summarize per-sample values of normalized mutual information (NMI), homogeneity (HOM), completeness (COM), and the composite R score, highlighting both central tendency and variability across samples. (B) Composite R score per sample for the MERFISH dataset across methods, illustrating the consistency of overall performance across individual samples within the SDMBench benchmark.

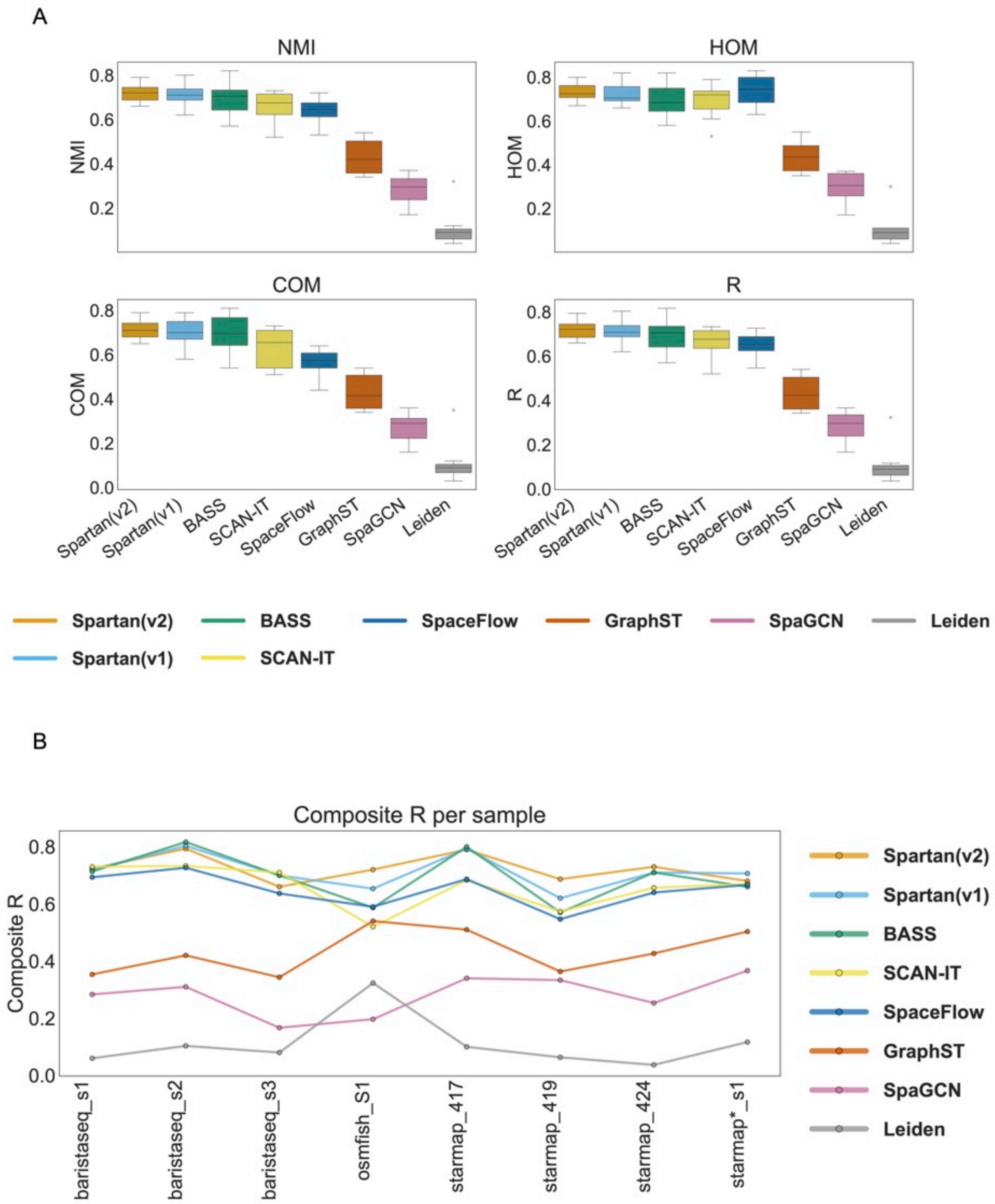


**Supplementary Figure 6.** Consolidated multi-metric benchmarking on small SDMBench datasets. Benchmarking results for spatial domain identification methods on SDMBench datasets with limited sample sizes, shown in a consolidated format. The figure combines results from BaristaSeq (three samples), STARmap (three samples), osmFISH (one sample), and STARmap* (one sample). (A) Distribution of clustering performance metrics across all samples from the consolidated datasets for each method. Boxplots summarize per-sample values of normalized mutual information (NMI), homogeneity (HOM), completeness (COM), and the composite R score. (B) Composite R score per sample across the consolidated datasets, illustrating the consistency of overall performance across methods on small-sample spatial transcriptomics data.

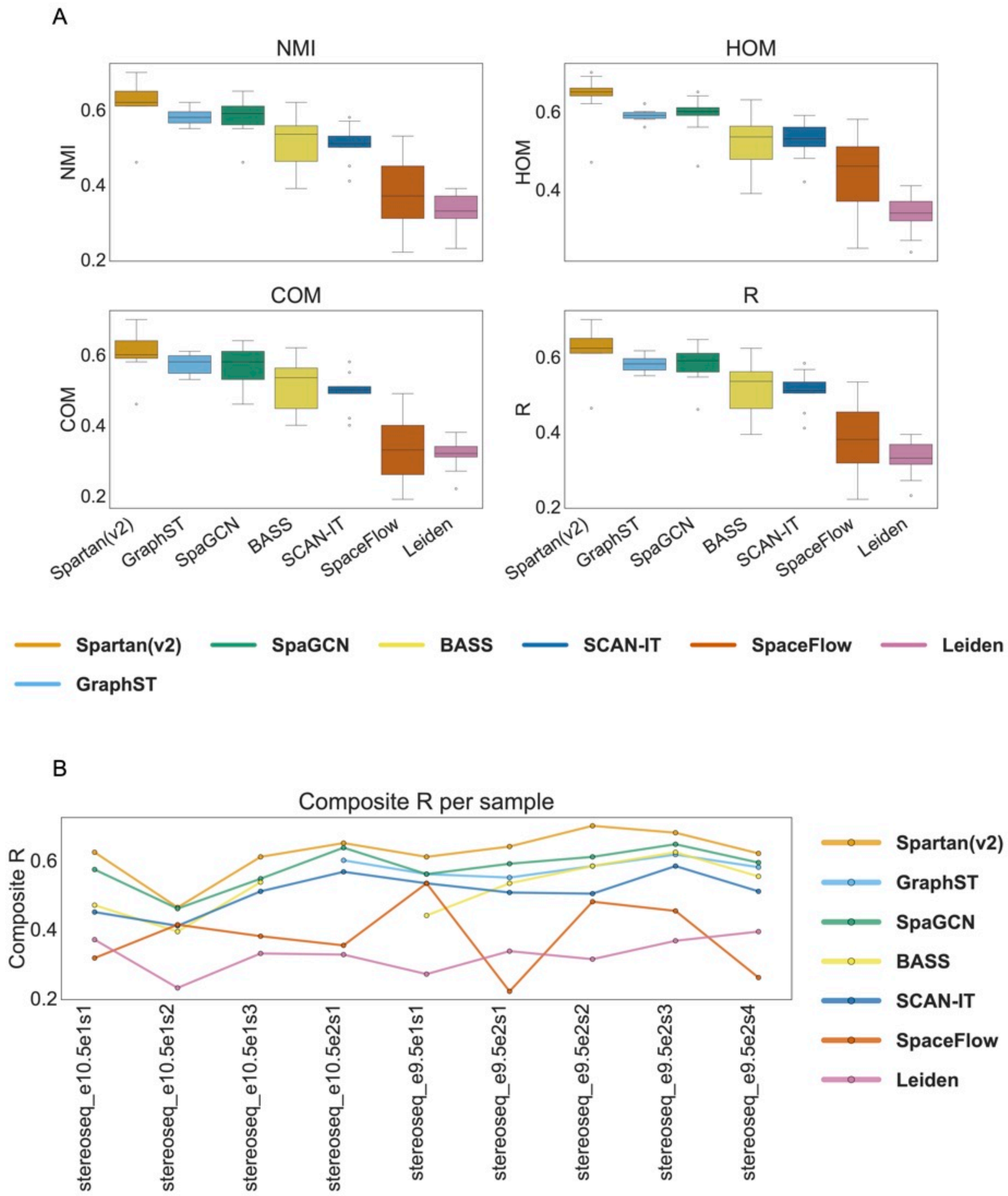


**Supplementary Figure 7.** Multi-metric benchmarking of spatial domain identification methods on the Stereo-seq dataset from SDMBench. (A) Distribution of clustering performance metrics across nine Stereo-seq samples from the SDMBench benchmark for all evaluated methods. Boxplots summarize per-sample values of normalized mutual information (NMI), homogeneity (HOM), completeness (COM), and the composite R score, highlighting both central tendency and variability across samples. (B) Composite R score per sample for the Stereo-seq dataset across methods, illustrating the consistency of overall performance across individual samples within the SDMBench benchmark.

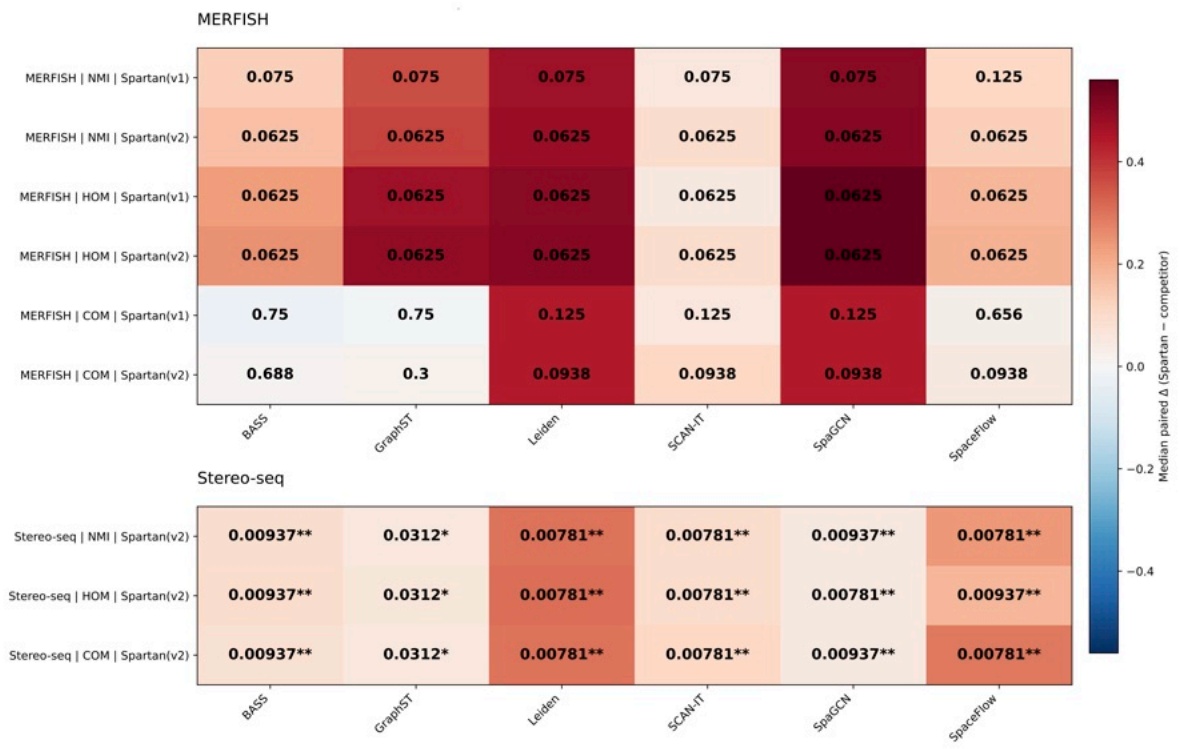


**Supplementary Figure 8.** Paired statistical comparison of Spartan and representative methods on SDM-Bench datasets. Paired two-sided Wilcoxon signed-rank tests were performed to compare both versions of Spartan (v1 and v2) against representative methods (BASS, GraphST, Leiden, SCAN-IT, SpaGCN, and SpaceFlow) across samples within MERFISH (n = 5) and Stereo-seq (n = 9) datasets. Heatmaps show median paired differences (Spartan – comparator) for NMI, HOM, and COM across samples. Cell annotations indicate Benjamini–Hochberg-adjusted P-values (false discovery rate) computed within each dataset and metric across all pairwise comparisons. Only paired samples present in both methods were included in each comparison.

#### Benchmarking against recently proposed methods

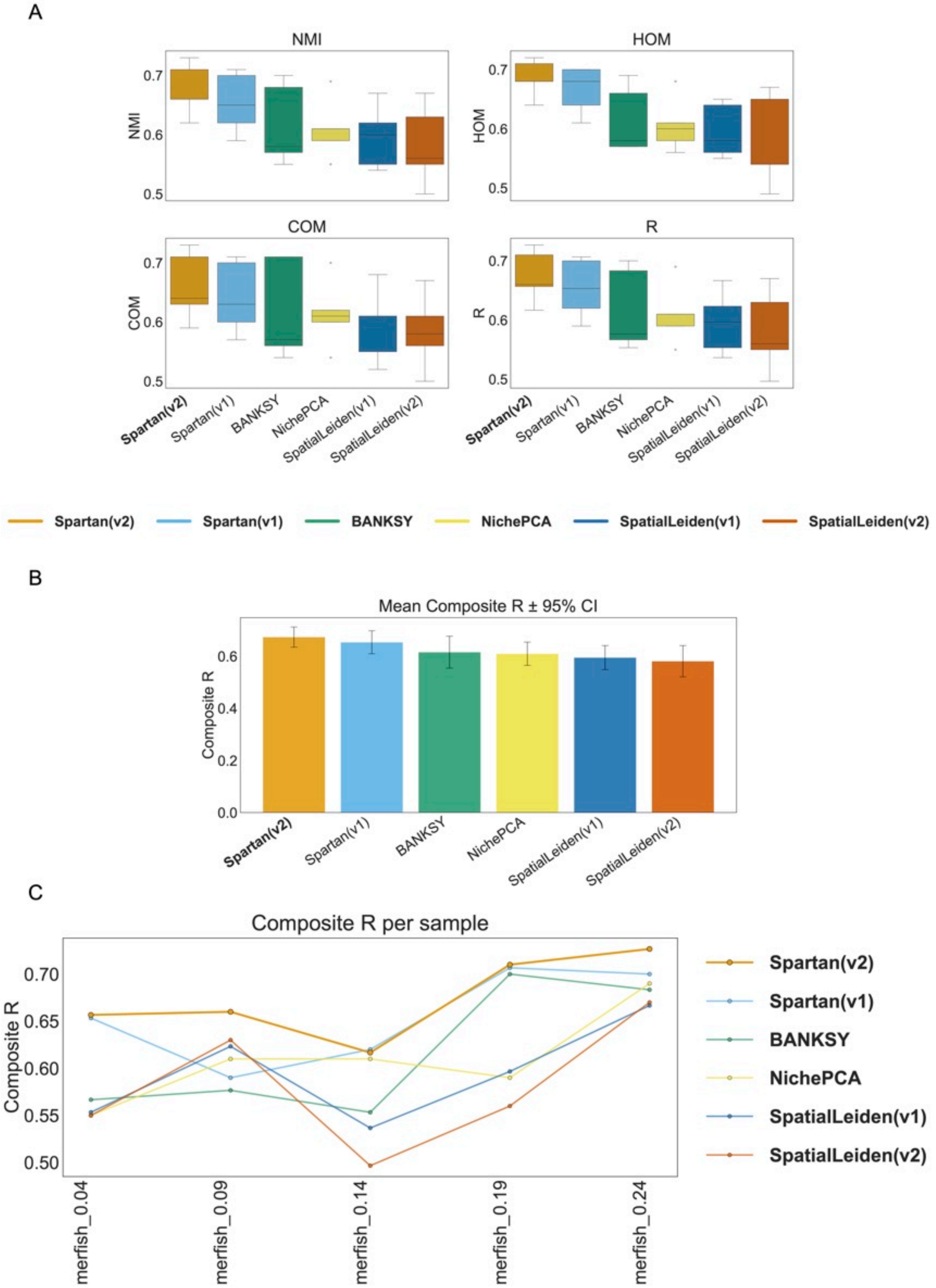


**Supplementary Figure 9.** Benchmarking against recently proposed methods on the MERFISH dataset (SDM-Bench). (A) Distribution of clustering performance metrics across MERFISH samples for Spartan and recent state-of-the-art spatial domain identification methods. Boxplots summarize per-sample values of normalized mutual information (NMI), homogeneity (HOM), completeness (COM), and the composite R score. (B) Mean composite R score with ±95% confidence intervals across samples for each method. (C) Composite R score per sample for the MERFISH dataset, illustrating consistency of performance across individual samples.

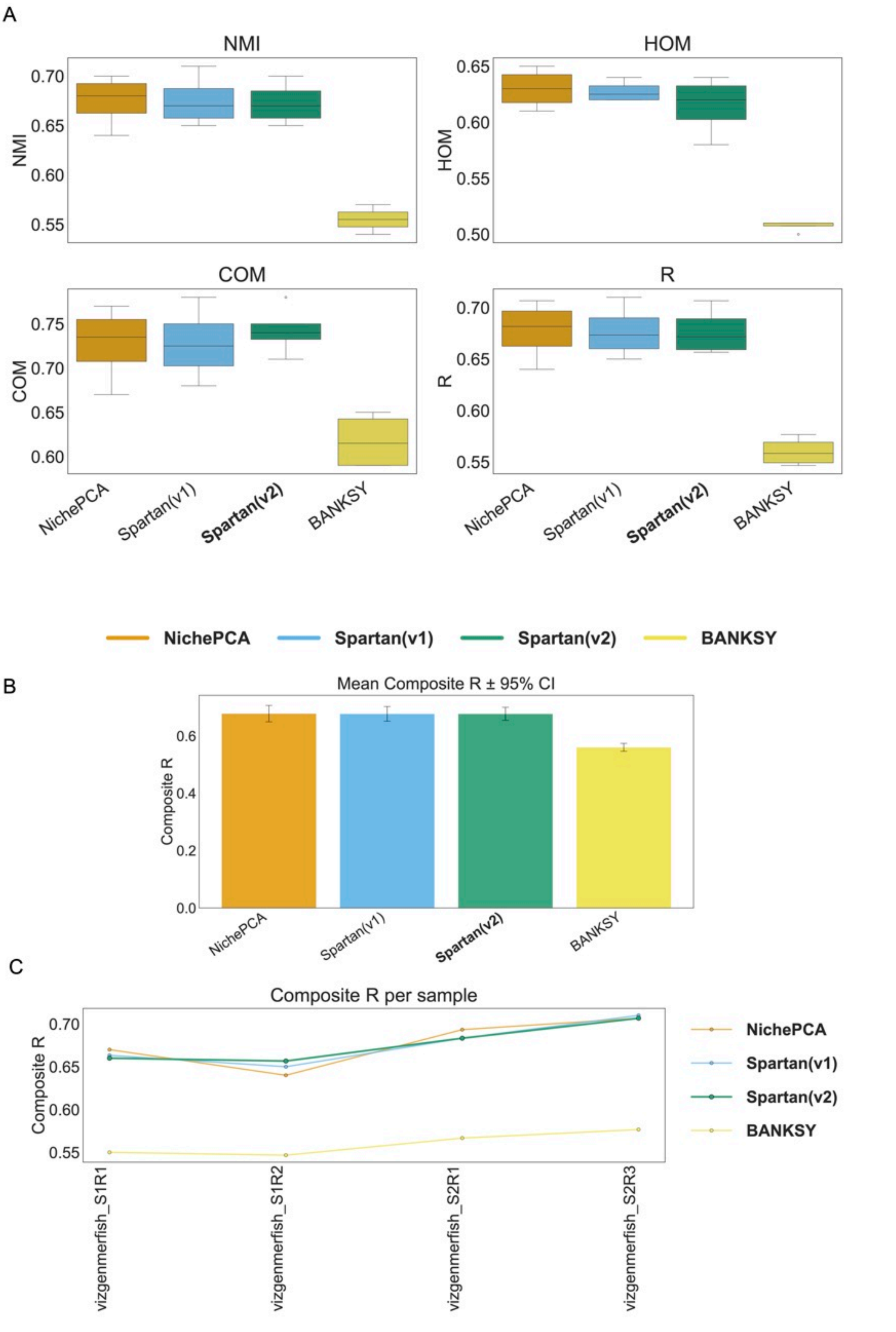


**Supplementary Figure 10.** Benchmarking against recently proposed methods on the Vizgen MERFISH dataset. (A) Distribution of clustering performance metrics across Vizgen MERFISH samples for Spartan and recent state-of-the-art methods, shown for NMI, HOM, COM, and the composite R score. (B) Mean composite R score with ±95% confidence intervals across samples for each method. (C) Composite R score per sample for the Vizgen MERFISH dataset.

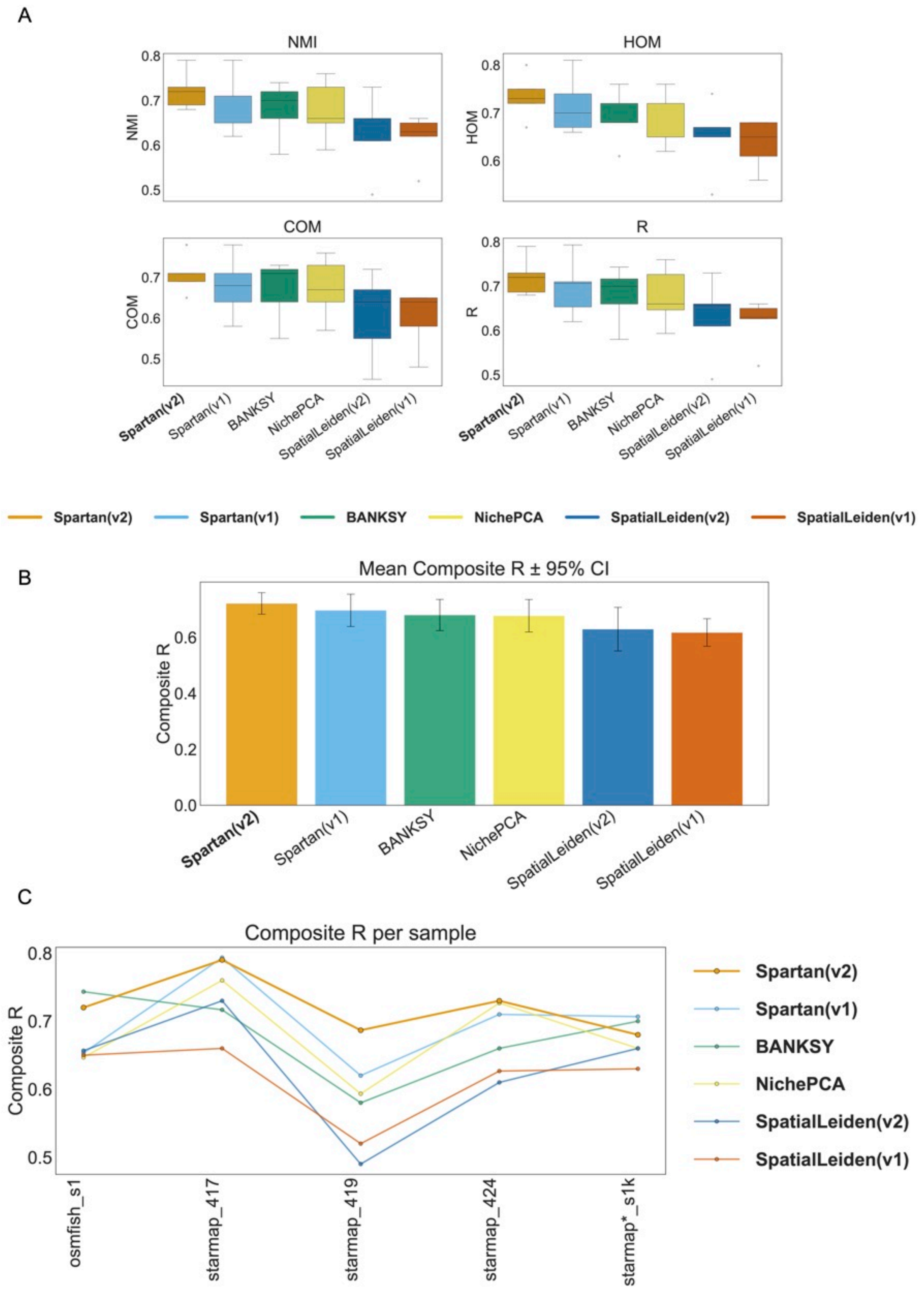


**Supplementary Figure 11.** Benchmarking against recently proposed methods on small SDMBench datasets. Benchmarking results for spatial domain identification methods on SDMBench datasets with limited sample sizes, including osmFISH (one sample), STARmap (three samples), and STARmap* (one sample), shown in a consolidated format. (A) Distribution of clustering performance metrics across all samples for each method, summarized using NMI, HOM, COM, and the composite R score. (B) Mean composite R score with ±95% confidence intervals across samples. (C) Composite R score per sample across the consolidated datasets.

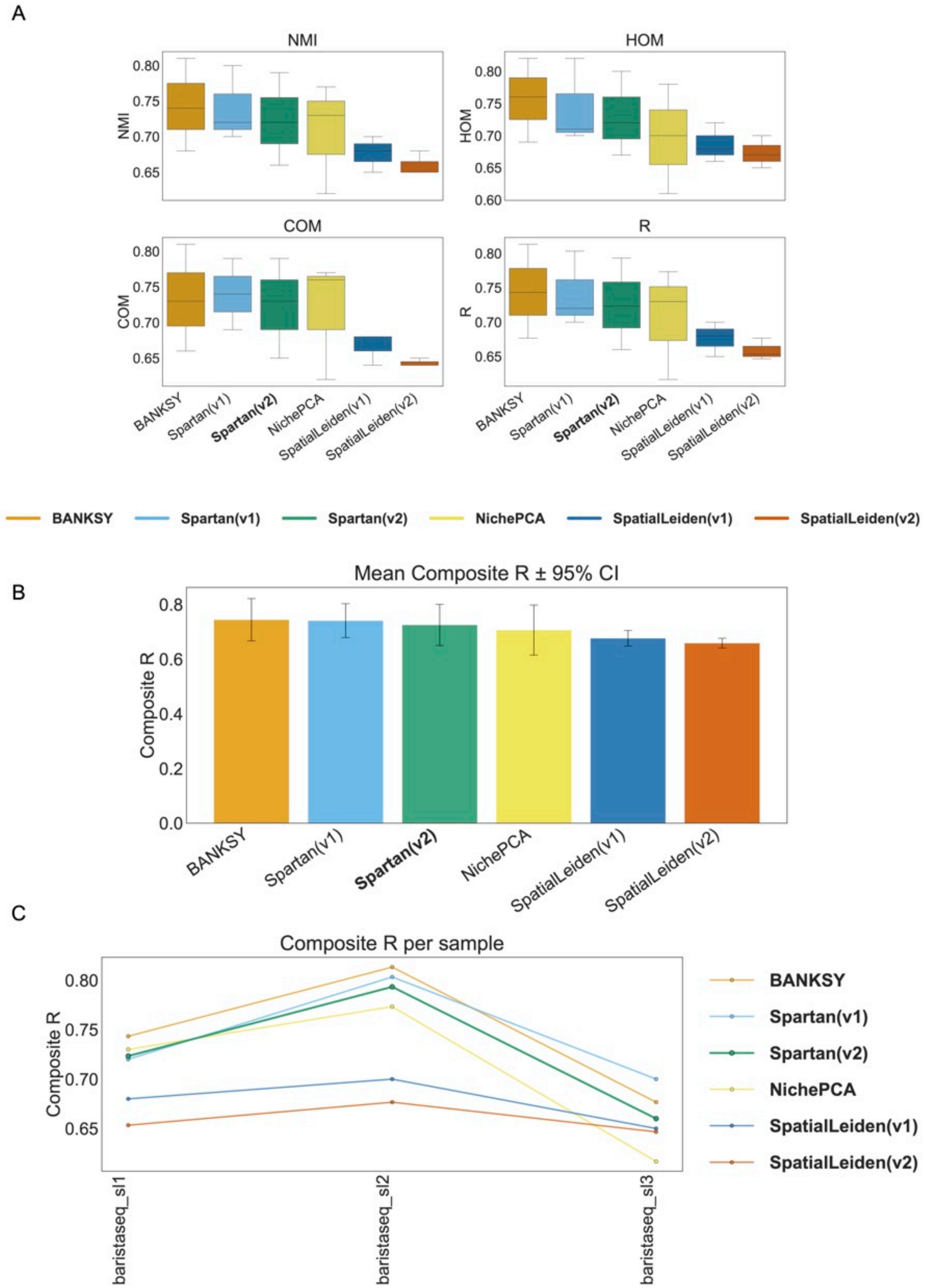


**Supplementary Figure 12.** Benchmarking against recently proposed methods on the BaristaSeq dataset (SDMBench). (A) Distribution of clustering performance metrics across BaristaSeq samples for Spartan and recent state-of-the-art methods, shown for NMI, HOM, COM, and the composite R score. (B) Mean composite R score with ±95% confidence intervals across samples for each method. (C) Composite R score per sample for the BaristaSeq dataset, illustrating consistency of performance across samples.

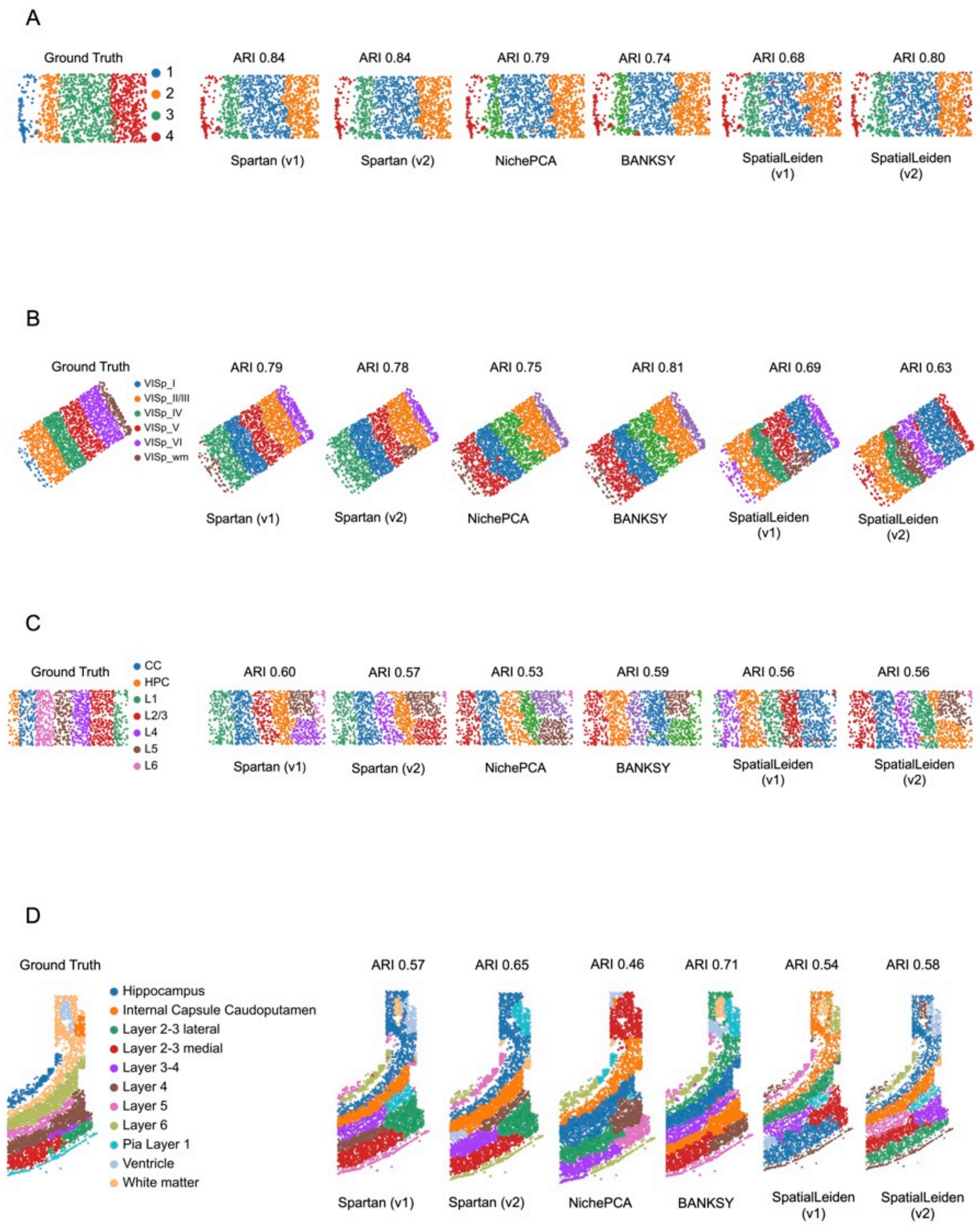


**Supplementary Figure 13.** (A–D) Spatial domain recovery across four representative datasets from the SDM-Bench benchmark collection. For each dataset, the leftmost panel shows ground-truth anatomical annotations, followed by clustering results from Spartan (v1), Spartan (v2), NichePCA, BANKSY, SpatialLeiden (v1), and SpatialLeiden (v2). Adjusted Rand Index (ARI) values reported above each map quantify concordance with ground-truth labels. (A) STARmap sample 417 (BZ5). (B) BaristaSeq Slice 2. (C) STARmap* sample. (D) osmFISH sample.

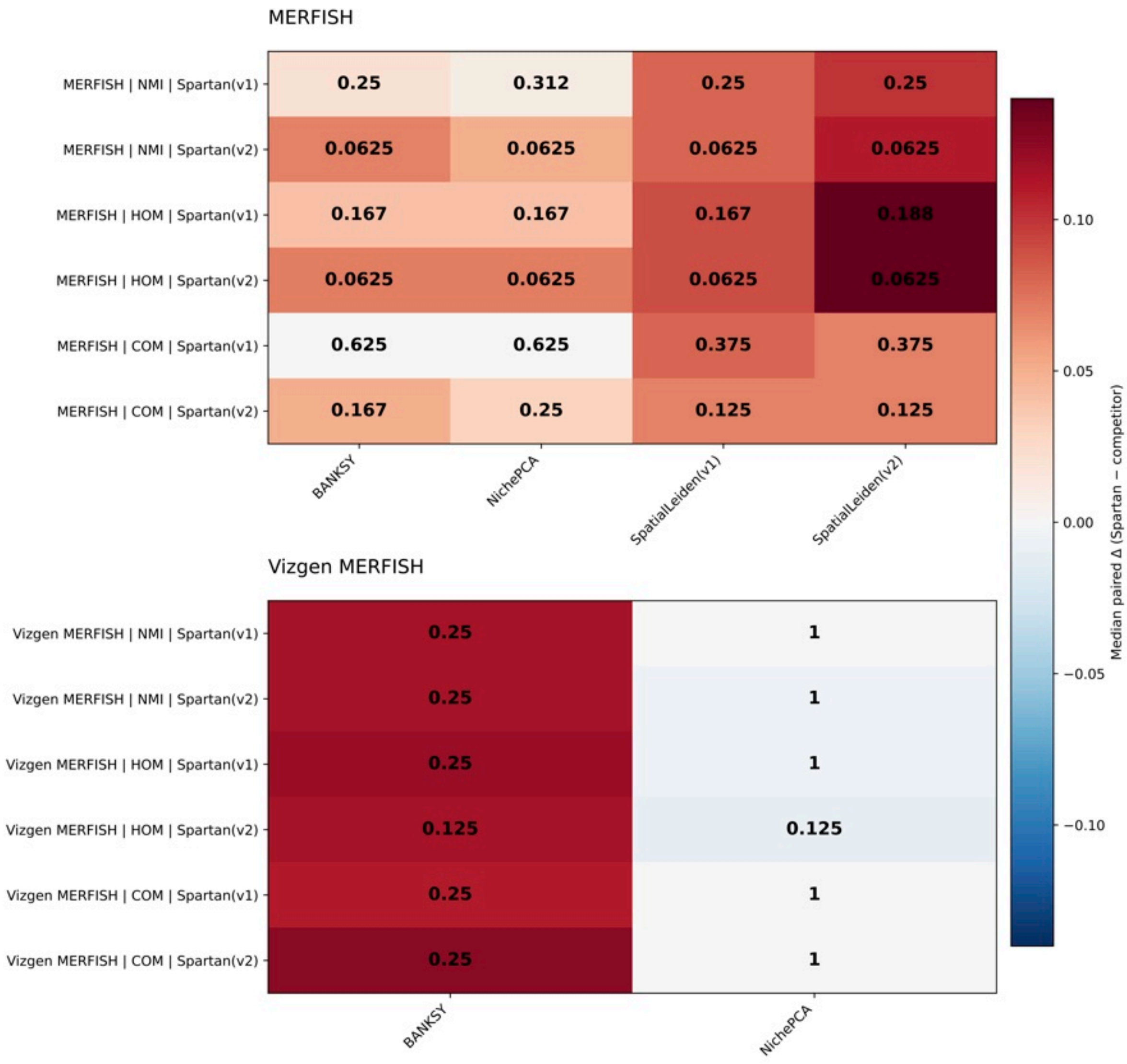


**Supplementary Figure 14.** Paired statistical comparison of Spartan and recently proposed methods. Paired two-sided Wilcoxon signed-rank tests were performed to compare both versions of Spartan (v1 and v2) against NichePCA, BANKSY, and SpatialLeiden (v1 and v2) across samples. Comparisons were conducted on MERFISH and Vizgen MERFISH datasets, with SpatialLeiden included for MERFISH only. Heatmaps show median paired differences (Spartan – comparator) for NMI, HOM, and COM across samples. Cell annotations indicate Benjamini–Hochberg-adjusted P-values (false discovery rate) computed within each dataset and metric across all pairwise comparisons. Only paired samples available for each method comparison were included.

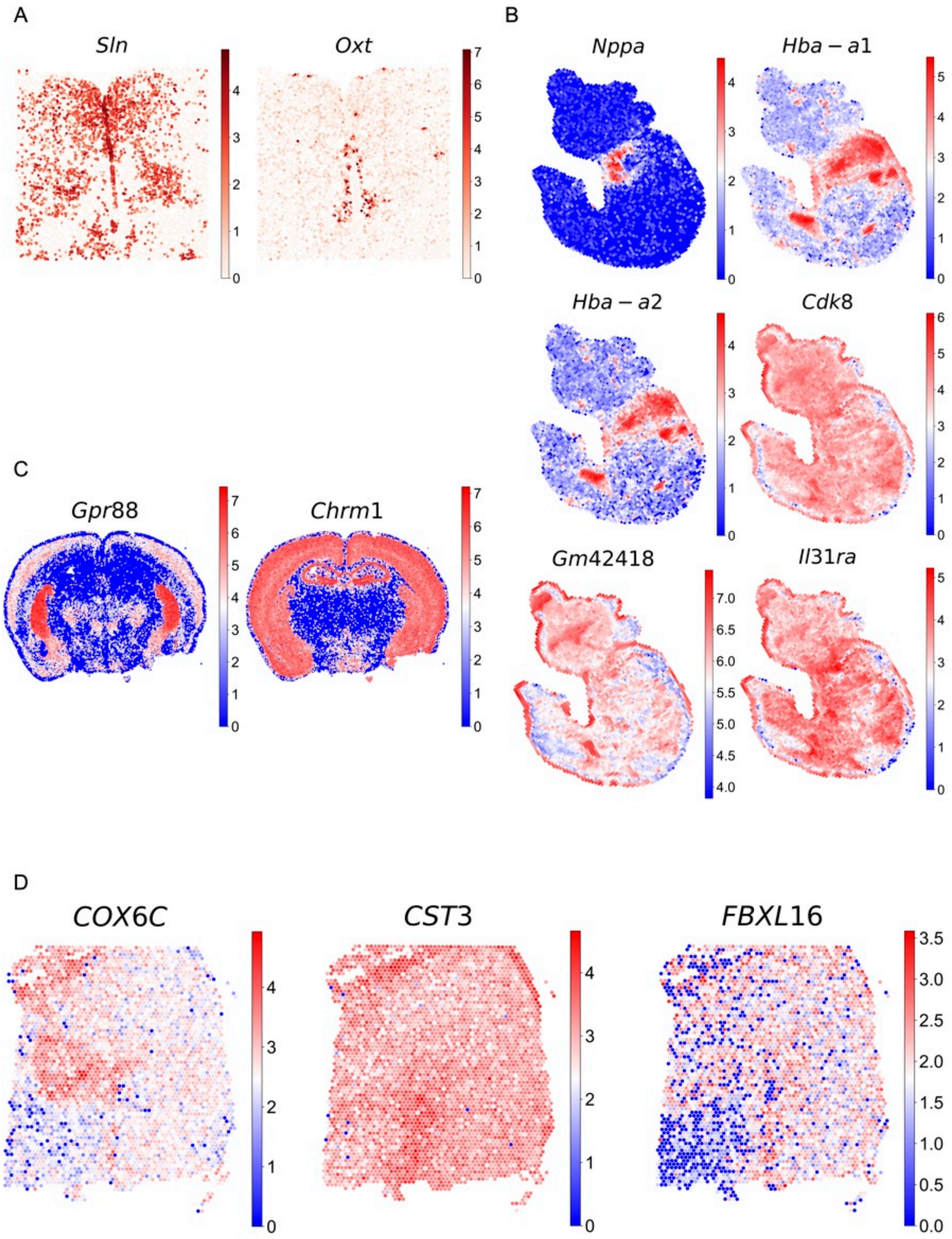


**Supplementary Figure 15.** Additional spatially variable genes (SVGs) corresponding to Fig. 4. (A–D) Spatial expression patterns of additional spatially variable genes that could not be included in the main figure due to space constraints. (A) MERFISH 0.19 sample. (B) Stereo-seq E9.5 E2S2 sample. (C) Vizgen MERFISH S2R3 sample. (D) 10x Visium human DLPFC sample 151673. Gene names are indicated above each spatial map. Color scales represent normalized expression intensity within each sample.

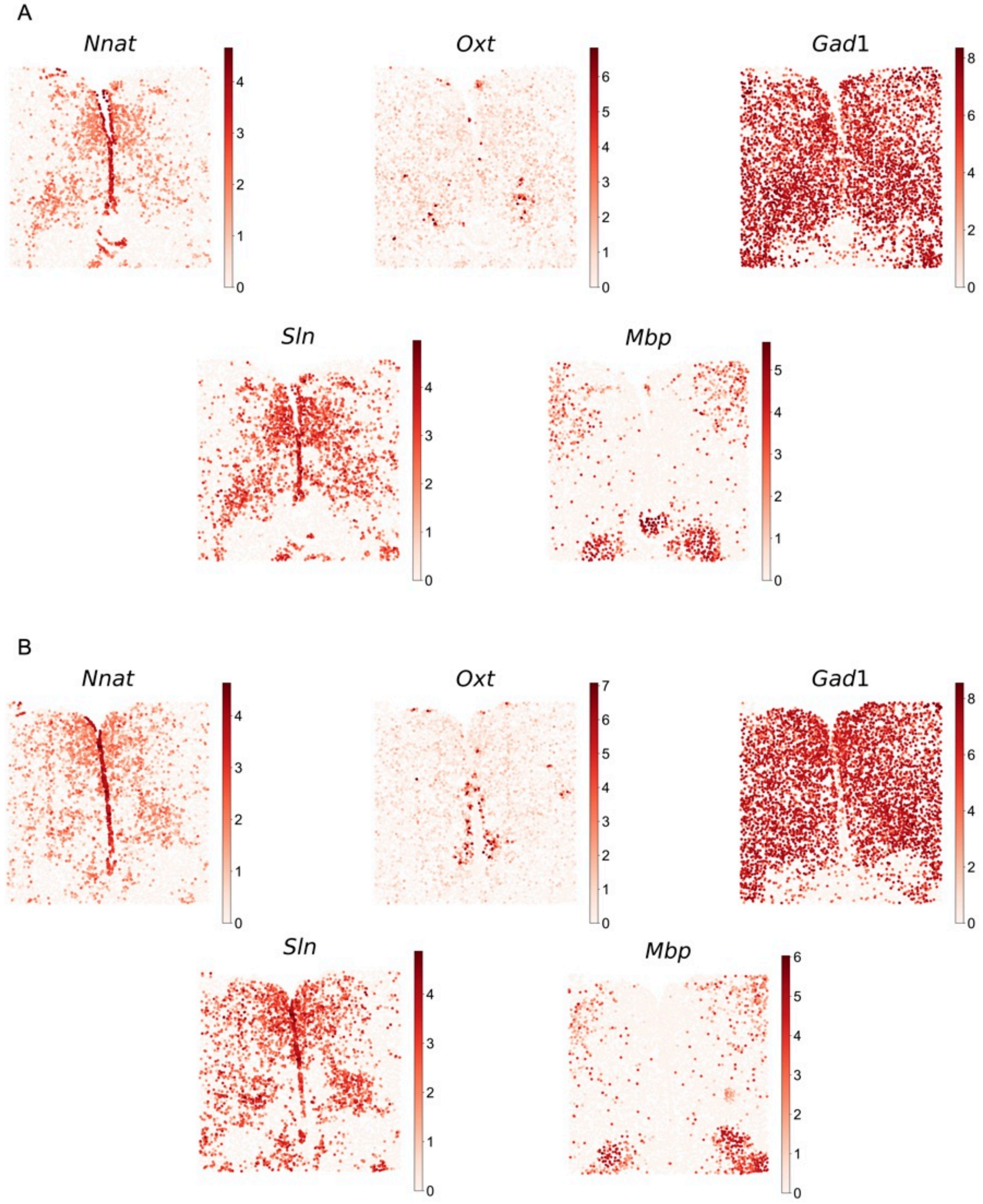


**Supplementary Figure 16.** Cross-sample evaluation of spatially variable genes (SVGs) in MERFISH dataset. (A–B) Spatial expression patterns of selected SVGs across two independent MERFISH samples. (A) Spatial expression in the MERFISH 0.04 sample. (B) Corresponding spatial expression in the MERFISH 0.19 sample. Gene names are indicated above each map. Color scales represent normalized expression intensity within each sample.

#### Supplementary Biological Analysis: Visium HD

The following supplementary figures focus on spatial domain identification and biological interpretation using Visium HD data, complementing the methodological validation and benchmarking results presented above. These analyses include high-resolution domain reconstructions and biological interpretation at cellular resolution.

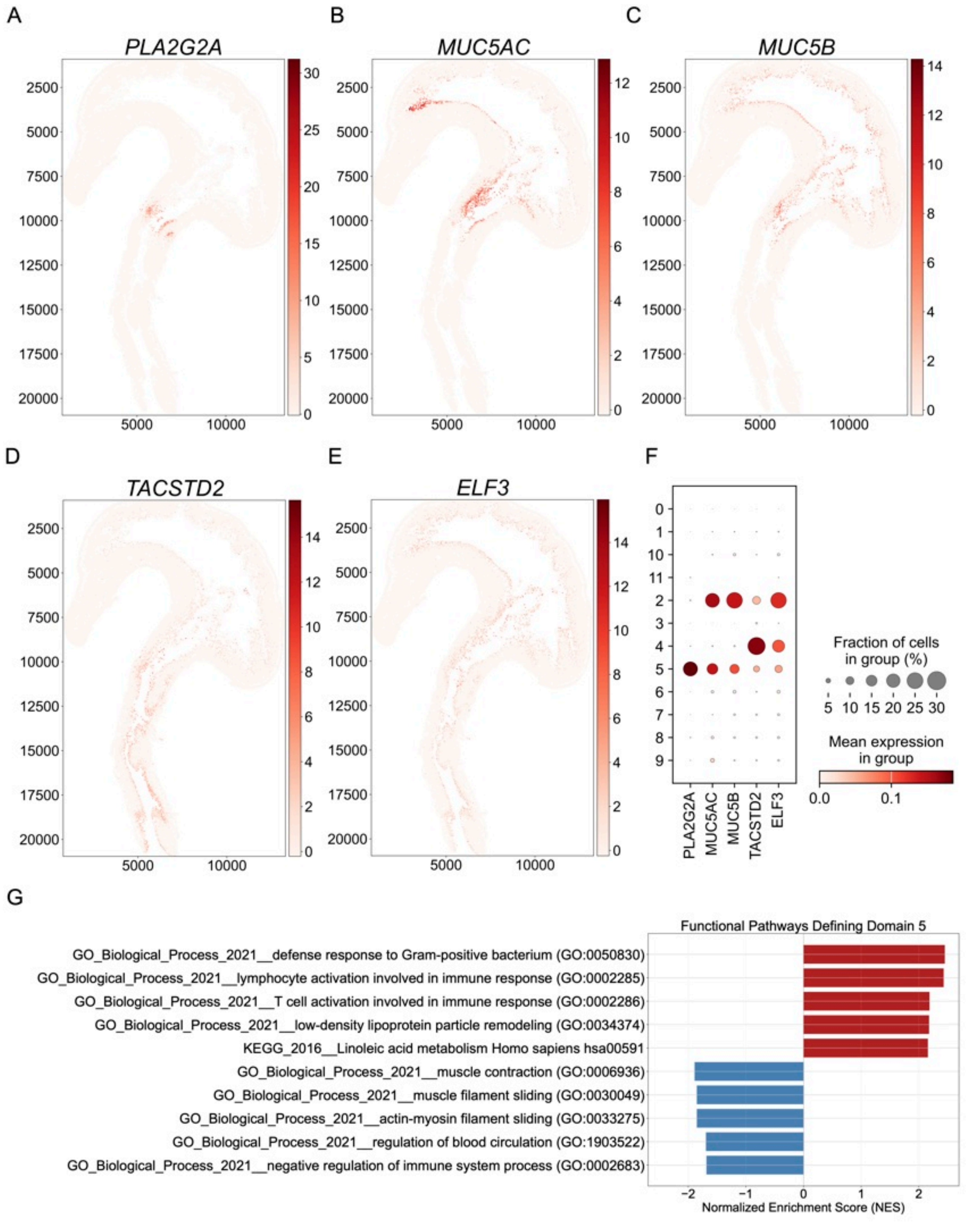


**Supplementary Figure 17.** Domain 5 validation by SVGs, marker enrichment, and functional pathway programs. (A-E) Spatial expression maps of representative Spartan spatially variable genes (SVGs) enriched in domain 0, supporting domain-specific transcriptional coherence. (F) Dotplot summarizing relative expression and detection frequency of selected marker genes across all Spartan domains, highlighting enrichment patterns specific to domain 5. (G) Top enriched functional pathways for domain 5 from preranked gene set enrichment analysis (GSEA), shown as bar plots.

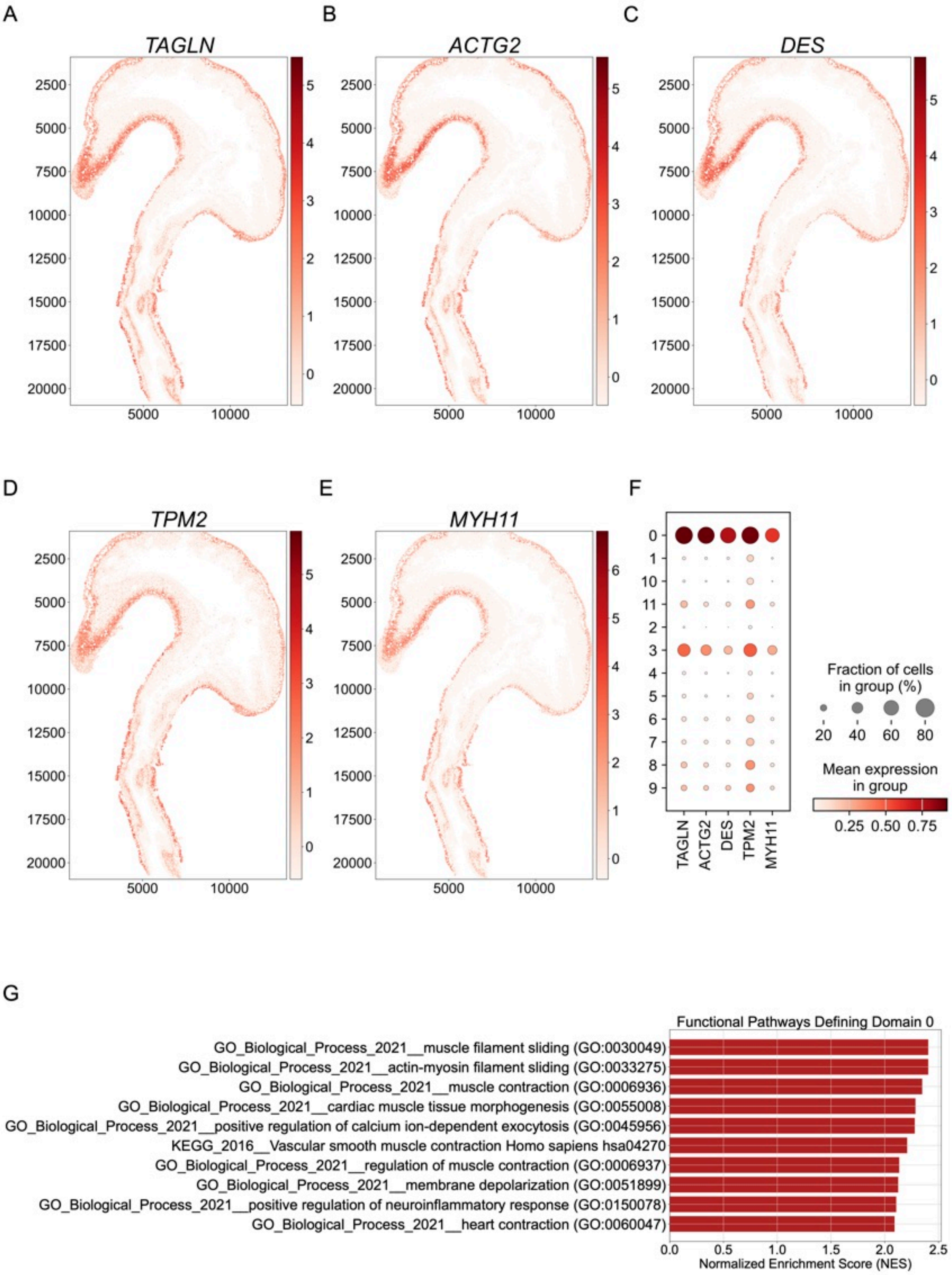


**Supplementary Figure 18.** Domain 0 validation by SVGs, marker enrichment, and functional pathway programs. (A-E) Spatial expression maps of representative Spartan spatially variable genes (SVGs) enriched in domain 0, supporting domain-specific transcriptional coherence. (F) Dotplot summarizing relative expression and detection frequency of selected marker genes across all Spartan domains, highlighting enrichment patterns specific to domain 0. (G) Top enriched functional pathways for domain 0 from preranked gene set enrichment analysis (GSEA), shown as bar plots.

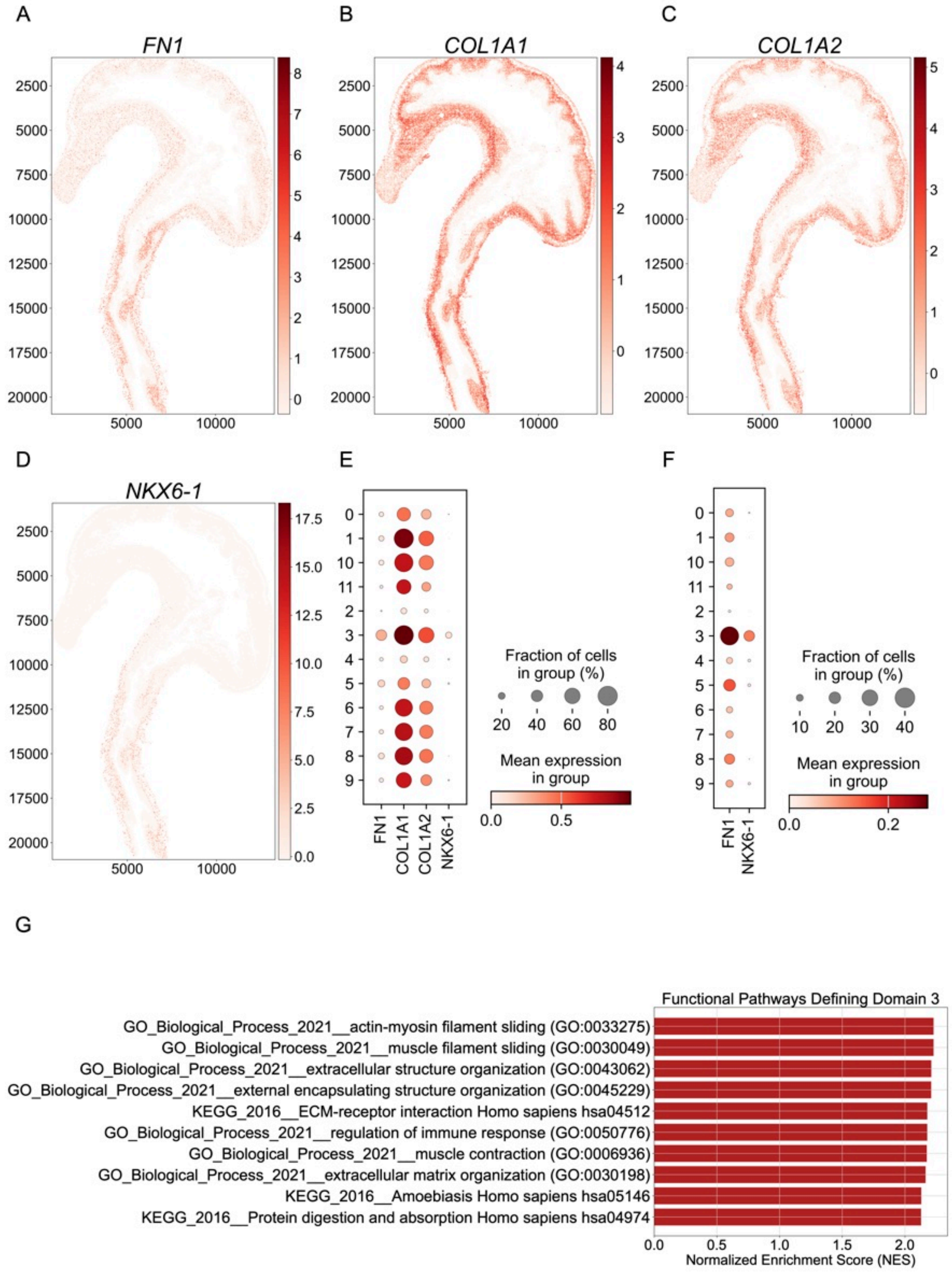


**Supplementary Figure 19.** Domain 3 validation by SVGs, marker enrichment, and functional pathway programs. (A-D) Spatial expression maps of representative Spartan spatially variable genes (SVGs) enriched in domain 3, supporting domain-specific transcriptional coherence. (E-F) Dotplot summarizing relative expression and detection frequency of selected marker genes across all Spartan domains, highlighting enrichment patterns specific to domain 3. (G) Top enriched functional pathways for domain 3 from preranked gene set enrichment analysis (GSEA), shown as bar plots.

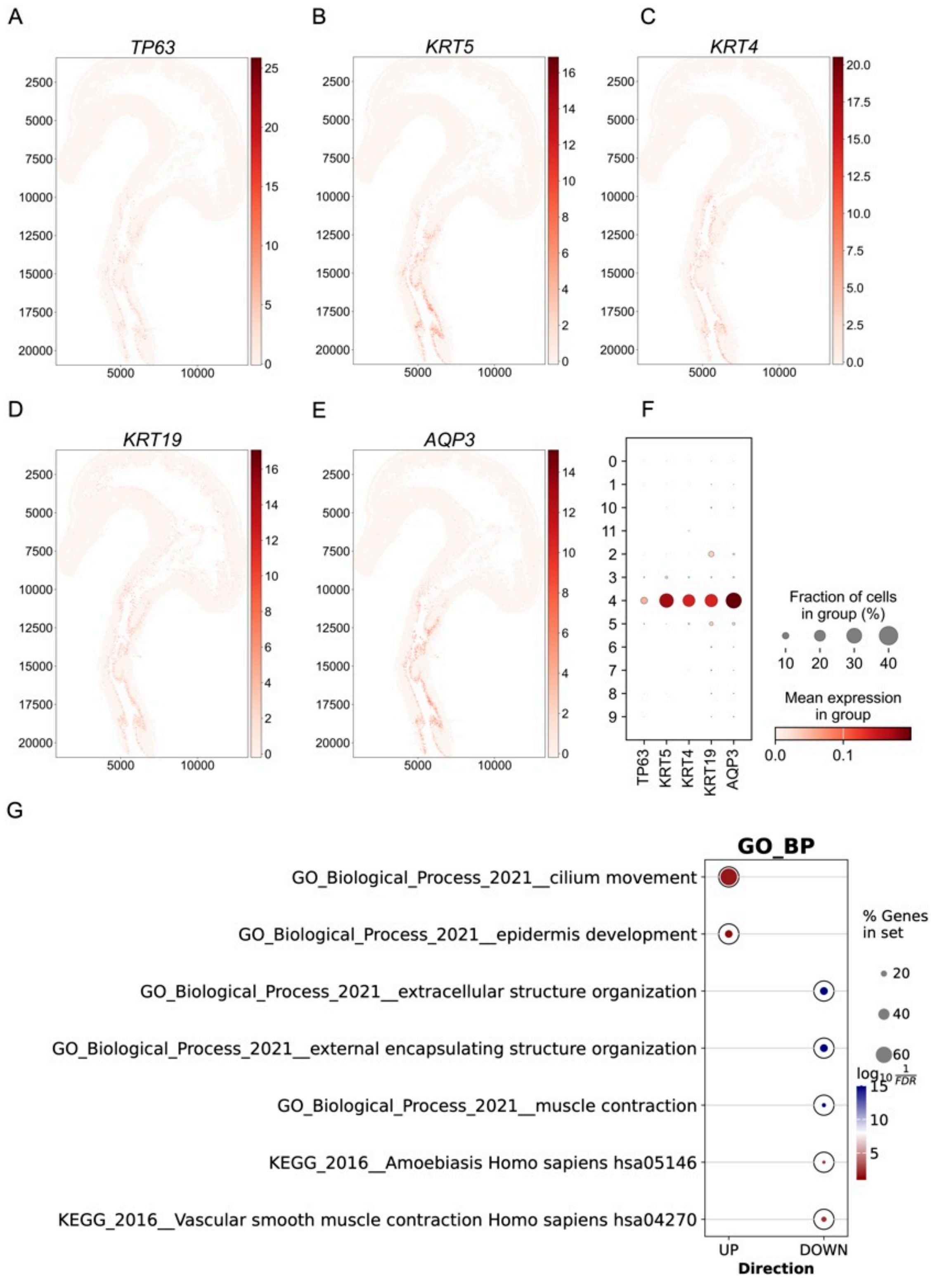


**Supplementary Figure 20.** Domain 4 validation by SVGs, marker enrichment, and functional pathway programs. (A-E) Spatial expression maps of representative Spartan spatially variable genes (SVGs) enriched in domain 4, supporting domain-specific transcriptional coherence. (F) Dotplot summarizing relative expression and detection frequency of selected marker genes across all Spartan domains, highlighting enrichment patterns specific to domain 4. (G) Functional pathway enrichment analysis for domain 4 (pre-ranked GSEA). Bar plots show the top positively enriched pathways (NES > 0) and the top negatively enriched pathways (NES < 0), highlighting programs upregulated versus depleted in domain 4 relative to the reference set.

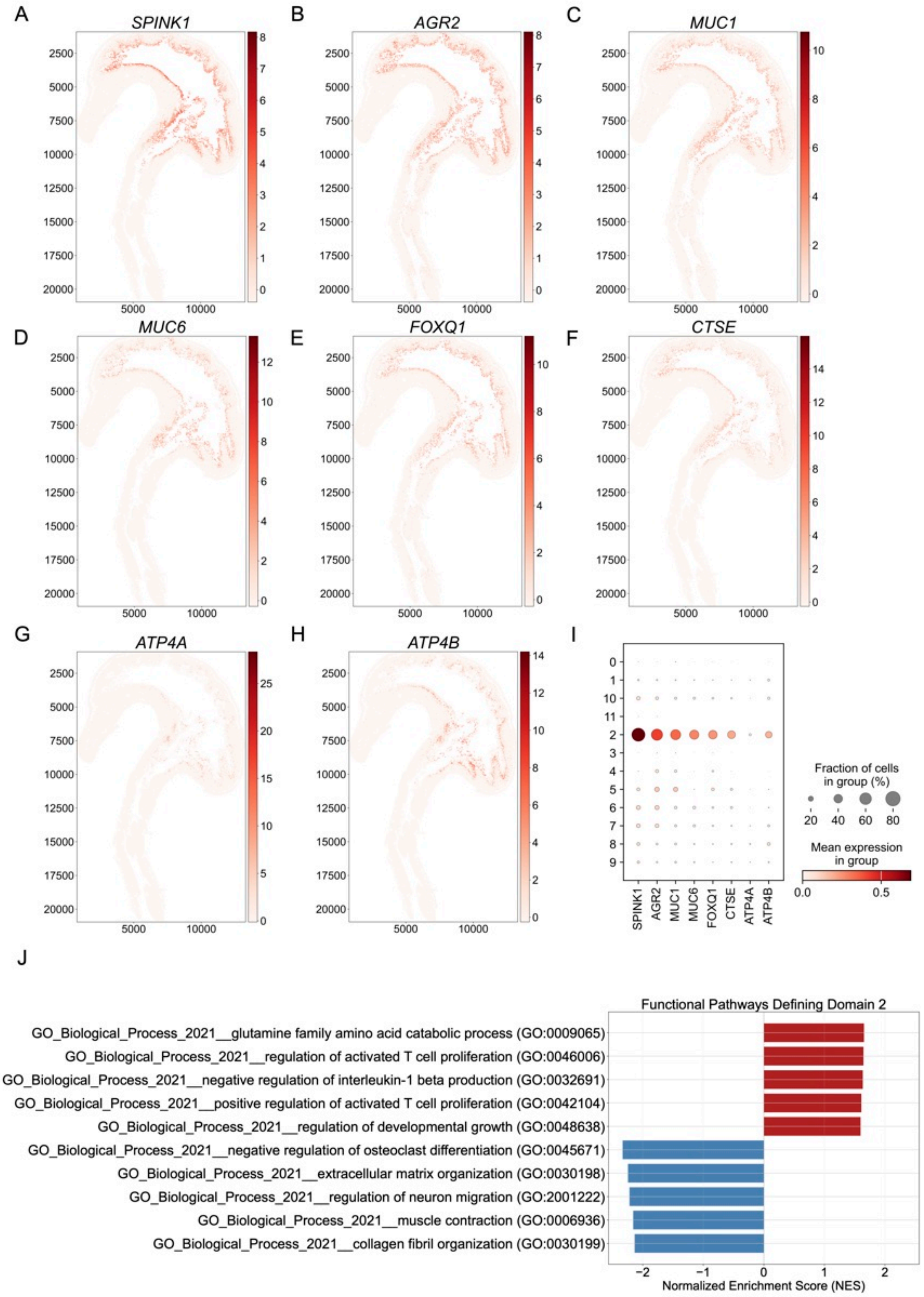


**Supplementary Figure 21.** Domain 2 validation by SVGs, marker enrichment, and functional pathway programs. (A-H) Spatial expression maps of representative Spartan spatially variable genes (SVGs) enriched in domain 2, supporting domain-specific transcriptional coherence. (I) Dotplot summarizing relative expression and detection frequency of selected marker genes across all Spartan domains, highlighting enrichment patterns specific to domain 2. (J) Functional pathway enrichment analysis for domain 2 (pre-ranked GSEA). Bar plots show the top positively enriched pathways (NES $> 0$) and the top negatively enriched pathways (NES $< 0$), highlighting programs upregulated versus depleted in domain 2 relative to the reference set.

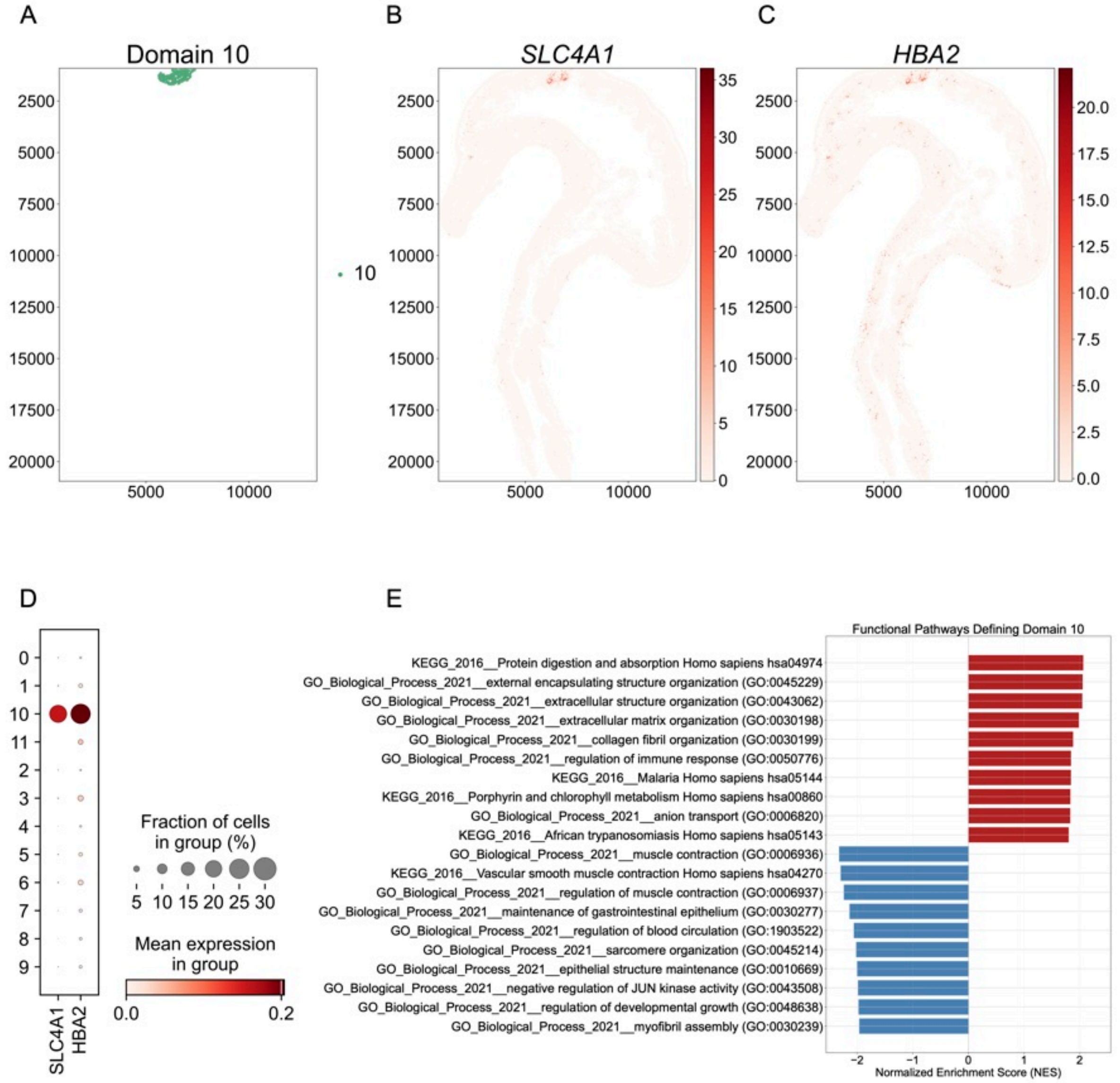


**Supplementary Figure 22.** Domain 10 validation by SVGs, marker enrichment, and functional pathway programs. (A) Spartan spatial domain map for the Visium HD section. Domain 10 is highlighted, showing its spatial localization within the tissue. (B-C) Spatial expression maps of representative Spartan spatially variable genes (SVGs) enriched in domain 10, supporting domain-specific transcriptional coherence. (D) Dotplot summarizing relative expression and detection frequency of selected marker genes across all Spartan domains, highlighting enrichment patterns specific to domain 10. (E) Functional pathway enrichment analysis for domain 10 (pre-ranked GSEA). Bar plots show the top positively enriched pathways (NES > 0) and the top negatively enriched pathways (NES < 0), highlighting programs upregulated versus depleted in domain 10 relative to the reference set.

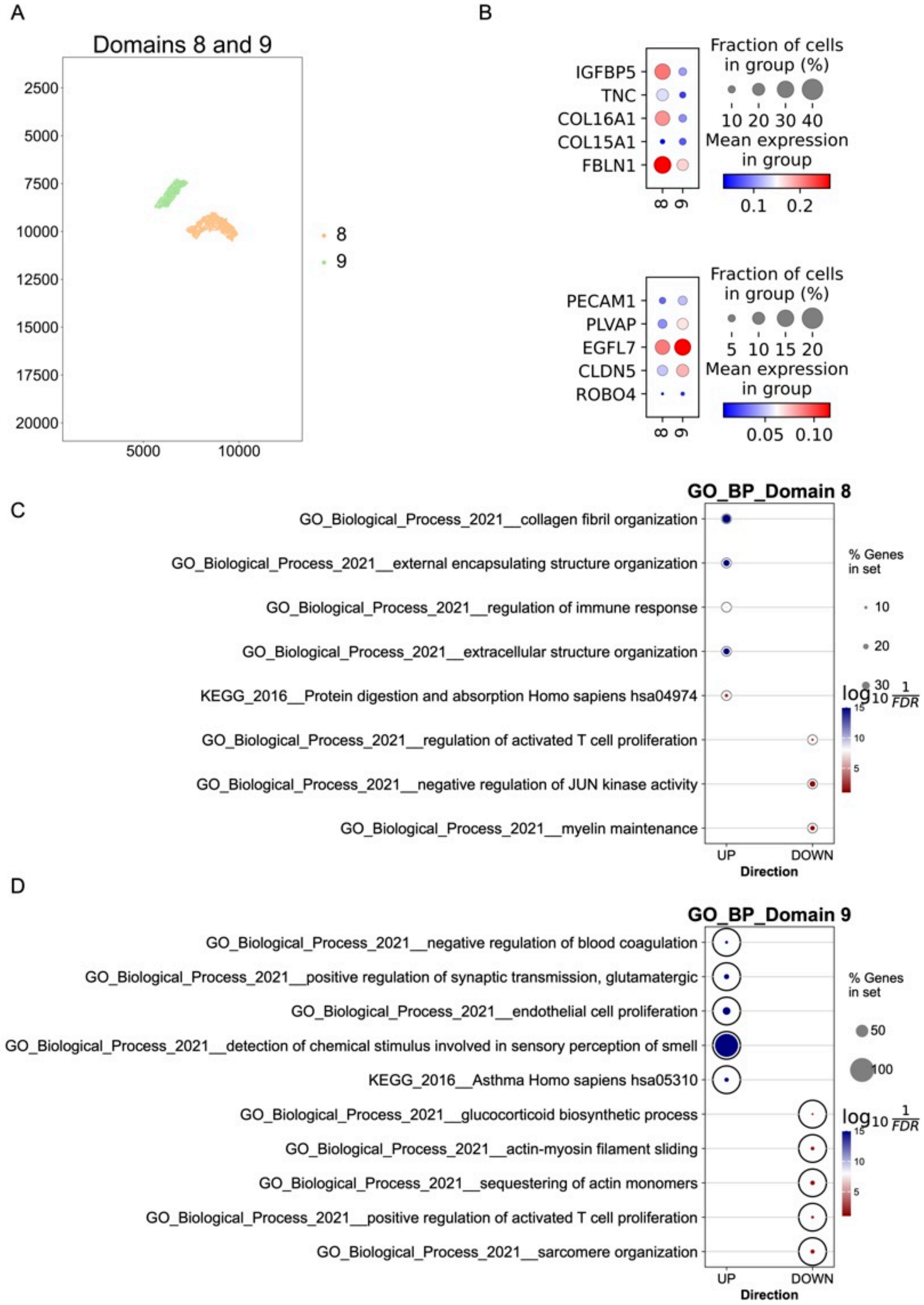


**Supplementary Figure 23.** Opposite-side GEJ-associated stromal programs resolved by Spartan (Domain 8 vs Domain 9). (A) Spatial domain map highlighting Domain 8 (right GEJ-associated stroma connecting into the stomach) and Domain 9 (left GEJ-associated stroma connecting into the stomach), demonstrating that Spartan identifies two spatially separated stromal programs across the junction rather than a single merged compartment. (B) Marker gene dot plots summarizing pairwise DE-derived gene panels for Domain 8 (top) and Domain 9 (bottom), supporting compartment-specific marker enrichment and reinforcing stromal specialization on each side of the junction. (C-D) GO Biological Process (GO_BP) enrichment overview for Domain 8 (C) and Domain 9 (D), showing pathways with positive enrichment (UP; positive NES) and negative enrichment (DOWN; negative NES), illustrating distinct functional polarization between the two GEJ-associated stromal compartments.

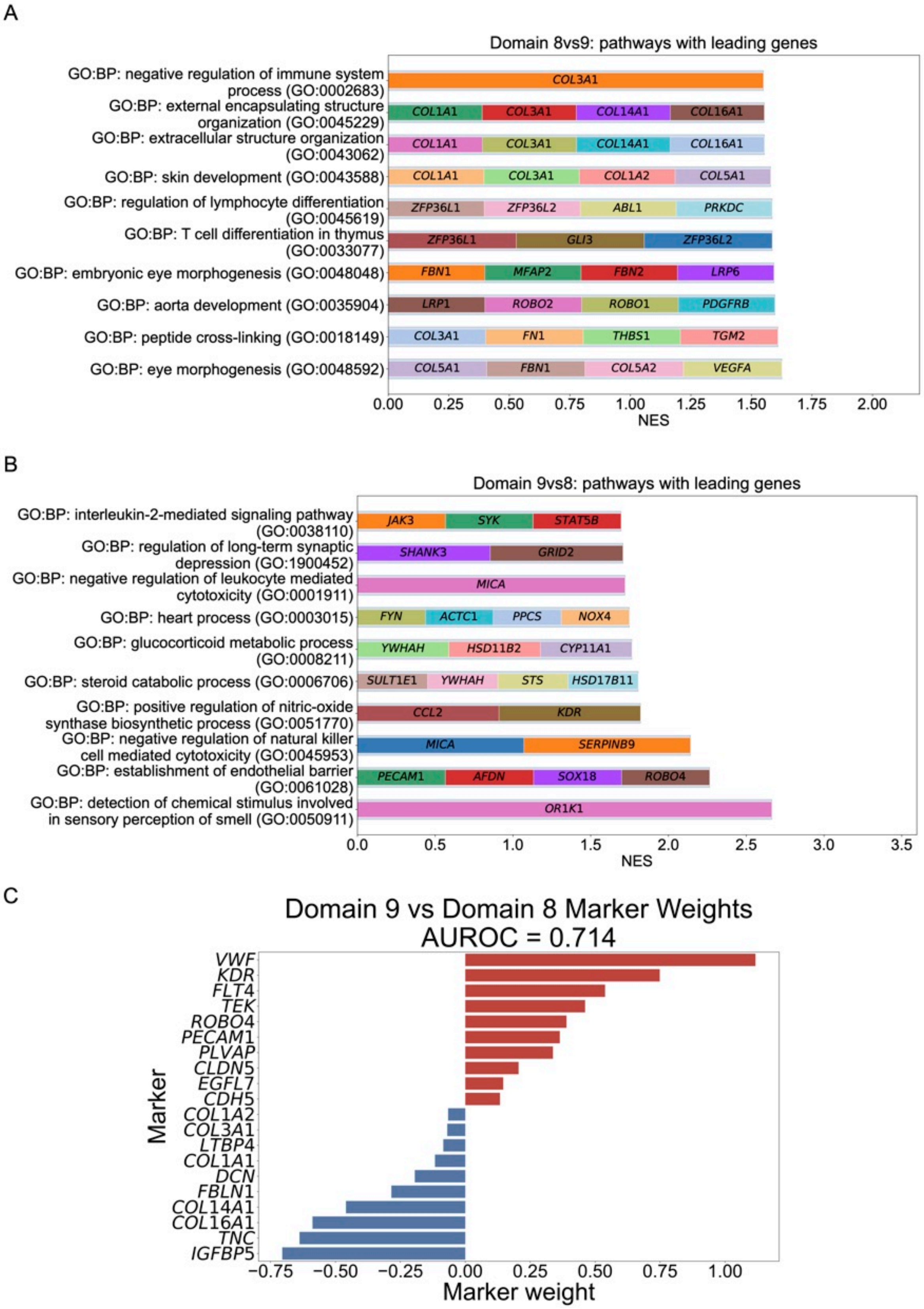


**Supplementary Figure 24.** (A–B) Functional pathway enrichment bar plots with leading-edge gene sets, derived from pairwise differential expression (DE) analysis, shown for Domain 8 (A) and Domain 9 (B). Pathways are ordered by normalized enrichment score (NES) and include both positively enriched and negatively enriched programs, highlighting divergent stromal functional signatures across the GEJ. (C) Weighted logistic regression classification performance and marker contributions for Domain 9 vs Domain 8, reported as AUROC (0.714) with corresponding marker weights, quantifying separability and identifying the most informative genes distinguishing the two GEJ-associated stromal niches.

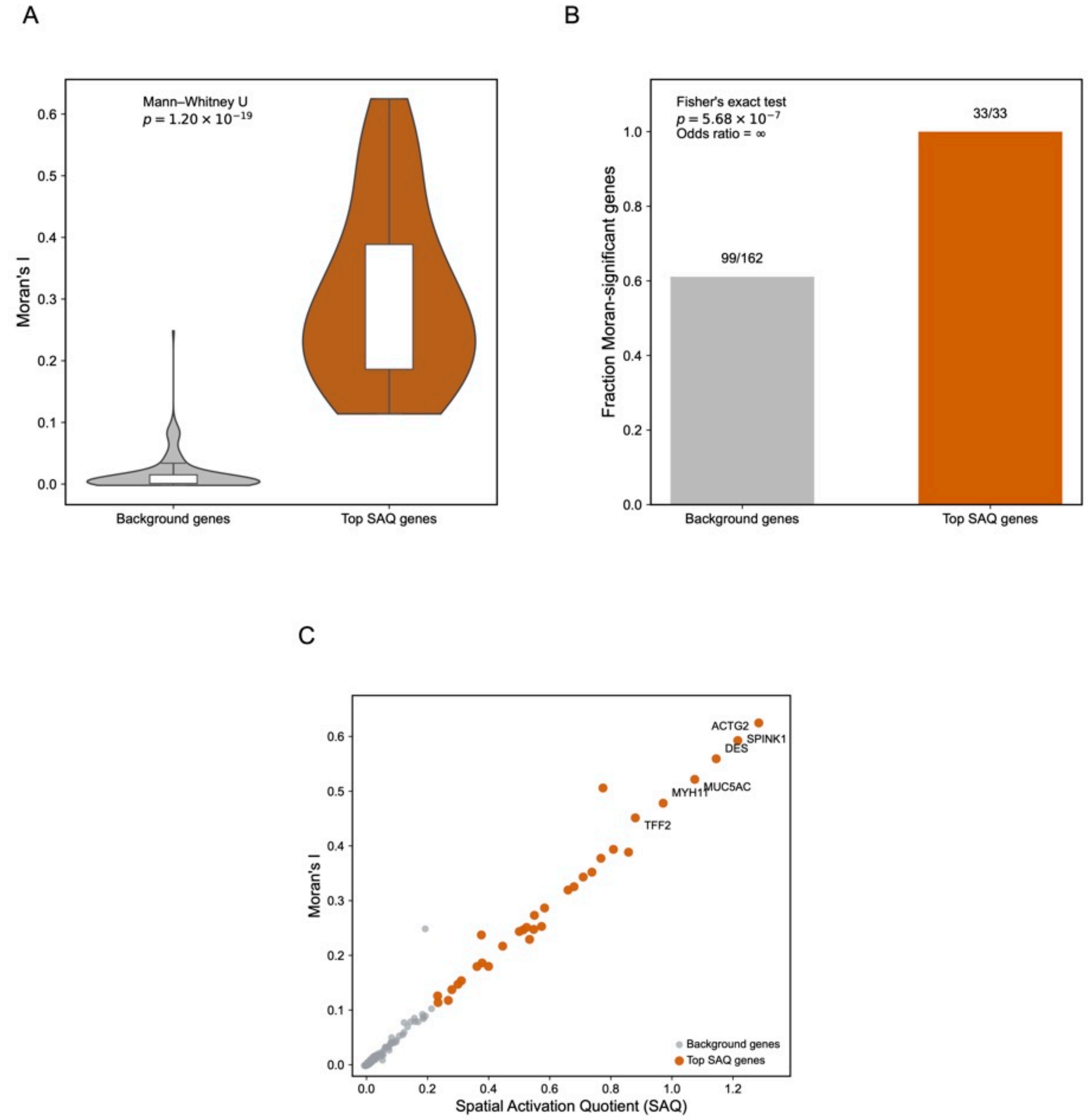


**Supplementary Figure 25.** Validation of SAQ-based spatially variable genes (SVGs) in the Visium HD sample. (A) Distribution of Moran's I values for candidate background genes ($G_{bg}$) and top-ranked SAQ genes ($G_{top}$). Top SAQ genes exhibit substantially higher spatial autocorrelation compared to background genes (Mann–Whitney U test $p = 1.20 \times 10^{-19}$), indicating that SAQ prioritizes spatially structured genes. (B) Fraction of genes identified as significantly spatially autocorrelated (FDR ¡ 0.05) in background and top SAQ gene sets. A higher proportion of Moran-significant genes is observed among top SAQ genes (33/33) compared to background genes (99/162), with significant enrichment (Fisher's exact test $p = 5.68 \times 10^{-7}$, odds ratio $= \infty$). (C) Relationship between SAQ score and Moran's I across candidate genes. Top SAQ-ranked genes (highlighted) are concentrated in the region of high spatial autocorrelation, demonstrating that SAQ ranking aligns with spatially structured gene expression patterns. Together, these results demonstrate that SAQ-based ranking identifies genes with strong and significant spatial autocorrelation, consistent with biologically meaningful spatial variation.

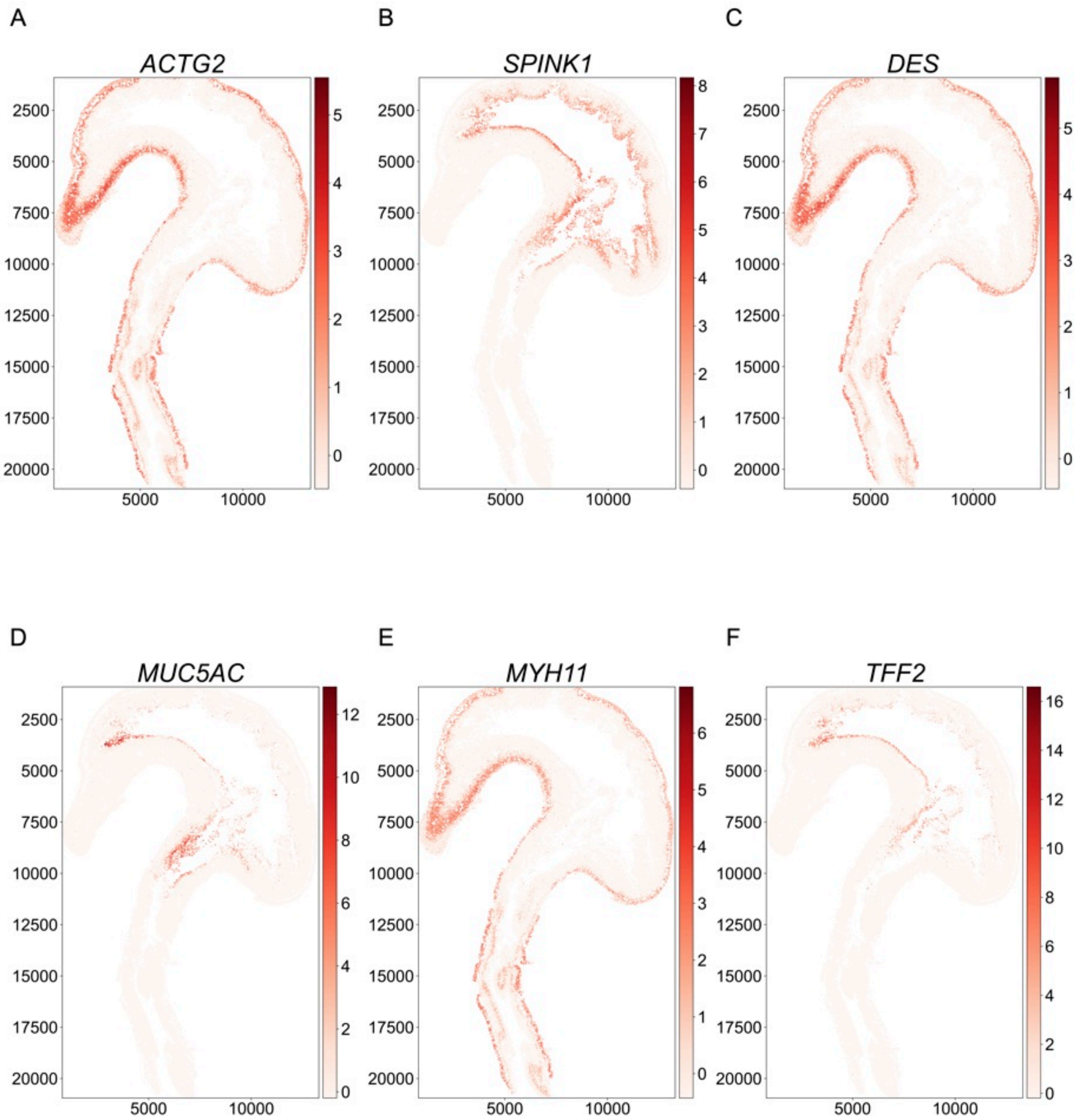


**Supplementary Figure 26.** Spatial expression patterns of top SAQ-ranked genes in the Visium HD sample. Spatial expression maps of the top six genes ranked by SAQ score (*ACTG2*, *SPINK1*, *DES*, *MUC5AC*, *MYH11, andTFF2*). These genes exhibit spatially coherent and domain-restricted expression patterns across the tissue, consistent with strong spatial autocorrelation and localized biological activity. The observed expression patterns illustrate that SAQ prioritizes genes with structured spatial variation, including markers associated with smooth muscle, epithelial, and glandular compartments of the developing esophagus and stomach.

### Supplementary Discussion

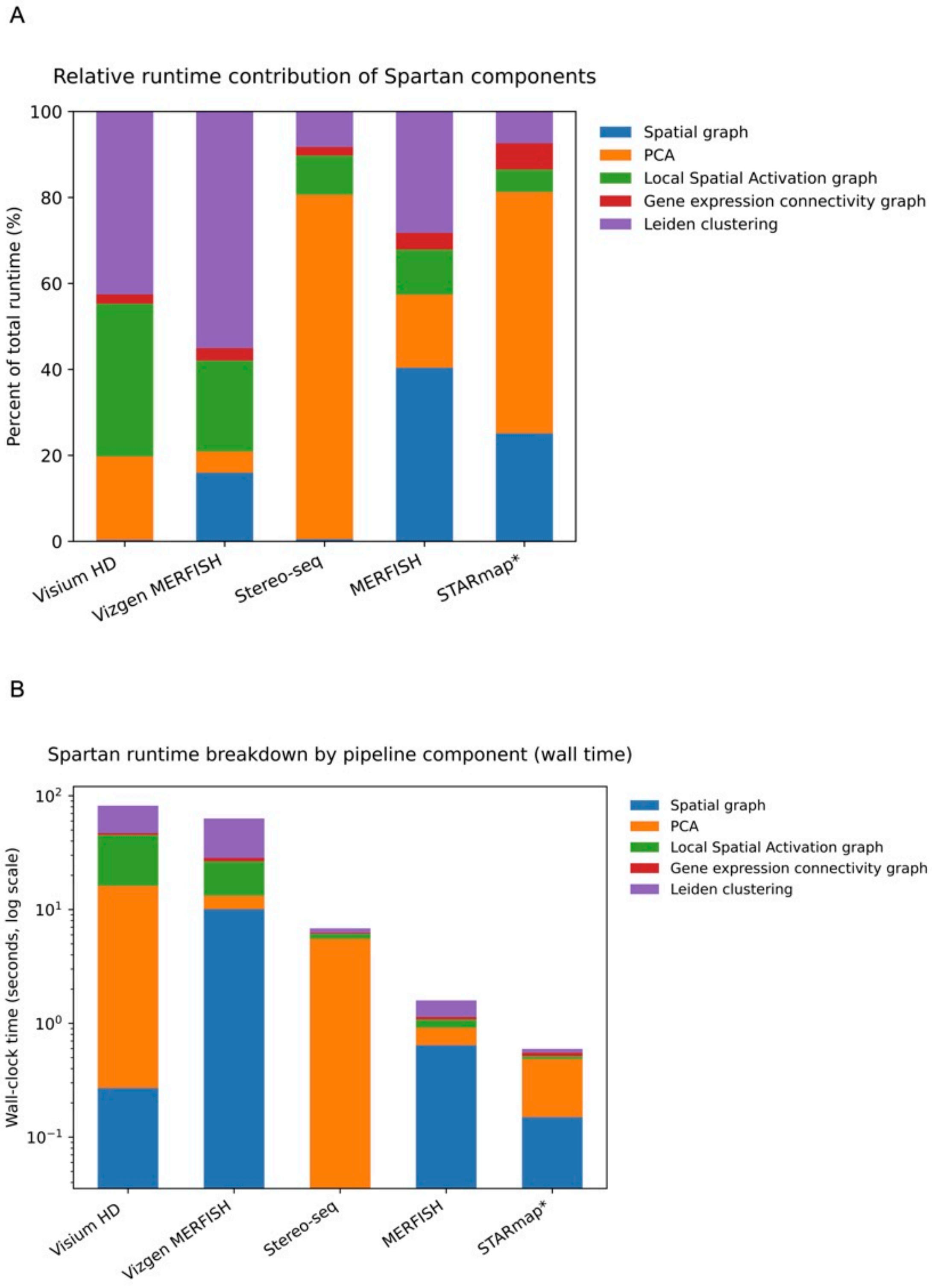


**Supplementary Figure 27.** (A) Relative contribution of individual pipeline components to total runtime across spatial transcriptomics technologies. Bars show the percentage of wall-clock time spent in spatial graph construction, PCA, Local Spatial Activation graph construction, gene expression connectivity graph construction, and multiplex Leiden clustering for a single sequential run. (B) Absolute wall-clock runtime breakdown for the same runs shown in (A), displayed on a logarithmic scale to visualize components with substantially different execution times.

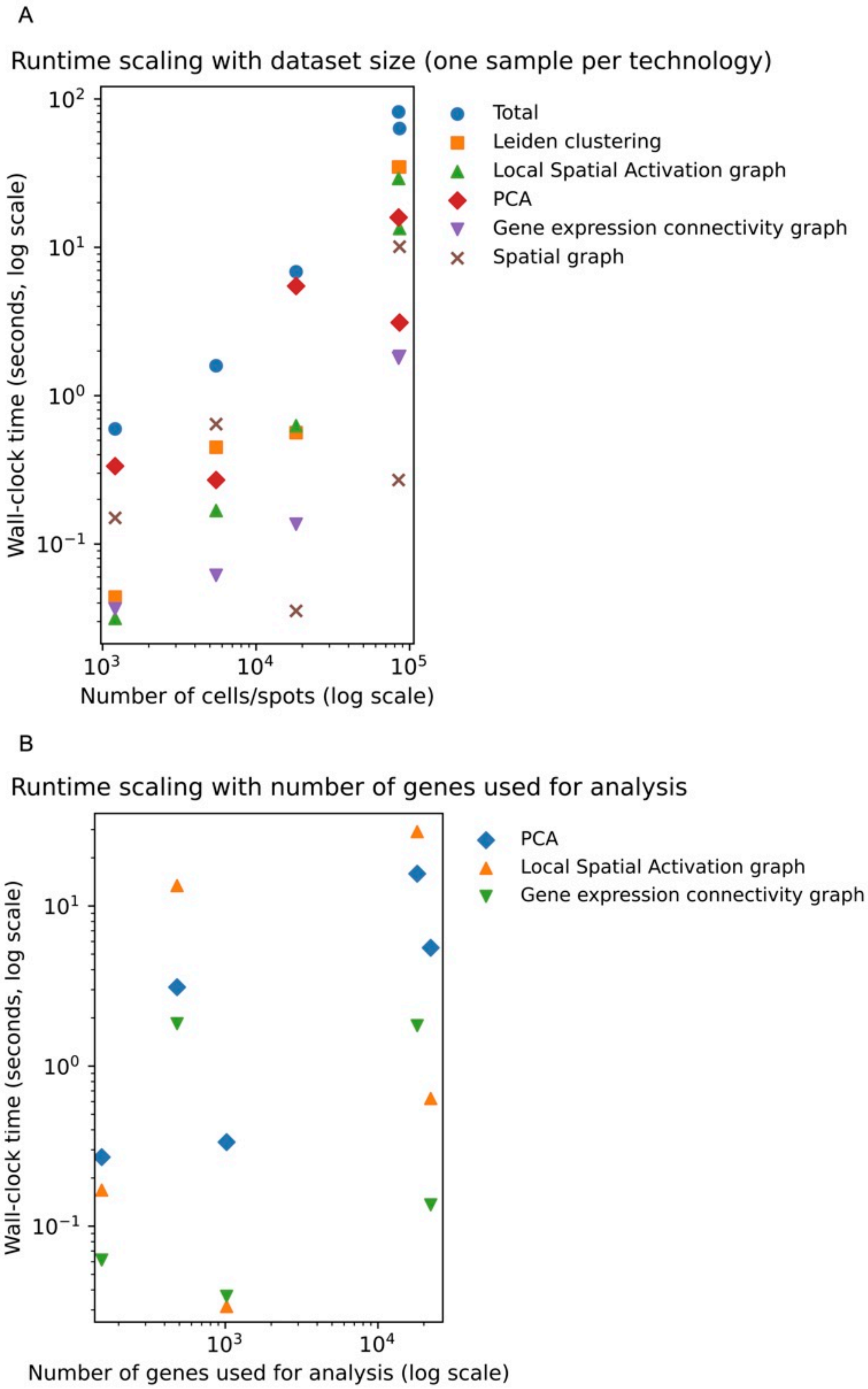


**Supplementary Figure 28.** (A) Runtime scaling with dataset size. Wall-clock time (log scale) is plotted against the number of spatial locations (log scale) for the full pipeline ("Total") and for individual components, using one representative sample per Spatial Transcriptomics technology. (B) Runtime scaling with the number of genes used for analysis. Wall-clock time (log scale) is shown as a function of the number of genes retained after preprocessing and used as input to PCA and graph construction, highlighting the dependence of PCA, Local Spatial Activation, and gene-expression connectivity steps on gene set size. All runtimes were measured as wall-clock time using per-component execution (*%%time*) and correspond to a single execution of the full Spartan pipeline with fixed hyperparameters and resolution.

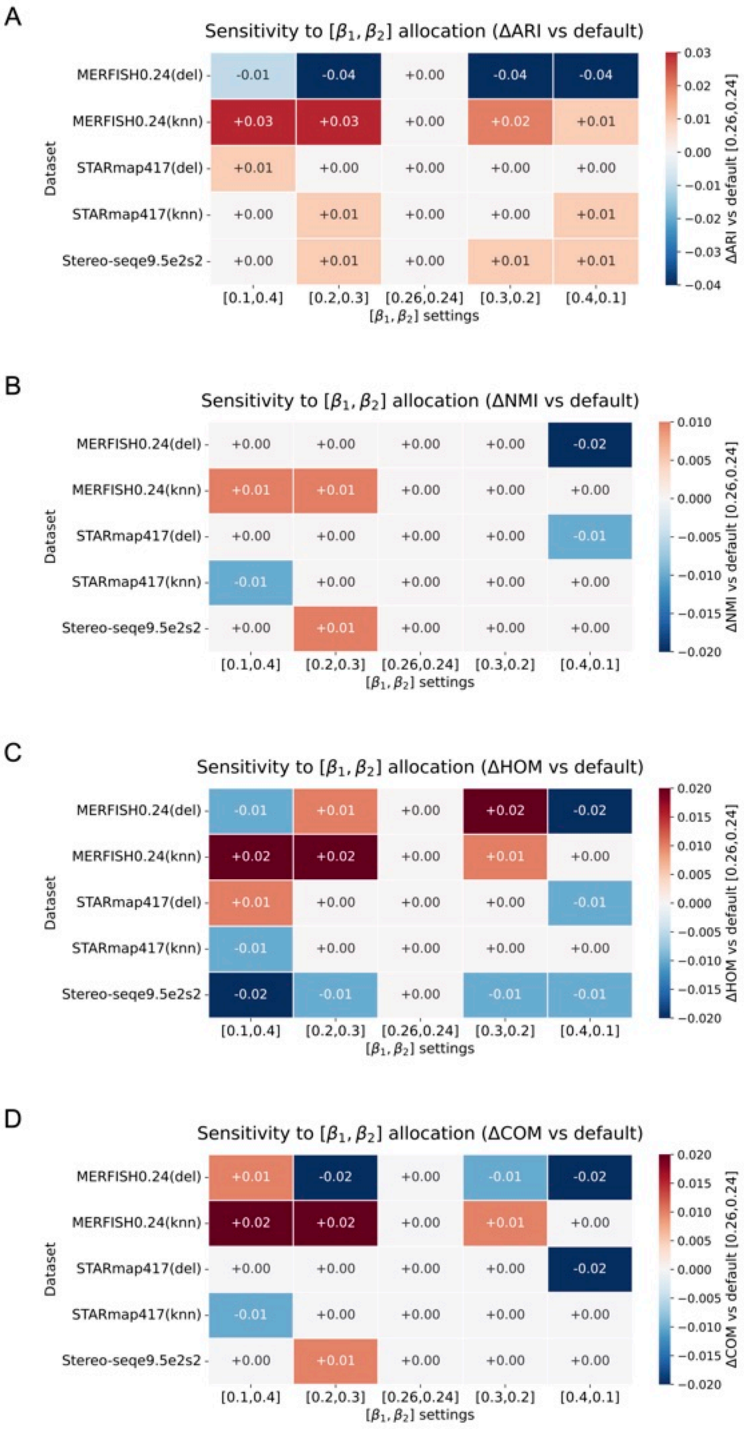


**Supplementary Figure 29.** Sensitivity of Spartan to $\beta_1$ and $\beta_2$ parameter allocation. (A–D) Sensitivity of Spartan's clustering performance to the allocation of weights between graph layers, evaluated across multiple spatial transcriptomics datasets. Heatmaps report changes in performance relative to the default parameter configuration ($\beta_1 = 0.26$, $\beta_2 = 0.24$), measured as ΔARI (A), ΔNMI (B), ΔHOM (C), and ΔCOM (D). Here, $(1 - \beta_1)$ controls the weight of the *Local Spatial Activation graph*, while $(1 - \beta_2)$ controls the weight of the *spatial graph*; the remaining weight is assigned to the gene-expression connectivity graph. Rows correspond to datasets (including alternative neighborhood constructions where applicable), and columns indicate alternative $(\beta_1, \beta_2)$ settings. Values denote absolute differences relative to the default configuration, with color encoding the direction and magnitude of change. Across datasets and metrics, performance varies smoothly across parameter settings, indicating that Spartan is robust to moderate changes in $\beta_1$ and $\beta_2$ and does not require fine-grained parameter tuning.

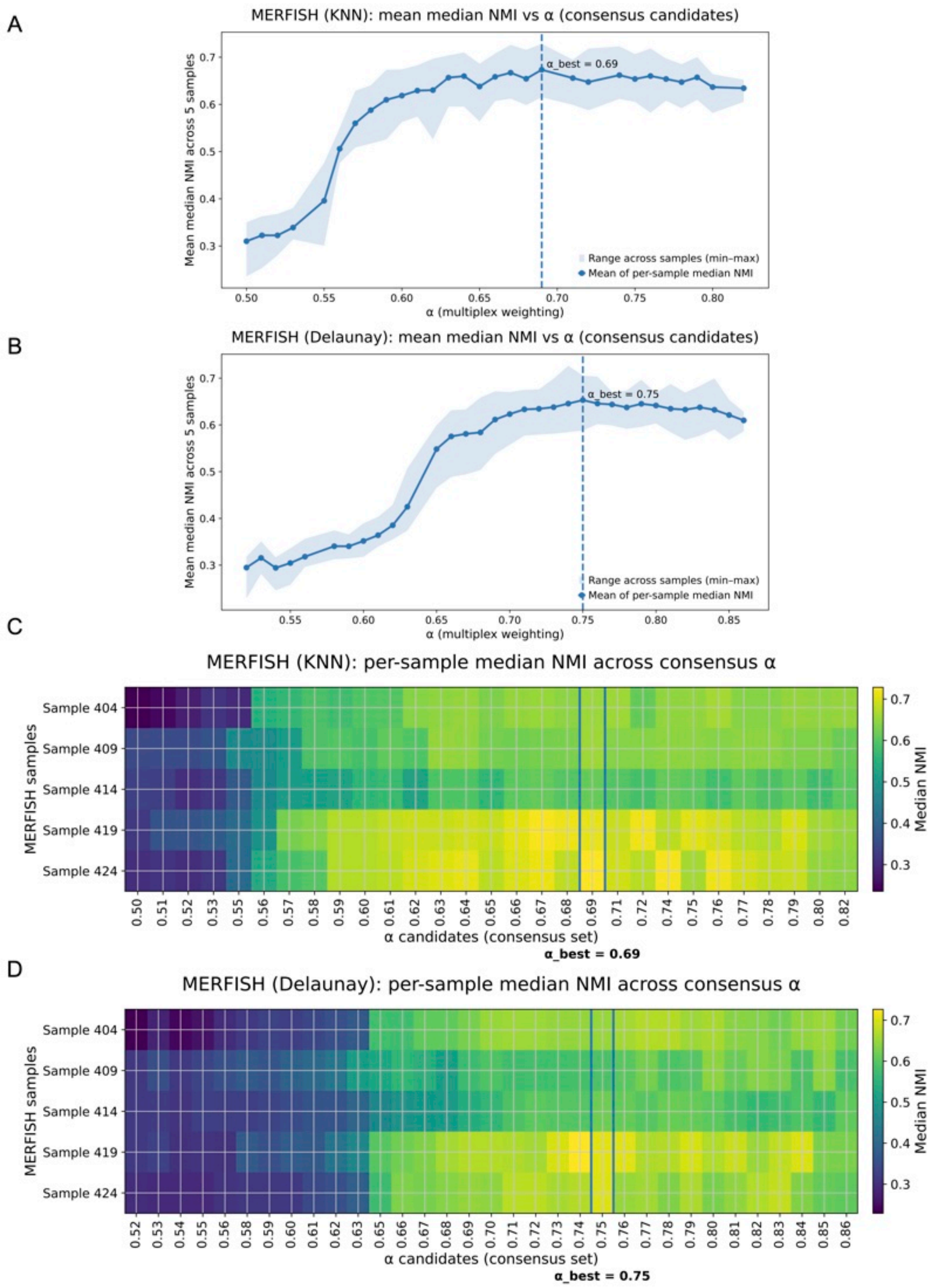


**Supplementary Figure 30.** Dataset-level $\alpha$ selection for the MERFISH dataset across spatial graph constructions. (A) MERFISH (KNN spatial graph): mean median normalized mutual information (NMI) across five samples plotted as a function of candidate $\alpha$ values retained in the consensus set. The solid line denotes the mean of per-sample median NMI values, and the shaded region indicates the range across samples (minimum–maximum). The vertical dashed line marks the selected dataset-level $\alpha_{\text{best}}$. (B) MERFISH (Delaunay spatial graph): mean median NMI across samples plotted against the same consensus $\alpha$ candidates, shown using the same conventions as in (A). (C) MERFISH (KNN spatial graph): heatmap of per-sample median NMI values across the consensus $\alpha$ candidates. Rows correspond to individual MERFISH samples and columns to candidate $\alpha$ values, illustrating cross-sample consistency at the selected $\alpha_{\text{best}}$. (D) MERFISH (Delaunay spatial graph): per-sample median NMI values across the consensus $\alpha$ candidates, shown analogously to (C). Together, these panels demonstrate that Spartan selects a stable, dataset-level $\alpha$ that yields consistently strong performance across samples and remains robust to the choice of spatial graph construction.

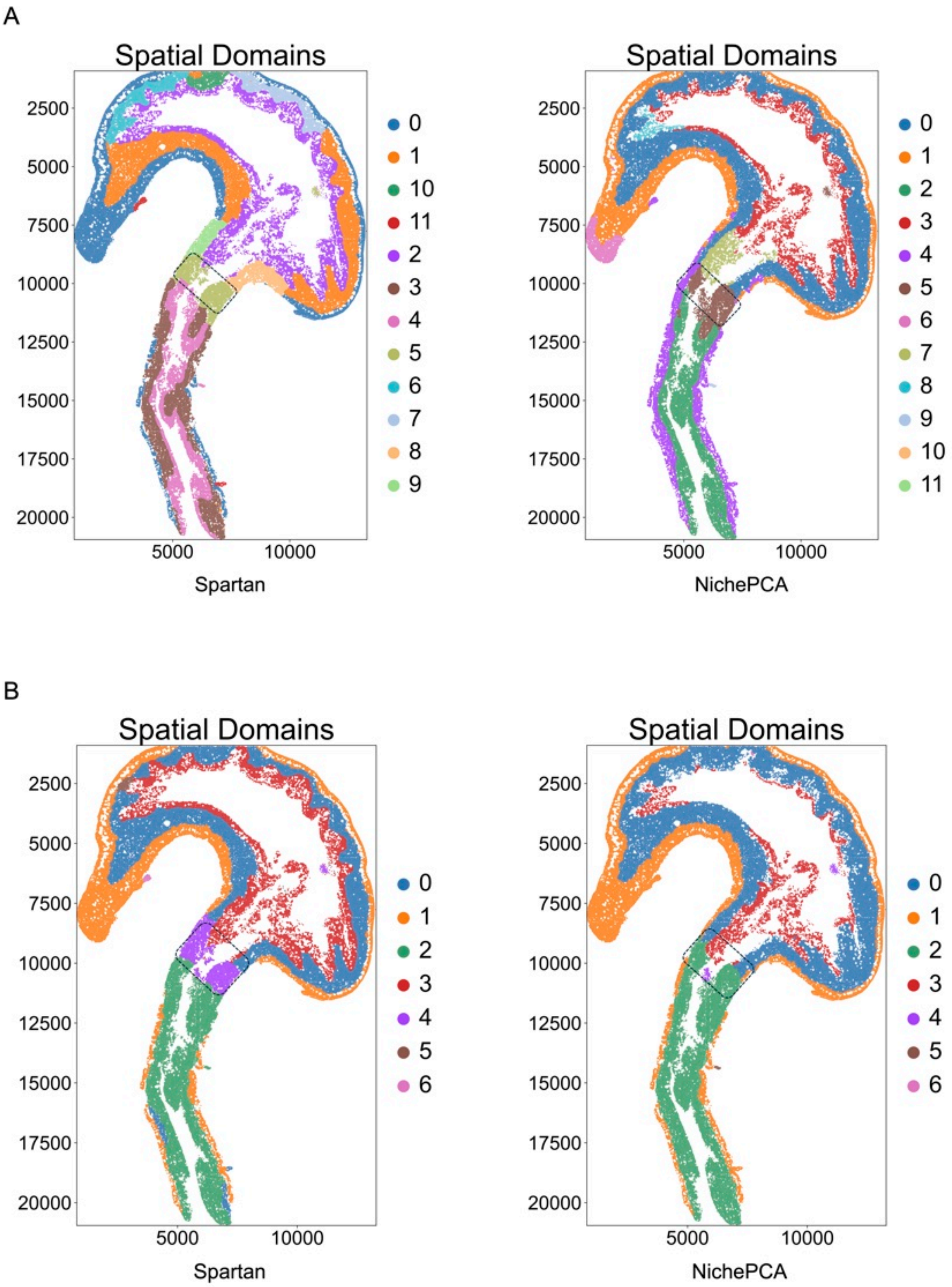


**Supplementary Figure 31.** (A–B) Spatial domain maps for Spartan and NichePCA at baseline ($R = 12$) (A) and low clustering granularity ($R = 7$) (B). For each method, hyperparameters were fixed to the main setting, and only the Leiden resolution was varied to obtain the target cluster count. The dashed box indicates the gastroesophageal junction (GEJ) region used as a visual reference across clustering scales.

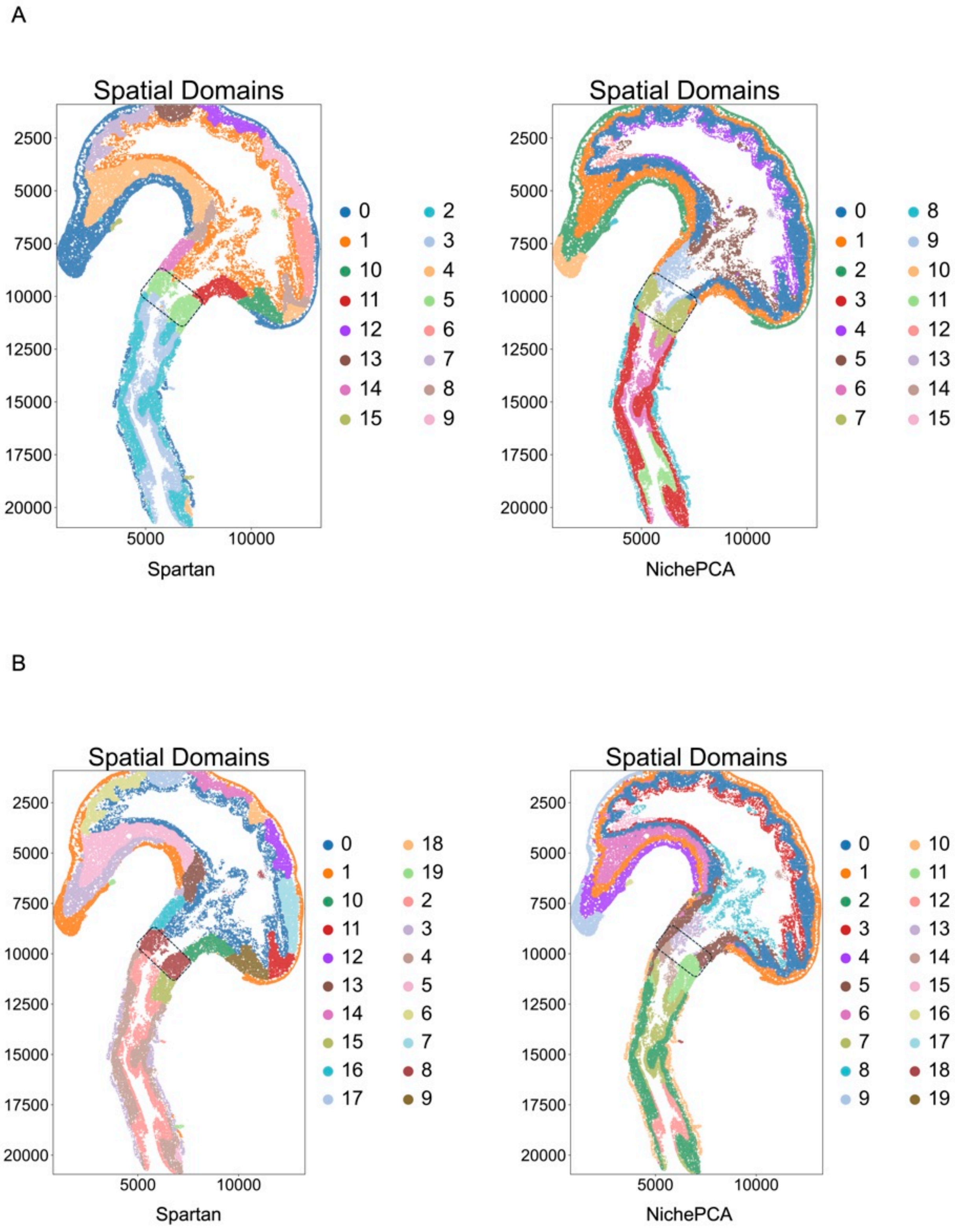


**Supplementary Figure 32.** (A–B) Spatial domain maps for Spartan and NichePCA at increased clustering granularity, $R = 16$ (A) and $R = 20$ (B). Method-specific hyperparameters were fixed, and only the Leiden resolution was varied. The dashed box marks the gastroesophageal junction (GEJ), illustrating the stability of the transition region under increasing cluster resolution.
